# A universal single-cell transcriptomics atlas of human lung decodes multiple pulmonary diseases

**DOI:** 10.1101/2024.12.17.628654

**Authors:** Fanjie Wu, Wenhao Cai, Hai Tang, Shikang Zheng, Haiyue Zhang, Yixin Chen, Yutong Han, Dingli Zhou, Ruihan Wang, Mingli Ye, Renke You, Amin Chen, Jiaqi Li, Xuegong Zhang, Weizhong Li

## Abstract

Human lung is a complex organ susceptible to various diseases. Single-cell transcriptomic studies provide rich data to targeting specific research questions. Here, we present uniLUNG, the largest lung transcriptomic cell atlas, comprising over 10 million cells across 20 disease states and healthy controls. We ensembled a universal hierarchical annotation framework and conducted a full benchmarking of data integration to define a standardized nomenclature and marker genes for lung cell types. Using uniLUNG, we identified Lym-monocyte and T-like B cells, new cell types in specific lung diseases, confirming their existence by comparing with external single-cell atlases. Additionally, we discovered the NSCLC-like SCLC subpopulation, a transitional malignant cell population associated with the transition from NSCLC to SCLC, which was validated and further characterized in spatial dimensions, revealing its complex role in tumour progression. Overall, uniLUNG represents a comprehensive range of human lung cell diversity, providing valuable data resources and a reliable foundation for lung single-cell research.

**HIGHLIGHTS:**

1. The largest scRNA atlas for human lung covers 10 million cells from 20 lung states.
2. A four-level universal cell annotation framework encompasses 120 lung cell types.
3. Comprehensive benchmarking on 18 strategies guides data integration.
4. Specific distribution of Lym-monocytes and T-like B cells in specific lung diseases.
5. The NSCLC-like SCLC subpopulation in transitional events of malignant cells from NSCLC to SCLC.

## INTRODUCTION

The lung is a complex organ susceptible to various diseases. Single-cell transcriptomics has revolutionized our ability to uncover lung cell heterogeneity and explore cellular relationships with both normal and diseased states. Over recent years, large datasets generated from single-cell studies have spurred the development of comprehensive reference atlases ^1, 2^ of human cells, significantly advancing our understanding of health and disease. Several efforts have focused on creating cellular atlases of healthy human lungs, revealing novel biological insights ^3–5^. However, limitations in sample collection, experimental biases, and disease-specific focuses ^6, 7^ have restricted the diversity of these datasets, impeding a comprehensive analysis of lung cell phenotypes across different conditions. Integrating datasets from multiple studies is essential to overcome these limitations.

The current trend in lung research is the development of integrative cellular atlases, combining diverse datasets to uncover rare cell types ^8–10^, characterize disease-associated cellular heterogeneity ^11, 12^, and develop new classification strategies ^13, 14^. Despite these advances, constraints on donor, dataset, sample and cell types diversity ^9, 10, 15–18^, as well as metadata completeness ^19, 20^ mean these atlases still fall short of capturing the full cellular diversity of the lung. A complete cellular atlas should encompass comprehensive cellular, phenotypic, physiological, and multi-omics data across all cell types and disease states. The Human Lung Cell Atlas (HLCA) ^21^ marked a significant milestone by integrating healthy and diseased lung data to create a reference for lung biology. However, it remains room to expand its coverage of lung diseases and previously generated data. Therefore, extensive utilization of existing datasets can enhance the representativeness of integrative cellular atlases and further emphasize and consolidate their role as reference resources by improving their accessibility.

In this study, we introduce uniLUNG (https://lung.unifiedcellatlas.org), the first 10-million-scale cells reference atlas for lung health and disease. Representing the most extensive and accessible human lung single-cell atlas to date, uniLUNG integrates diverse datasets derived from individuals of various ethnicities, ages, and sexes, including cases of major lung diseases. The atlas employs a four-level hierarchical universal annotation framework (uHAF) to classify 120 known lung cell types, offering a detailed representation of lung biology under both healthy and diseased states. Additionally, we conducted a rigorous benchmarking of data integration methods on million-scale cells datasets to guide the construction of uniLUNG and offer a reference for future large-scale atlas integration efforts. By harmonizing existing single-cell datasets, uniLUNG establishes a unified platform for advancing our understanding of lung cell biology.

We discovered three novel cell subpopulations through case studies with uniLUNG. First, our investigation of various disease contexts identified two distinct cell types—lymphoid monocytes and T-like B cells—that preferentially localize to specific lung conditions, revealing potential commonalities among these diseases and their unique characteristics compared to others. By mapping to external single-cell atlases, we verified the existence of these two new cell types. Second, our analysis of lung cancer unveiled a novel transitional malignant cell subpopulation, NSCLC-like SCLC, which is associated with the progression from non-small cell lung cancer (NSCLC) to small cell lung cancer (SCLC). We validated its presence in spatial transcriptomic samples of NSCLC and highlighted its complex roles in lung adenocarcinoma (LUAD) and lung squamous cell carcinoma (LUSC) through spatial co-localization. These new findings highlight uniLUNG as the most comprehensive single-cell reference atlas for the lung, generating novel insights that other existing single-cell lung atlases have not provided, thereby underscoring its value in the study of healthy and diseased lung tissue.

## RESULTS

### The construction the of uniLUNG through supervised integration

#### The representativeness of uniLUNG core datasets

We first constructed the core atlas and subsequently expanded it into a complete atlas by transforming labels to address computational challenges associated with processing millions of individual cells. We collected 18 public datasets ^3, 6, 8, 13, 14, 21–31^ from 508 donors across 12 lung statuses—including healthy, developmental, aging, cystic fibrosis (CF), chronic obstructive pulmonary disease (COPD), COVID-19, interstitial lung disease (ILD), idiopathic pulmonary fibrosis (IPF), large cell carcinoma (LCC), LUAD, LUSC, and SCLC—to develop the uniLUNG core atlas (Supplementary Table 1). To ensure representativeness and reference value, we balanced cell numbers across different lung conditions during dataset selection and prioritized datasets that provided complete original cell annotations and comprehensive metadata. This metadata includes donor characteristics such as age, sex, ethnicity, disease status, and smoking history, along with sample-specific information like type, collection site, experimental protocol, and sequencing platform (Figure 1a and 1b). We also retained critical information relevant to specific diseases, including COVID-19 severity and tumour staging.

**Figure 1.**
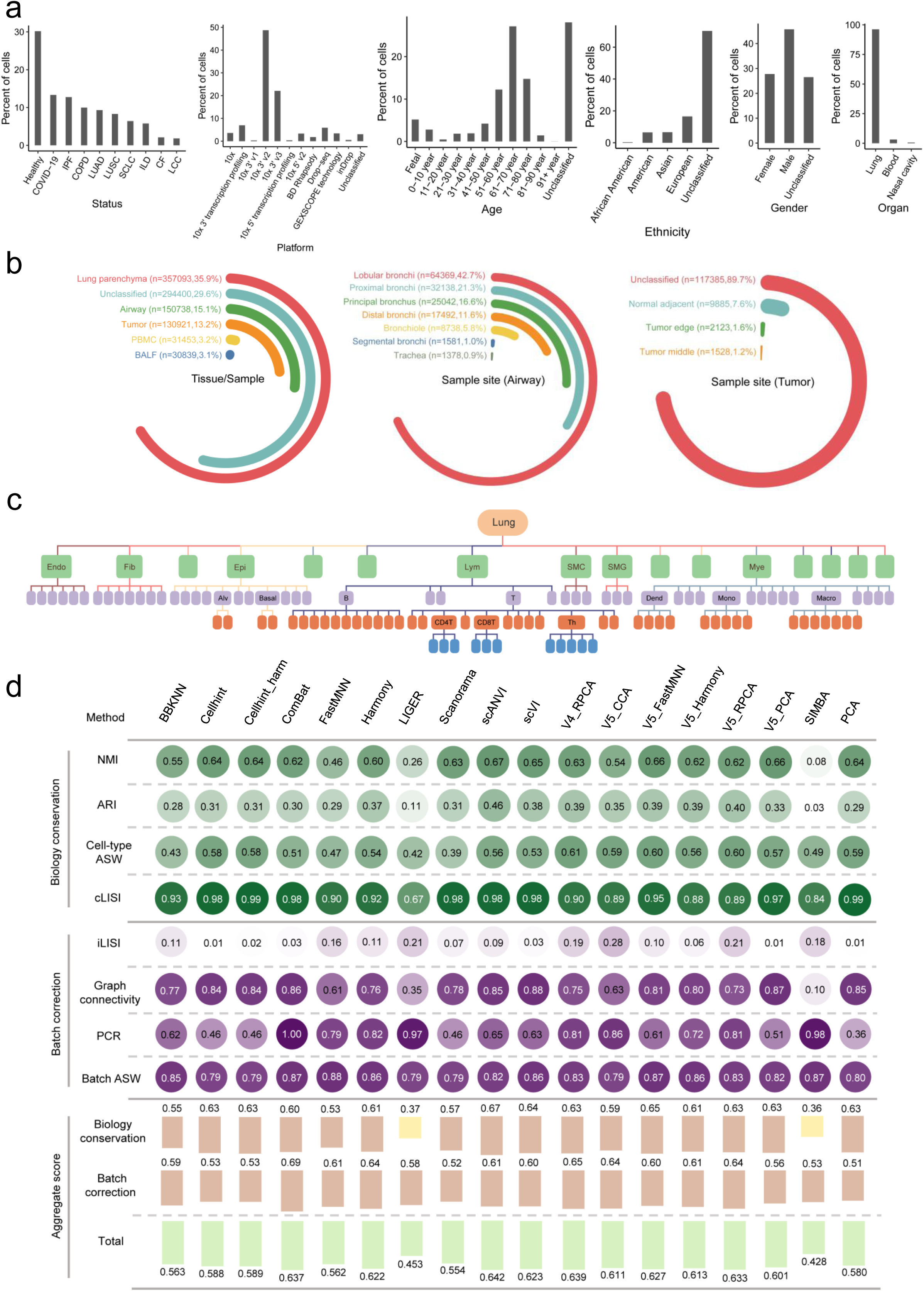
Composition of uniLUNG core and benchmarking of the dataset integration. **(a)** Demographic metadata of the uniLUNG core. **(b)** Anatomical information of the samples in uniLUNG core. **(c)** Hierarchical distribution of cell types in the uniLUNG core. **(d)** Integration benchmark test for the uniLUNG core. Batch correction scores and biological preservation scores display the average values for each respective metric. The final total score is a weighted combination of the biological preservation score (60%) and the batch correction score (40%).

#### A four-level universal hierarchical annotation framework (uHAF) in uniLUNG

Current single-cell lung studies often use different cell classification standards, which creates confusion when trying to understand findings from different studies. A large-scale single-cell integrative atlas provides a valuable opportunity for achieving standardized cell type annotation across experiments and datasets. Based on extensive existing research and consensus^32–36^, particularly the insights from LungMAP/CellCards ^4^, we have manually developed uHAF (https://lung.unifiedcellatlas.org/#/uHAFTree) for human lung cell-level annotation (Figure 1c), encompassing 120 known lung cell types (see Methods and Supplementary Table 2).

#### Data integration benchmarking optimizes embedding

After uniformly preprocessing the original matrices from all datasets, we obtained 995,444 cells to establish the uniLUNG core. To effectively mitigate batch effects while preserving biological differences, we evaluated 18 data integration strategies, encompassing 12 major and latest methods of data integration ^37–48^ (see Methods). For supervised integration methods requiring known cell type labels, we created uniform identity labels by mapping original cell annotations to the uHAF (Supplementary Table 3). We then assessed the effectiveness of these strategies via scIB ^49^, applying eight metrics that evaluate batch effect correction and biological variation preservation (Figure 1d). Based on these evaluations, we selected the top-performing method scANVI to construct the uniLUNG core.

#### Manual cell type re-annotation achieves maximum cell diversity with high accuracy

After data integration, we systematically re-annotated all cells in the uniLUNG core according to the uHAF. Our goal was to assign higher-resolution labels where feasible, leading to an initial division of the core into 9 main cell types (Figure 2a and Supplementary Figure 1a), which were further subdivided into 67 distinct cell types (Figure 2b-c, Supplementary Figure 1b and 2). While the re-annotation generally aligned with the original labels at the primary level (Figure 2d), we conducted a thorough evaluation of accuracy and diversity to validate the unified annotations.

**Figure 2.**
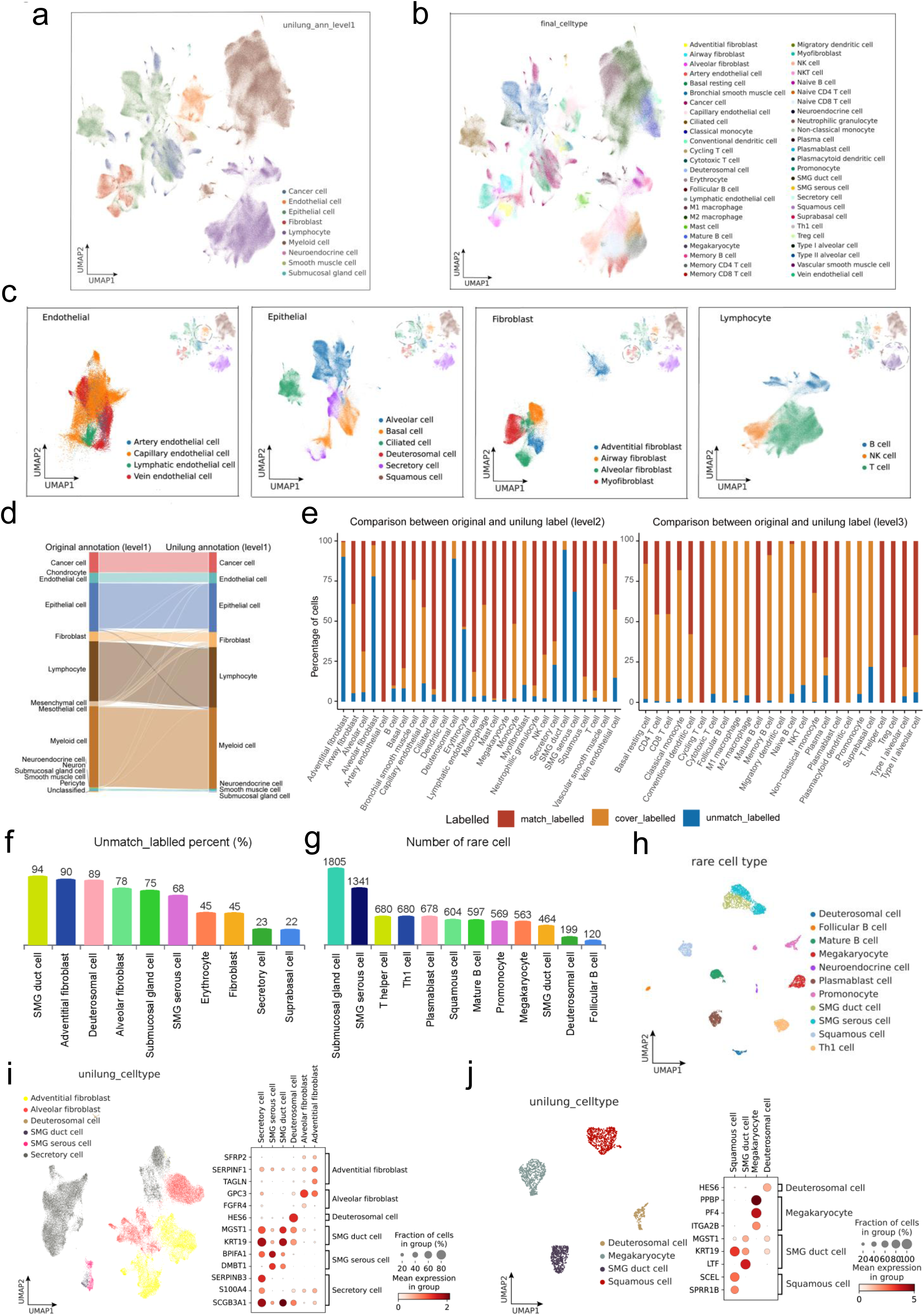
uniLUNG core achieved a reliable re-annotation based on uHAF. **(a)** UMAP of the uniLUNG core coloured by the re-annotation results at the first level. **(b)** UMAP of the uniLUNG core coloured by the final re-annotation labels. **(c)** Hierarchical annotation of selected main cell types in the uniLUNG core. **(d)**Comparison between the re-annotation and the original labels of uniLUNG core reveals good consistency at the first level of annotation. **(e)** Percentage of matching, unmatching, or covering labels in the second and fourth level of re-annotation relative to the original labels, calculated separately for each cell type. **(f)** Cell types with a unmatch_labelled percentage greater than 20% in the re-annotation compared to the original labels across all annotation levels. **(g)** Rare cell types and their quantities in the uniLUNG core re-annotation. Cell types with fewer than 2000 cells across all annotation levels are categorized as rare cells. **(h)**UMAP of all rare cell types. **(i-j)**UMAP of cell types at the second and third levels with the unmatch_labelled percentage greater than 20%, coloured based on the re-annotation. Gene expressions confirm the accuracy of the re-annotation results.

For accuracy, we classified the new annotations into three categories based on their correspondence with original labels: match_labelled, cover_labelled (different annotation granularity), and unmatch_labelled (Figure 2e and Supplementary Figure 3). We assessed the proportion of unmatched annotations across four annotation levels (Supplementary Tables 4-7), identifying 10 cell types with unmatch_labelled proportions exceeding 20% (14.9% of total types) as potentially misannotated (Figure 2f). Following an analysis of marker gene expression and input from multiple team members, we refined the re-annotation results (Figure 2i and Supplementary Figure 4a), effectively correcting biases and enhancing accuracy of the original annotations.

Regarding diversity, we flagged 12 cell types with fewer than 2000 cells (17.9% of all types) as potential rare cell types (Figure 2g and 2h), such as squamous cells, megakaryocytes, and submucosal gland cells. The expression profiles of their marker genes supported the re-annotations (Figure 2j and Supplementary Figure 4b). These rare cell types, reported in only a few studies, demonstrate the effectiveness of the uniLUNG core re-annotation in preserving cellular diversity. Overall, the re-annotation process highlights the advantages of integrating cell atlases from large-scale datasets, achieving maximum cellular diversity with high accuracy.

#### The creation of complete uniLUNG via label transfer from the core atlas

Based on the uniLUNG core, we utilized CellTypist ^50^ to expand the consensus annotation of cells to other datasets through label transfer (see Methods), ultimately generating a comprehensive, universally applicable transcriptomic atlas of human lung cells in both healthy and various disease states (uniLUNG: https://lung.unifiedcellatlas.org). This atlas integrates 71 published single-cell datasets covering 20 lung statuses (healthy, development, aging, and 17 diseases, see Methods and Supplementary Table 8), comprising 10,389,138 cells from 67 cell types obtained from 1944 donors, along with biological and demographic metadata such as age, sex, race, disease status, and sampling region. The construction of uniLUNG is founded on a unified information framework, encompassing high-quality data preprocessing, uHAF and CellTypist-based annotations, assembly and storage in the unified giant table (uGT) ^51^, and a user-friendly data portal for public access. Gene expression counts and metadata are standardized and integrated into the uGT database, facilitating cell-centric integration and comparative gene expression analyses across different cell types and diseases.

### uniLUNG reveals new cell types across health and multiple lung diseases

#### Identification of two distinct cell subpopulations across multiple diseases via uniLUNG

The human lung is an organ susceptible to a wide range of diseases, yet our understanding of cellular diversity across both healthy and diseased states remains limited. Existing studies have primarily focused on specific pulmonary diseases due to sample type and source constraints, lacking a comprehensive view of the lung’s cellular landscape across multiple diseases. Our integration of uniLUNG across multiple lung states offers an unparalleled resource for single-cell research in pulmonary diseases. To explore cellular similarities and differences between healthy and diseased lungs, we analysed 1,643,385 cells in uniLUNG from tissue samples representing 18 lung conditions (Supplementary Table 9). After correcting batch effects using Harmony ^41^, the cells were categorized into five major cell types (Figure 3a-b). Given the variation in donor numbers and cell counts across disease datasets, we applied scCODA ^52^ to assess and compare the relative abundance of cell types (excluding cancer cells) across different lung conditions. Epithelial cells, lymphocytes, and myeloid cells showed significant abundance differences between lung states (Figure 3c). Recognizing that epithelial malignancy in lung cancer may contribute to these differences, we isolated lymphocyte and myeloid subpopulations from the original data. After further batch correction with Harmony, we subdivided these into T cells, B cells, NK cells, macrophages, mast cells, and monocytes (Supplementary Figure 5a-b), revealing notable shifts in monocyte and B cell abundance across conditions (Supplementary Figure 5c).

**Figure 3.**
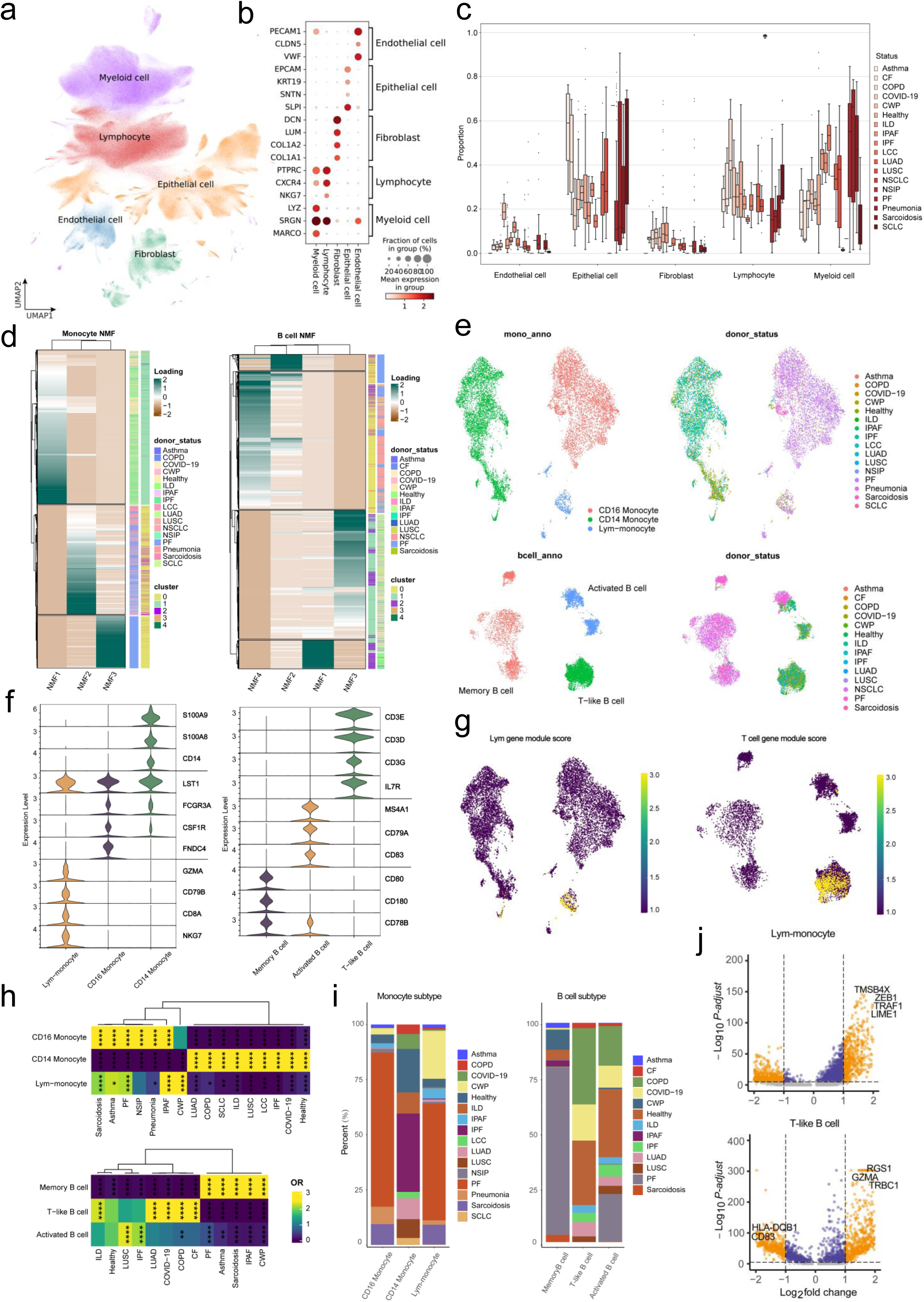
Analysis of lung health and multiple diseases. **(a-b)** Identification of major cell types. **(c)** Comparison of the abundance of major cell types in different lung states. **(d)** NMF analysis divided B cells and monocytes into 3 and 4 subpopulations, respectively. **(e)** UMAP visualization of B cell and monocyte subpopulations. **(f)** Characteristic gene expression in B cell and monocyte subpopulations. **(g)** lymphocyte feature gene set score in monocytes and T-cell feature gene set score in B cells, showed lymphocyte feature of Lym-monocyte and T cell feature of T-like B cell, respectively. **(h)** Distribution bias of B cells and monocyte subpopulations in different lung states. **(i)** The percentage of B cells and monocyte subpopulations derived from different lung states. **(j)** Gene differential analysis of T-like B cells and Lym-monocytes.

We utilized NMF and Louvain algorithms for unsupervised clustering of monocytes and B cells, identifying three distinct subpopulations (Figure 3d, Supplementary Figure 6 and 7a). Validation of these subpopulations was performed through the expression analysis of canonical markers for myeloid, monocyte, lymphocyte, and B cell lineages (Supplementary Figure 7b). Among the clusters, we identified a new monocyte subpopulation with high expression of lymphocyte markers, termed Lym-monocytes, and a new B cell subpopulation expressing T cell markers, termed T-like B cells. Both subpopulations were observed across various lung disease conditions (Figure 3e-f). Gene set enrichment analysis revealed that lymphocyte signature genes (e.g., CD79B, CD19, PTPRC, GZMA, CD8A, CD3D, IL7R, CXCR4, NKG7, CD8B) and T cell signature genes (e.g., CD3E, CD3G, CD8A, CXCL13, GATA3, IL7R, CD4) were highly enriched in Lym-monocytes and T-like B cells, respectively (Figure 3g).

#### Heterogeneity and characterization of the two distinct cell subpopulations

By calculating the odds ratio (OR) ^53^ and cell proportions across various lung conditions, we further highlighted the heterogeneity of monocyte and B cell subpopulations, particularly Lym-monocytes and T-like B cells, in different lung diseases (Figure 3h-i). For instance, CD14 monocytes were broadly distributed with consistent proportions across multiple lung states, while CD16 monocytes exhibited specific distributions that largely excluded CD14 monocytes but overlapped with Lym-monocytes in the diseases of PF, IPAF, and CWP. Memory B cells showed a preference for PF, IPAF, and CWP, while T-like B cells were broadly distributed across COPD, COVID-19, LUAD, ILD, and other lung diseases. Differential expression analysis revealed key marker distinctions between Lym-monocytes and T-like B cells. Lym-monocytes upregulated lymphocyte markers (e.g., TMSB4X, ZEB1, TRAF1, LIME1), whereas T-like B cells showed elevated expression of T cell markers (e.g., RGS1, GZMA, TRBC1) and downregulation of certain B cell markers (e.g., CD83, HLA-DQB1) (Figure 3j). GO enrichment analysis indicated that Lym-monocytes were enriched for lymphocyte- and T cell-related processes, while T-like B cells were associated with mitochondrial respiration and energy metabolism pathways (Figure 4a). We found a significant increase in the proportion of T-like B cells among all B cells in severe/critical COVID-19 patients compared to those with mild/moderate symptoms (Figure 4b). Previous studies have indicated T cell exhaustion in severe COVID-19 patients ^54^, which is associated with impaired mitochondrial respiration and oxidative stress ^55^. The increase in T-like B cells may be one of the repair mechanisms in response to these impaired functions in these patients. Additionally, the T-like B cell gene signature in LUAD tumour was highly correlated with a gene signature of exhausted T cells (Figure 4c). The concentrated distribution of T-like B cells in LUAD may reflect a remodelling of the tumour microenvironment (TME) in response to T cell exhaustion in tumour ^56^, potentially leading to a similar functional compensation mechanism. Subsequently, we used SCENIC ^57^ analysis to explore the differential transcription factor regulatory networks between monocyte and B cell subpopulations. In monocytes, FOS and ETS exhibited high activity in Lym-monocytes, with FOS also highly active in the other two monocyte subtypes (Figure 4d). Its downstream targets included diverse marker genes such as FCN1, PTPRC, MKI67, and SERPINA1 (Supplementary Figure 8b). ETS, in contrast, showed elevated activity specifically in Lym-monocytes, with its downstream targets largely representing lymphocyte-associated markers (Figure 4e). Moreover, the transcription factors and regulatory networks active in CD14 monocytes differed significantly from those in both Lym-monocytes and CD16 monocytes (Supplementary Figure 8). In B cells, each of the three B cell subpopulations exhibited distinct transcription factor activity. Notably, TFDP1 and KLF6 were highly active in T-like B cells, where their downstream targets included T cell-associated markers such as IL7R and CTSA (Supplementary Figure 9).

**Figure 4.**
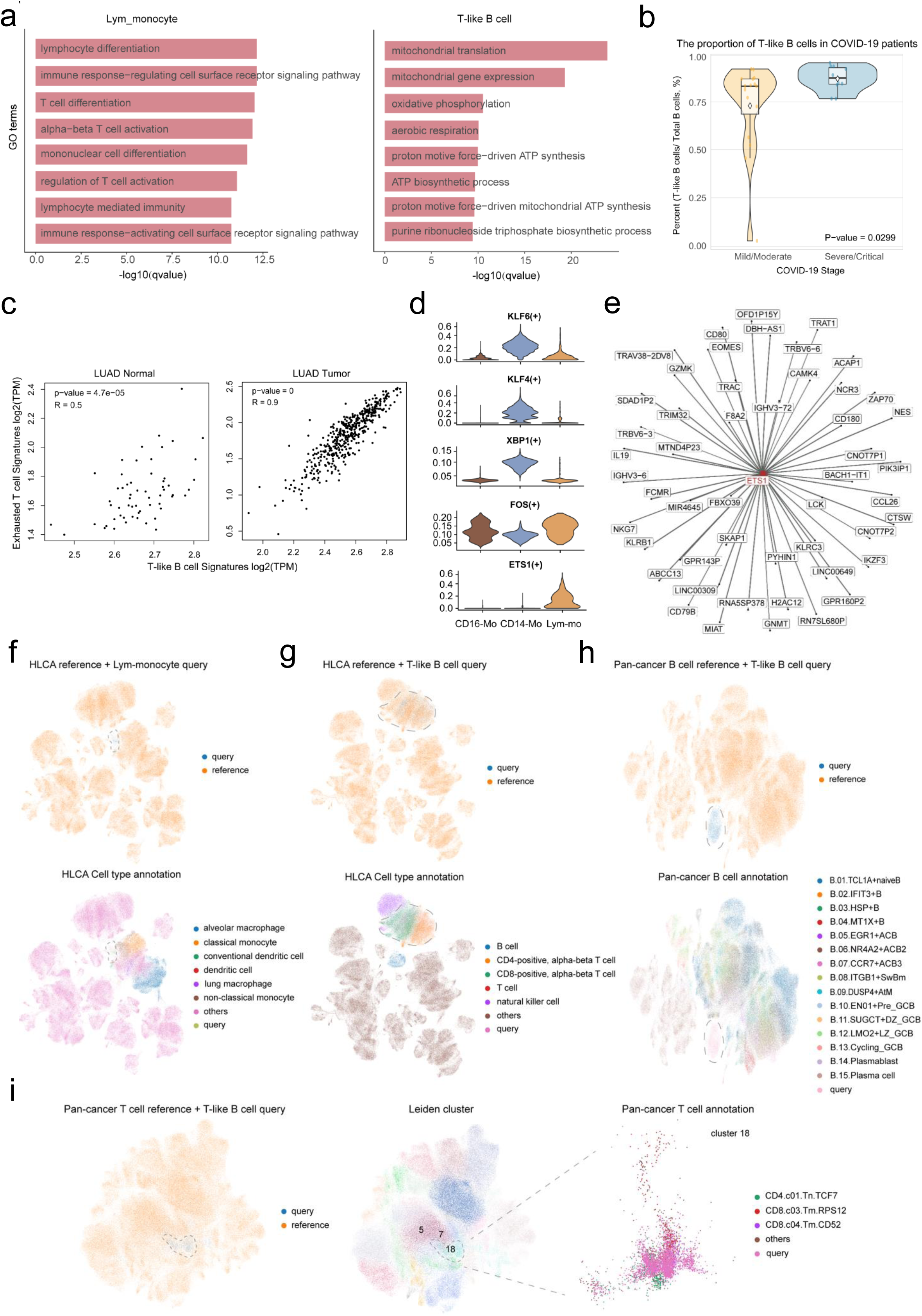
Characterization and verification of T-like B cell and Lym-monocyte. **(a)** GO enrichment analysis of T-like B cells and Lym-monocytes. **(b)** Comparison of the proportion of T-like B cells among total B cells between mild/moderate and severe/critical COVID-19 patients. **(c)** Correlation analysis between exhausted T cells and T-like B cell gene signatures, performed separately in normal and tumour samples of LUAD. **(d)**Differential activation of transcription factors across different monocyte subsets. **(e)** Transcription factor regulatory network centred on ETS1, which specifically activated in Lym-monocyte, highlighting its lymphoid characteristics. **(f)** UMAP visualization of Lym-monocytes mapped to the HLCA, highlighting surrounding cell types. **(g)** UMAP visualization of T-like B cells mapped to the HLCA, highlighting surrounding cell types. **(h)** UMAP visualization of T-like B cells mapped to the pan-cancer B cell atlas, displaying all cell types. **(i)** UMAP visualization of T-like B cells mapped to the pan-cancer T cell atlas. Unsupervised clustering shows T-like B cells primarily cluster in a distinct group, with surrounding cell types highlighted.

#### Validation of the new cell subpopulations by mapping to external datasets

To confirm the presence of Lym-monocytes and T-like B cells, we used scArches ^58^ to map them onto other existing lung atlases. In the HLCA ^21^, Lym-monocytes mapped close to non-classical monocytes, reinforcing their monocyte-like characteristics, while T-like B cells clustered with CD4 and CD8 T cells, reflecting their similarity to T cells (Figure 4f-g). Given their existence in lung cancer samples, we further mapped them using the LuCA ^13^. Lym-monocytes aligned with classical monocytes and macrophages, while T-like B cells localized near CD4, CD8, and regulatory T cells (Supplementary Figure 10a-b). We also projected T-like B cells onto a published pan-cancer B ^59^ and T cell atlas ^53^, which included lung cancer samples. Although T-like B cells exhibited some overlap with known B cell populations, they did not correspond to any established cancer-associated B cell subtypes (Figure 4h). In the T cell atlas, despite their close similarity to CD4/CD8 T cells, unsupervised clustering identified T-like B cells as a distinct subpopulation, emphasizing their divergence from existing T cell subsets (Figure 4i). These results reveal that T-like B cells and Lym-monocytes, as newly identified cell types, are widely present in existing datasets, despite not being prominently featured in previous lung atlas studies. This underscores the significant potential of uniLUNG, compared to existing lung atlases, in uncovering the diversity of lung cell types.

In summary, uniLUNG unveils the dynamic cellular landscape of the human lung across health and disease, particularly highlighting the heterogeneity of B cells and monocytes. The identification of Lym-monocytes and T-like B cells—two novel cell types—reveals their distinct characteristics and their preference for specific lung conditions, showcasing the unparalleled scope of uniLUNG in enhancing our understanding of lung cell diversity and cross-disease insights.

### uniLUNG unveils NSCLC-like SCLC subpopulation in NSCLC’s transformation to SCLC

#### Identification of novel transitional malignant cell subpopulation based on the lung cancer sub-atlas

Lung cancer, a leading cause of cancer-related deaths globally, is broadly classified into SCLC and NSCLC, the latter of which encompasses subtypes such as LUAD, LUSC, and LCC. LUAD, LUSC, and SCLC collectively represent the most prevalent types, comprising approximately 80-85% (LUAD and LUSC) and 10-15% of all lung cancer cases (https://www.cancer.org/cancer/types/lung-cancer), respectively. Tailored treatment approaches are necessary due to variations in the originating cells and clinical characteristics of different lung cancers. However, histological transition events directly contribute to alterations in lung cancer characteristics, leading to drug resistance and poor prognosis ^60, 61^. To explore the possible histological transformation events between different lung cancer subtypes, we analysed 290, 89, and 21 patients for LUAD, LUSC and SCLC from the lung cancer sub-atlas of uniLUNG, respectively, part of whom had tumour staging information (Supplementary Table 10). Initially, all 1,683,056 cells were categorized into five main cell types, with 310,555 epithelial cells further identified as basal cells, ciliated cells, secretory cells, AT1 cells, AT2 cells, and malignant cells (Figure 5a-c). Malignant cell identification was based on comprehensive marker gene expression, CNV scores, and cell differentiation potential (Figure 5c, Supplementary Figure 11-12).

**Figure 5.**
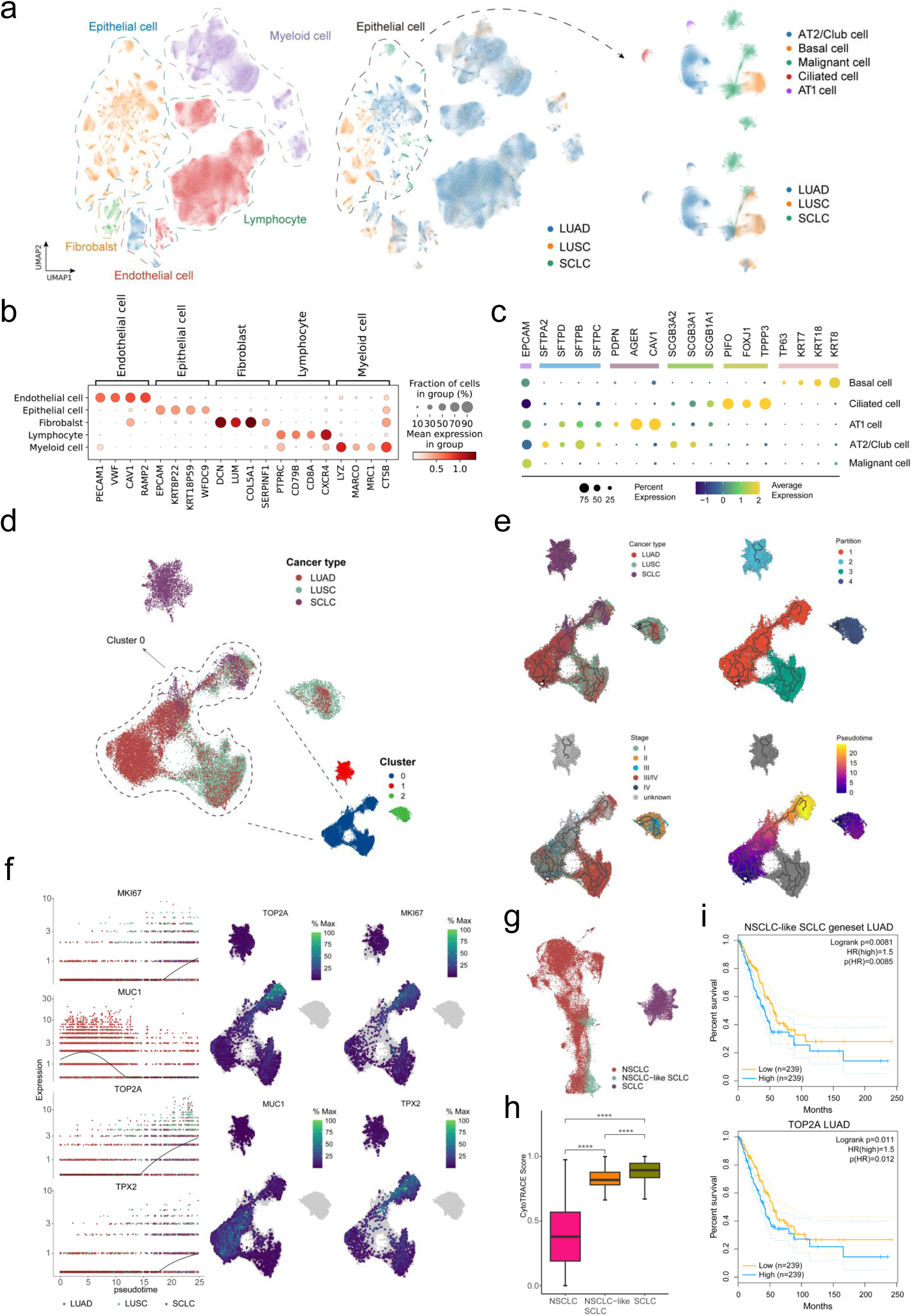
Single cell transcriptome analysis of lung cancer subtype transition. **(a-b)** All cells were initially classified into five major cell types, with epithelial cells further subdivided into five distinct subpopulations. **(c)** Marker gene expression analysis of epithelial cell subpopulations, showing that the malignant cell subset expresses few markers of normal epithelial subpopulations, except for EPCAM. **(d)** UMAP visualization of all malignant cells, coloured by cancer type. Unsupervised clustering revealed a cluster containing malignant cells from LUAD, LUSC, and SCLC patients, suggesting shared characteristics between malignant cells from different subtypes of lung cancer. **(e)** Trajectory analysis of all malignant cells identified four independent trajectories, distributed across different partitions. The largest partition encompasses trajectories representing malignant cells from LUAD, LUSC, and SCLC patients, which were established as the main trajectory. Based on clinical information, the direction of this main trajectory was determined to follow LUAD-LUSC-SCLC. **(f)** Genes showing significant expression changes along the trajectory. **(g)** UMAP visualization was reconstructed by extracting all cells from the main trajectory and an independent SCLC malignant cell subpopulation. A small subset of SCLC patient-derived malignant cells within the main trajectory displayed similarities to NSCLC malignant cells, with differences from the independent SCLC malignant cell subset, and were named the NSCLC-like SCLC subpopulation. **(h)** The differentiation potential of the NSCLC-like SCLC subpopulation lies between the NSCLC and SCLC malignant cell subpopulation. **(i)** Survival analysis was performed on LUAD samples from TCGA based on the NSCLC-like SCLC signature gene set and TOP2A expression. The results indicate that the NSCLC-like SCLC subpopulation is associated with poor prognosis in LUAD patients.

Clustering of all malignant cells revealed similarities between cancer cells from LUSC and LUAD, the two main NSCLC subtypes, while SCLC malignant cells formed a distinct cluster (Figure 5d), highlighting significant biological differences between SCLC and NSCLC. Previous studies suggest that histological transitions between lung cancer subtypes may contribute to resistance to targeted therapies ^61–65^. To investigate potential transitions between these subtypes, we performed trajectory analysis on malignant cells, which were divided into four distinct partitions, each representing an independent trajectory (Figure 5e). In the largest partition (Partition 1), malignant cells from all three lung cancer types coexisted, forming the primary transformation pathway (Figure 5e top left and top right). By integrating tumour staging information (Figure 5e bottom left), we identified the direction of this trajectory (Figure 5e bottom right), revealing an evolutionary path from LUAD to LUSC, and ultimately to SCLC (Figure 5e top left). In the early stages, LUAD-derived cells predominated, transitioning to LUSC cells, with a small subset of malignant cells ultimately emerging as SCLC in some patients (Figure 5e top left). Throughout this trajectory, proliferation-related genes such as MKI67, TOP2A, and TPX2 showed increased expression ^66–68^, while the metastasis-associated gene MUC1 decreased ^69^ (Figure 5f), indicating dynamic shifts in gene expression. To further explore malignant cell transitions between LUAD, LUSC, and SCLC, we separated all cells along the trajectory (Partition 1) and the independent SCLC subpopulation (Partition 2), and rebuilt their UMAP (Figure 5g). A distinct subset of SCLC malignant cells present on the trajectory was designated as NSCLC-like SCLC to distinguish it from the independent SCLC malignant cell population (Fig. 5g). Notably, this subpopulation exhibited a differentiation potential intermediate between NSCLC and SCLC cells, suggesting that the NSCLC-like SCLC subgroup represents a transitional state in the progression from NSCLC to SCLC (Figure 5h). Survival analysis based on the NSCLC-like SCLC gene signatures and TOP2A (which shows significant expression changes along the trajectory and is also present in the NSCLC-like SCLC gene signatures) in TCGA LUAD samples indicated that the NSCLC-like SCLC is associated with poor prognosis in LUAD patients (Figure 5i).

#### Validation and characterization of NSCLC-like SCLC cells in NSCLC patients at spatial resolution

We integrated spatial transcriptomic data from 12 tumour samples of 5 LUAD patients and 8 tumour samples of 3 LUSC patients ^70^. Cells were categorized into normal, tumour-like, and tumour cells using unsupervised clustering, CancerFinder ^71^ for malignant cell identification, and EPCAM expression (a carcinoma cell marker that is abnormally highly expressed in a variety of epithelial cancers ^72, 73^) (Figure 6a-b). In each sample, EPCAM expression was spatially concentrated in regions identified as tumour and tumour-like cells (Figure 6c, Supplementary Figure 13). Scoring each sample with the NSCLC-like SCLC signature gene set revealed tumour cells with similar characteristics in some LUAD and LUSC tumour regions, confirming the presence of the NSCLC-like SCLC subpopulation (yellow highlighted regions) in NSCLC (Figure 6d, Supplementary Figure 14).

**Figure 6.**
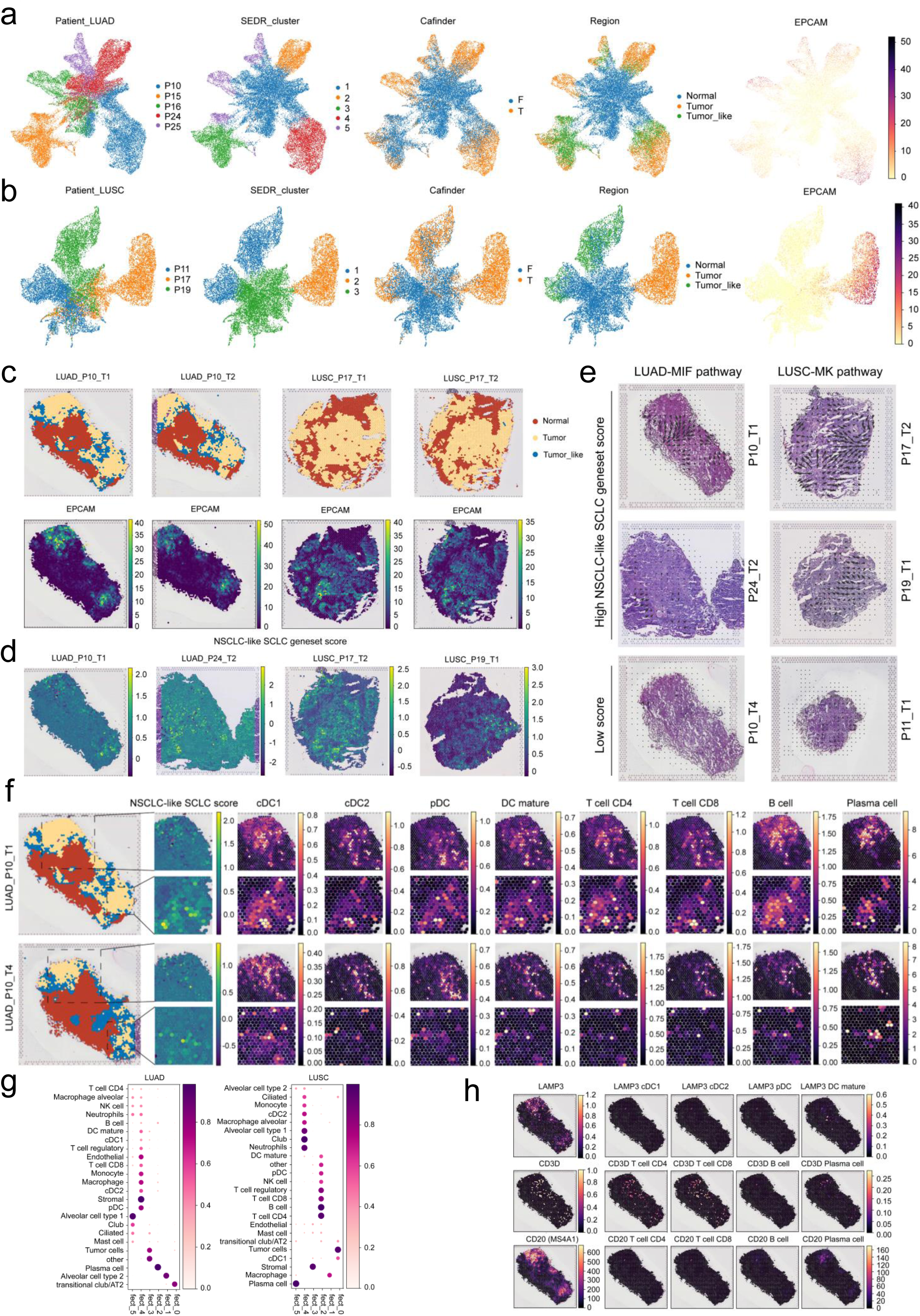
Spatial transcriptome verification and analysis of NSCLC-like SCLC subpopulation. **(a-b)** Spatial transcriptomic samples of LUAD and LUSC were integrated separately. Based on clustering results from SEDR and CancerFinder identification of malignant cells, all cells were classified into tumour cells, normal cells, and tumour-like cells, with EPCAM expression tested. **(c)** Spatial partitioning of regions containing tumour cells, normal cells, and tumour-like cells in LUAD and LUSC samples. **(d)** Scoring of NSCLC-like SCLC characteristic gene sets in LUAD and LUSC samples. Results revealed NSCLC-like SCLC subpopulation characteristics (highlighted in yellow) in the tumour cell regions of some LUAD and LUSC samples, suggesting the presence of the NSCLC-like SCLC subpopulation. **(e)** LUAD and LUSC samples with or without prominent NSCLC-like SCLC features were grouped into high and low scoring categories. Spatial communication analysis of signal flow in these groups indicated that in both LUAD and LUSC high-scoring samples, signals related to tumour progression were observed flowing from tumour cell regions to surrounding normal regions. **(f)** Two tissue sections from the same LUAD patient exhibited varying levels of NSCLC-like SCLC features, with the sample showing prominent NSCLC-like SCLC features displaying a more pronounced Immune cell infiltration in the tumour cell regions. **(g)** NMF analysis revealed the clustering of multiple immune cells at the same factor level in both LUAD and LUSC. **(h)** Marker expression of key cell types in tertiary lymphoid structures (TLS) in LUAD samples with prominent NSCLC-like SCLC features suggests the presence of TLS in these samples.

In samples exhibiting a distinct NSCLC-like SCLC subpopulation, we identified several gene expression patterns (Supplementary Figure 15a). Notably, cancer-associated fibroblast (CAF) markers ^74^ (Pattern 6) were consistently detected across multiple LUAD and LUSC samples (Supplementary Figure 15a-b), suggesting widespread CAF infiltration in NSCLC. Basal cell markers ^16, 75^ (Pattern 4) and markers for type II alveolar and club cells ^16, 75^ (Pattern 9) were enriched in LUSC and LUAD (Supplementary Figure 15a-b), respectively, reflecting their distinct malignant origins ^76, 77^. Further spatial analysis of ligand-receptor co-localization revealed that ligands such as COL1A1, COL1A2, FN1, and LUM co-localize with ITGB1 in tumour-adjacent normal regions, indicating active extracellular matrix (ECM)-receptor interactions ^78, 79^ in NSCLC (Supplementary Figure 16 and Supplementary Table 11). To explore the spatial role of the NSCLC-like SCLC subpopulation in NSCLC, we applied COMMOT ^80^ to analyse intercellular communication in tumour samples with high and low NSCLC-like SCLC scores across LUAD and LUSC (Supplementary Table 12-13). In LUAD samples with pronounced NSCLC-like SCLC features, MIF signalling was observed flowing from malignant to normal regions (as indicated by the black arrows), a pathway closely linked to patient survival ^81^ (Figure 6e). Similarly, in LUSC, the MK pathway exhibited a comparable flow pattern (Figure 6e), indicating its role in promoting epithelial-mesenchymal transition (EMT) and tumour proliferation ^82^.

#### The effect of NSCLC-like SCLC cells on NSCLC patients in spatial dimension

Using the LuCA as a reference, we applied cell2location ^83^ to deconvolute cell types in LUAD and LUSC samples. The analysis showed increased abundance of cDC1, cDC2, and pDC in LUSC tumour regions, whereas LUAD tumours were enriched with cDC1, cDC2, pDC, DC mature, T cell CD4, T cell CD8, B cells and Plasma cells (Supplementary Figure 17-20). Notably, plasma cells were concentrated within the tumour in LUAD, while in LUSC, they predominantly resided in the surrounding normal tissue (Supplementary Figure 21a). Additionally, the presence of the NSCLC-like SCLC subpopulation significantly elevated immune cell abundance in both LUAD and LUSC (Figure 6f, Supplementary Figure 21b). NMF analysis further confirmed distinct immune infiltration patterns between these tumour types (Figure 6g). Importantly, lymphocytes enriched at the tumour core or margin in LUAD were identified as key components of tertiary lymphoid structures (TLS) ^84, 85^. Marker gene analysis confirmed the presence of TLS-associated cell types, such as DC-LAMP (LAMP3), CD3D-expressing T cells (CD3D), and B cells (CD20) ^86, 87^ (Figure 6h). This suggests that the NSCLC-like SCLC subpopulation may contribute to TLS formation in LUAD.

By analysing the relationship between spatial spot composition and gene expression variability within cell types, we found that the NSCLC-like SCLC subpopulation significantly reshapes intercellular communication in LUAD and LUSC (Supplementary Figure 22). In LUAD samples with high NSCLC-like SCLC characteristics, CD4 T cells showed a loss of dependency on type II alveolar cells, tumour cells, and transitional club/AT2 cells, instead relying on cDC1 cells, indicating a shift in communication partners (Supplementary Figure 22a). Additionally, while mast cells and monocytes were critical communication hubs in both LUAD groups, their intercellular dependencies were significantly reduced in the low-scoring group (Supplementary Figure 22a). In LUSC, cDC2 cells in the high-scoring group exhibited strong dependencies on macrophages and stromal cells, whereas these dependencies were nearly absent in the low-scoring group (Supplementary Figure 22b). We then performed receptor effect analysis to highlight how sending cells influence gene expression in specific receptor cell types, providing context for differential gene expression in certain cellular couplings (Supplementary Figure 23-24). For CD4 T cells and cDC2 cells, which exhibited significant changes in intercellular dependency in LUAD and LUSC, we assessed the impact of the NSCLC-like SCLC subpopulation on these interactions through sender similarity analysis. We found that LUAD samples with lower NSCLC-like SCLC scores retained sender profiles of specific lineages more effectively (Supplementary Figure 23), whereas the opposite was observed in LUSC (Supplementary Figure 24). Finally, given the distinct interactions between tumour cells and other cell types in the LUAD group (Supplementary Figure 22a), we further analysed how tumour cells, as single senders, affect gene expression in different NSCLC-like SCLC scoring contexts (Supplementary Figure 25).

Overall, using the lung cancer sub-atlas of uniLUNG, we reconstructed the malignant cell transition trajectory among LUAD, LUSC, and SCLC, identifying a novel subpopulation of malignant cells with NSCLC-like characteristics—termed NSCLC-like SCLC. This subpopulation likely represents a critical transitional state during the progression from NSCLC to SCLC. The presence of this subpopulation was validated in spatial transcriptomic samples of tumours from LUAD and LUSC patients, where we extensively analysed the gene expression profiles and intercellular signalling networks within these tumours. Subsequent spatial analyses further revealed the distinct impacts of this subpopulation on immune cell infiltration and cell type dependencies in LUAD and LUSC patients. These findings provide significant insights into the histological transition mechanisms in lung cancer and offer potential therapeutic targets for patients with transformed SCLC, while also underscoring the value of uniLUNG as a comprehensive data resource in human lung disease research.

## DISCUSSION

We constructed uniLUNG, the largest single-cell atlas of the human lung, encompassing approximately 10.4 million cells from 71 datasets, which span 67 cell types and capture the cellular landscape across healthy and diseased lungs. The atlas supports visual exploration and preliminary analysis, enabling cell-level retrieval and comparison under various conditions, as well as targeted selection of lung cells for personalized analysis. uniLUNG demonstrates substantial utility as an integrated resource, overcoming the limitations of individual datasets and supporting broad applications in lung health and disease research. Our analyses of various lung diseases based on uniLUNG have identified novel cell subpopulations, including Lym-monocytes and T-like B cells. Additionally, we discovered a critical transitional malignant cell subpopulation involved in the NSCLC to SCLC transition process, confirming its spatial distribution through spatial transcriptomics and further elucidating its complex impact on LUAD and LUSC.

B cells and monocytes are pivotal in the immune responses to various lung diseases ^88–91^. We have identified two novel subsets, Lym-monocyte and T-like B cell, which display distinct characteristics and preferentially emerge in specific lung pathological states. In patients with PF, CWP, IPAF, and sarcoidosis, Lym-monocyte enrichment correlates with a decrease in CD14 monocytes and an increase in CD16 monocytes. The decline in CD14 monocytes is associated with reduced lung function ^92, 93^, while the rise in CD16 monocytes typically indicates heightened inflammation and CD4 T cell activation ^94, 95^. Lym-monocyte enrichment may also reflect close interactions with lymphocytes. In LUAD and COVID-19 patients, we found that the emergence of T-like B cells is associated with T cell exhaustion, potentially serving a compensatory role. T-like B cells are also notably concentrated in CF, COPD, and ILD patients, suggesting the presence of T cell exhaustion and a similar mitochondrial function repair mechanism in these conditions. Previous studies have confirmed the presence of exhausted T cells in COPD ^96, 97^ and ILD ^98^ patients. Furthermore, the formation of T-like B cells contributes to the development of a complex immune microenvironment and influences disease progression through frequent interactions with T cells ^99–101^. Importantly, we validated the unique characteristics of Lym-monocytes and T-like B cells through comparisons with several existing cell atlases (HLCA, LuCA, Pan-caner B cell atlas and Pan-cancer T cell atlas), confirming their classification as novel populations. By integrating data from various lung disease studies, our findings highlight common features that exist among different lung diseases, aiding in establishing connections between them. This may provide a preliminary molecular basis for applying existing therapies targeted at specific diseases to others, or for developing new treatments applicable to multiple lung diseases.

We identified a distinct malignant cells subpopulation, designated as the NSCLC-like SCLC cell, in patients with SCLC. This subset demonstrates intermediate proliferation and differentiation capabilities between mature NSCLC and SCLC cells, potentially representing a transitional malignant state during the conversion from NSCLC to SCLC, and is associated with acquired EGFR-TKI resistance ^61, 63, 64^. We detected the NSCLC-like SCLC subset in spatial transcriptomic samples from LUAD and LUSC, suggesting an increased risk of SCLC transformation in these cases. In NSCLC patients harbouring this subset, tumours showed a high concentration of cancer-associated fibroblasts (CAFs), which facilitate tumour progression through various pathways ^102, 103^. The extracellular matrix (ECM), recognized as a key tumour component and therapeutic target ^104, 105^, was also actively involved. Our analysis indicated that NSCLC-like SCLC activates the MIF pathway in LUAD, potentially driving tumour progression via downstream ERK1/2, AMPK, and AKT pathways ^106, 107^. Conversely, in LUSC, this subset may enhance tumour proliferation, survival, and anti-apoptotic mechanisms through the MK pathway and its downstream Notch, JAK/STAT, and MAPK pathways ^82^. Furthermore, this subset is associated with increased lymphocyte infiltration in NSCLC tumours and the formation of tertiary lymphoid structures (TLS) in LUAD. While TLS generally indicate a favourable prognosis in cancers ^108^, including NSCLC ^85^, alterations in their composition or spatial organization may correlate with poor outcomes ^86, 109^, warranting further investigation into the specific impacts of NSCLC-like SCLC-driven TLS formation in LUAD. Cell dependency analysis suggests that NSCLC-like SCLC may cause greater structural disruption in LUAD compared to LUSC, a finding supported by sender similarity analysis. In summary, our integration of spatial multi-omics data positions the NSCLC-like SCLC subpopulation as a potential intermediate stage in the conversion from NSCLC to SCLC, highlighting its general impact in NSCLC patients and specific effects in LUAD and LUSC. These insights may assist in developing warning systems for SCLC transformation in NSCLC patients and lay the groundwork for new treatment strategies targeting therapy-resistant NSCLC arising from SCLC transformation.

Although uniLUNG provides broader diversity and extensive coverage, it is constrained by the limitations inherent to current single-cell studies in population sampling, disease spectrum, and biological insights. For instance, the donors in uniLUNG are primarily from European (37.5%) and Asian (45.7%) populations, lacking comprehensive representation of more diverse population groups. The majority of uniLUNG data (∼90.3%) represent lung cells from healthy individuals and patients with lung cancer, ILD, or COVID-19, while data from other diseases remain limited. Integrating large-scale atlases also poses computational challenges, as significant experimental design variations across datasets lead to batch effects. Direct batch correction of large expression matrices incurs high computational costs, yet differential expression analysis without batch correction introduces biases from non-biological factors. The quality of uniLUNG heavily depends on integration strategies, which directly influence its ability to capture lung cell subtype heterogeneity—reflected by the substantial proportion of unclassified cell types in the uHAF. To strengthen uniLUNG’s capacity to comprehensively characterize human lung cells, our ongoing goals include continuous supplementation with new single-cell transcriptomic and multimodal data and refinement of cell type annotations.

Overall, as the largest unified reference atlas of human lung single-cell data and an interactive data portal, uniLUNG enables users to explore cellular diversity and phenotypic differences in the human lung under healthy and various disease states, revealing new insights not previously identified in earlier integrated atlases. It facilitates in-depth exploration of lung diseases, aiming to address key issues in lung biology and pathology, and provides valuable resources for decoding lung diseases and exploring therapeutic approaches on a broader scale.

## MATHODS

### Data collection and processing

To builg uniLUNG, we collected cells from various research studies and public databases, including GEO, HCL, HCA, HLCA, DISCO, TISCH, LuCA, HTAN, CancerSEA, CancerSCEA, LungMAP, HuBMAP, UCSC Cell Browser, and Single Cell Expression Atlas, ^1, 2, 4, 13, 14, 16, 19, 21, 110–115^. The datasets cover a diverse range of populations, including healthy individuals (fetal, adult and elderly population) and patients with multiple lung diseases (Asthma, CF, COPD, COVID-19, CWP, ILD, IPAF, IPF, NSIP, PF, LUAD, LUSC, LCC, SCLC, NSCLC, pneumonia and sarcoidosis). For each dataset, we collected the expression matrices and accompanying metadata, which includes information about sequencing technologies, donor age and gender, ethnicity, organ type, sample tissues, sampling site, disease status, smoking, covid and tumour stage, and original annotations. In uniLUNG, we refer to this information as metadata.

To eliminate the impact of different preprocessing methods on data integration, we chose to only use datasets that provide the raw matrices to construct uniLUNG. Additionally, datasets from different sources may have used different versions of the reference genome during the alignment process, resulting in differences in the number and symbol of genes across datasets. To assemble all the data into uniLUNG, we used a list of 43,878 genes (Supplementary Table 14) approved by the HUGO Gene Nomenclature Committee (HGNC) to convert gene names from datasets that were originally named with Ensemble ID to HGNC symbols. Genes that were not detected in the sequencing were filled with zeros. Subsequently, we performed quality control using a standard pipeline described by Hao Y et al in 2021 ^40^. We removed cells with fewer than 200 expressed genes or with mitochondrial gene percentages exceeding 20%. Based on gene counts versus UMI counts, we manually determined the number of unique molecular identifiers (UMIs) for each sample, ranging from 50,000 to 200,000 UMIs. DecontX ^116^ was applied for environmental RNA correction, with a maximum allowable environmental contamination level set to 0.2. Subsequently, we selected 2000 highly variable genes (HVGs) and computed PCA based on 30 latent dimensions. Finally, we utilized DoubletFinder ^117^ to identify and remove doublets from the dataset.

### Benchmarking of data integration

Benchmarking for data integration methods was performed in uniLUNG core. It consists of 18 public datasets covering 11 lung states, sourced from 12 different sequencing platforms and technologies. To mitigate batch effects arising from diverse donor samples and sequencing technologies, we opted to partition large datasets encompassing multiple sequencing platforms and experimental designs. Ultimately, the uniLUNG core was subdivided into 38 independent datasets, representing batches requiring correction. We mapped the original labels to the uHAF for consistent cell naming. Subsequently, each cell’s original label was assigned a corresponding cell type in the uHAF, aiming for higher resolution whenever possible. This newly assigned label was utilized for supervised data integration tasks.

We selected the latest and widely-used methods for benchmarking. These included methods implemented in Python such as CellHint, Combat, Harmony, Scanorama, SIMBA, scVI, and scANVI, as well as methods executed in R like BBKNN, LIGER, SeuratV4 (RPCA), SeuratV5 (CCA and RPCA), and FastMNN. Additionally, we conducted control tasks in both Python and R using only PCA-based computations, labelled as PCA and SeuratV5_PCA, respectively, to assess performance without employing any specific integration methods. Some methods could be run with different parameters or versions, such as CellHint and CellHint_harmonize, SeuratV4_RPCA and SeuratV5_RPCA. Furthermore, certain methods were already integrated into SeuratV5, including SeuratV5_FastMNN and SeuratV5_Harmony. We treated these integrated methods as new tasks for individual testing purposes. Due to memory usage constraints, the CCA method based on SeuratV4 was excluded from the evaluation. For all tasks run in R, SeuratV4 was utilized by default for methods not explicitly specified. The conversion of dataset formats between R and Python was facilitated by SeuratDisk.

SIMBA utilized all the genes for computation while other integration tools used in this work were conducted on 2000 highly variable genes (HVGs) with default parameters. These HVGs were selected to ensure consistent processing across methodologies. Among all strategies, scANVI and scVI utilize the original counts of HVGs as input, while the other methods use scaled matrices normalized with default parameters. Both scANVI and CellHint_harmonize methods permit guidance of data integration through known cell type labels. We mapped the original labels to the uHAF for consistent cell naming. Following this mapping, each cell’s original label was assigned a corresponding cell type in the uHAF, aiming for higher resolution whenever possible. This newly assigned label was employed for supervised data integration tasks.

In the benchmarking, apart from cell-type LISI (cLISI) and integration LISI (iLISI), other metrics (NMI, ARI, Cell type ASW, Graph connectivity, PCR comparison, and Batch ASW) were computed using the default parameters of the scIB ^49^ framework. The cLISI was employed to assess the accuracy of cell clustering, while the iLISI characterized the effectiveness of data integration. These metrics were computed using the R package LISI ^41^ in the Python environment. We calculated cLISI values for batch and clustering, and iLISI values for cell annotation and clustering. The LISI scores range from 1 to N, where N represents the number of batches. For iLISI, a value of 1 indicates complete separation, with higher iLISI values indicating better mixing effects. Conversely, for cLISI, a value of 1 signifies optimal re-clustering after fusion, with higher values suggesting poorer preservation of biological variance post-mixing. Thus, we normalized these scores to the range of 0-1, where higher values indicate better performance. The calculation formula is as follows:

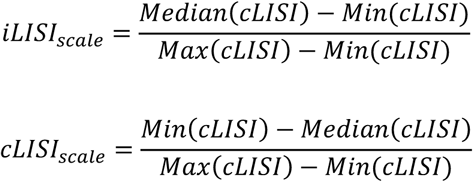

We evaluated the quality of each integration using eight metrics, with four assessing batch correction quality and four quantifying the preservation of biological variance post-integration. Batch correction and biological conservation scores were calculated as the respective averages of the corresponding metrics. The overall score was then determined by a weighted average of 0.4:0.6 for batch correction and biological conservation scores, respectively.

### Integration and manual cell type re-annotation of uniLUNG core

Based on benchmark results, we opted for scANVI with the highest overall score for integration of the uniLUNG core. It was executed using the unified original cell labels named through uHAF and default parameters. Following data integration, we computed the nearest neighbour graph based on 30 latent dimensions with neighbours set to k = 30. Subsequently, the graph underwent the first round of Leiden clustering at a resolution of 0.2. Leveraging uHAF, we assigned primary level labels to all cells, encompassing endothelial cells, epithelial cells, fibroblasts, lymphocytes, myeloid cells, neuroendocrine cells, smooth muscle cells, mucous gland cells, and cancer cells, with cancer cells retaining their original annotations.

Subsequently, except for neuroendocrine and cancer cells, each cell subtype underwent computation of a new nearest neighbour graph using scANVI’s 30 latent dimensions, with k set to 30. The new graph underwent a second round of Leiden clustering at a resolution of 0.3. Based on the clustering outcomes, each cell subtype was further delineated, completing the second-level annotation, and the same strategy was employed for the third and fourth rounds of Leiden clustering

### Cell type label transfer from the uniLUNG core to extension

After completing the integration and annotation of the uniLUNG core, we leveraged CellTypist to transfer its labels to the uniLUNG extension, following best practices in large-scale data label transfer established by CellTypist (https://celltypist.readthedocs.io). Specifically, we first sampled the uniLUNG core, setting the criterion to not exceed 20,000 cells per cell type (mode=’each’, n_cells=20000), while ensuring balanced representation of each cell type during sampling (balance_cell_type=True). Subsequently, we employed a rapid training process with a limited number of iterations, setting parameters as follows: n_jobs=10, max_iter=10, use_SGD=True. From the resultant initial models obtained through rapid training, we extracted the top 300 significant genes from each cell type, ranked based on their absolute regression coefficients relevant to the given cell type. These genes, after cross-cell type combinations, were further utilized in model training. During downstream model training, we opted for a traditional logistic regression framework over SGD to achieve unbiased probability estimation, albeit at the expense of longer runtime (max_iter=100). Notably, we trained two models using the uniLUNG core as a reference, differing in whether cancer cells were included. The model excluding cancer cells was employed for label transfer across all datasets except lung cancer, while the model including cancer cells was specifically used for annotating lung cancer datasets. Upon model training completion, we performed individual annotations on all datasets within the uniLUNG extension, with parameters set to defaults except for setting majority_voting=True. Following label transfer, the cell labels in the uniLUNG extension serve as valuable references for personalized cell type annotation in user analyses based on data from uniLUNG. Furthermore, these cell labels can offer guidance for supervised data integration before users proceed with downstream analysis using standard data from uniLUNG. As demonstrated in all case studies, we conducted targeted batch effect correction using scANVI on the involved datasets. The two CellTypist models are available from our atlas website (https://lung.unifiedcellatlas.org/download/model_all.pkl and https://lung.unifiedcellatlas.org/download/model_noca.pkl).

### Data standardization and database architecture

During data preprocessing, we standardized gene expression matrices for each dataset to ensure consistency across all genes involved. Correcting batch effects posed another significant challenge. The interpretation of batch effects may vary depending on research questions and study designs. While some studies view batch effects as interfering factors to be eliminated, others consider them biologically relevant. Existing batch effect correction methods are typically tailored for specific downstream analysis tasks, highlighting the importance of selecting appropriate correction methods based on research questions and analytical objectives. Therefore, we normalized gene expression values across all datasets solely based on library size to address differences caused by sequencing depth, ensuring comparability of gene expression values across batches. Other sources of variance were preserved along with the expression matrix. Results obtained from the online platform and downloaded data undergo only normalization, with users required to select suitable methods for data integration to mitigate batch effect influences in further analyses. Additionally, we provide pre-integrated atlases for direct download and downstream analysis, including the uniLUNG core and six sub-atlases

All expression information and metadata are stored in a unified manner using the uGT deployed in a cloud storage repository. The uGT is specifically designed as a NoSQL database for column-based storage, enabling fast retrieval of high-dimensional data. This database overcomes the limitations of traditional SQL databases in terms of the number of columns, addressing the issue of storage for cells with a large number of features (at least 40,000-50,000 columns). We process all data into a standardized format, where each cell is stored as an independent row, with columns 1-43878 containing the gene expression values for that cell, and the descriptive information from the metadata added in subsequent columns.

### Web portal construction and data availability in uniLUNG

We introduced a graphical user interface portal(https://lung.unifiedcellatlas.org), providing users with access to the data. The data processing workflow in uniLUNG and the basic structure of uHAF and uGT database are consistent with the hECA framework ^51^. The data portal website was built using the JavaScript-based Vue framework, and the visualization of statistical data results was achieved using the Echarts plugin. To promote widespread usage of uniLUNG in forthcoming single-cell lung studies and to enhance its utility as a reference and guidance tool, we offer various data access modalities to cater to diverse user requirements through download interface. This encompasses the integrated core atlas and sub-atlases that can be directly employed for downstream analysis, as well as the capability to freely filter cells based on specific conditions and to acquire corresponding expression data. It is crucial to note that the latter data, obtained through filtering, are normalized based on sequencing depth, and users will need to conduct their own data integration aligned with their research objectives prior to analysis. Additionally, the download section also provides all data and comprehensive information on uHAF (https://lung.unifiedcellatlas.org/#/uHAFItems).

### Atlas analysis for multiple lung diseases

#### Data integration and main cell type annotation

We used Harmony ^41^ for data integration while removing batch effects between datasets. Following integration, we computed the nearest neighbour graph using 3,000 highly variable genes based on 30 principal components, with neighbours set to k = 30. After an initial round of Leiden clustering at a resolution of 0.1, all cells were classified into the following categories: endothelial cells (PECAM1, CLDN5, VWF), epithelial cells (EPCAM, KRT19, SNTN, SLPI), fibroblasts (DCN, LUM, COL1A2, COL1A1), lymphocytes (PTPRC, CXCR4, NKG7), and myeloid cells (LYZ, SRGN, MARCO). Subsequently, we performed a second round of clustering on lymphocytes and myeloid cells at a resolution of 0.2, leading to the identification of macrophages (CD163, MRC1, MARCO), mast cells (GATA2, MS4A2, TPSAB1, TPSB2), monocytes (FCN1, VCAN, SRGN), B cells (CD74, CD79B, CD80), NK cells (KLRD1, NKG7, GNLY), and T cells (CD3E, CD3G, CD3D).

#### Cell abundance calculation

In the analysis of lung health and multiple diseases, significant disparities exist among datasets representing different lung statuses, primarily stemming from imbalances in donor numbers, sample sizes, and cell quantities. These discrepancies prevent us from directly comparing changes in cell proportions across different lung states to obtain accurate results. Therefore, we opted to employ scCODA to compute and compare changes in cell abundance. Cell abundance reflects the extent of change for each cell type relative to a reference cell, which remains relatively stable across different lung statuses. In scCODA, we set lung statuses and individual donors as covariates, utilizing the automatically computed results (endothelial cells) from scCODA as the reference cell type (reference_cell_type=“automatic”). Additionally, we set the false discovery rate (FDR) to 0.05.

#### Data sampling of Monocytes and B cells

We extracted monocyte and B cell subsets from the raw data, filtering out samples with low cell counts. This process resulted in 108,813 monocytes and 244,857 B cells. To remove batch effects between samples, we used Harmony and performed unsupervised clustering at a resolution of 0.01. To accommodate data size limitations for subsequent analyses while preserving the original embedding space features, we applied the SketchData function in Seurat for uniform cell sampling, using the ‘LeverageScore’ method. Ultimately, we retained approximately 10% of the monocytes (11,000 cells) and about 5% of the B cells (12,500 cells).

#### NMF analysis of Monocytes and B cells

We performed non-negative matrix factorization (NMF) on the sampled monocyte and B cell populations using the R package NMF ^118^. The logarithmically transformed original expression matrix served as the input data. We selected the standard “brunet” option and conducted 50 iterations, testing rank values from 2 to 8. The optimal rank was determined by identifying the rank corresponding to the maximum change in the cophenetic value across different ranks. For monocytes, the optimal rank was found to be 3, while for B cells, it was 4.

#### Cell type annotation and AUCell score calculation of Monocyte and B cell subpopulations

Based on the Leiden clustering and NMF results, monocytes were categorized into CD16 monocytes (FCGR3A, CSF1R, FNDC4, LST1), CD14 monocytes (CD14, S100A8, S100A9), and lymphocyte-like monocytes (CD79B, CD8A, GZMA, NKG7). B cells were divided into memory B cells (CD79B, CD80, CD180), activated B cells (CD79A, MS4A1, CD83), and T cell-like B cells (CD3D, CD3E, IL7R, CD3G). We then combined lymphocyte characteristic gene sets (CD79B, CD19, PTPRC, GZMA, CD8A, CD3D, IL7R, CXCR4, NKG7, CD8B) and T cell characteristic gene sets (CD3E, CD3G, CD8A, CXCL13, GATA3, CD3D, IL7R, IL2RB, CD8B, CD4) to score gene modules for both monocyte and B cell subpopulations.

#### Odds ratios (OR) calculation

According to the method of Zheng et al. ^53^, the distribution preference of monocytes and B cell subsets in different pulmonary states was calculated.

#### Differential expression analysis

First, we used the R package Libra ^119^ to aggregate counts in each sample (as pseudo-bulk), applying edgeR’s generalized linear model and likelihood ratio test to assess differences in cell subsets while accounting for covariates (batch effects). We included donor ID as a covariate in our differential expression analysis, setting the design in edgeR ^120^ as ‘model.matrix(∼donor_ID + group)’, where donor ID represents batch effects and group indicates different cell subsets. Additionally, we employed Seurat’s FindMarkers function to calculate differential genes for Lym-monocytes and T-like B cells, selecting overlapping genes from the edgeR analysis as the final differential gene list.

#### Enrichment analysis

We conducted GO enrichment analysis using clusterProfiler ^121^. The parameters of pvalueCutoff and qvalueCutoff were both set to 0.01.

#### Correlation analysis of signatures

The signatures of T-like B cell (top 30) and Exhausted T cell (HAVCR2, TIGIT, LAG3, PDCD1, CXCL13, TIM3, PD1, LAYN) ^55, 122^ were selected as gene set, and LUAD tumour samples and normal tissue samples in TCGA were selected for correlation analysis based on GEPIA2 ^122^.

#### Transcription factor analysis by SCENIC

We applied SCENIC ^57^ to analyse active transcription factors in monocytes and B cells. The co-expression networks were constructed, and then the potential target genes of each transcription factor were identified by GRNBoost based on co-expression. DNA-motif analysis was conducted using RcisTarget to detect potential transcription factor binding motifs. And the regulatory activity of each transcription factor in each cell was scored by AUCel.

#### Reference query mapping for Lym-monocytes and T-like B cells

We utilized scArches ^58^ to map Lym-monocytes and T-like B cells to reference datasets. For Lym-monocytes, we selected HLCA and LuCA as reference sets, while for T-like B cells, we added pan-cancer B cell and T cell atlases ^53, 59^ as additional references. Due to the substantial difference in cell numbers between the sampled Lym-monocytes and T-like B cells and the reference datasets, we uniformly downsampled all reference datasets to 200,000 cells, excluding those with fewer than 200,000 cells. These reference sets were then used to train the scVI model, with the number of highly variable genes set to 3,000 and 500 iterations performed. We extracted expression matrices of the same genes from Lym-monocytes and T-like B cells to train the scVI model on the query dataset, also running 500 iterations. Finally, we linked the reference and query datasets within the same embedding space based on the model.

### Atlas analysis for lung cancers

#### CNV score estimation

We utilized the R package inferCNV ^123^ to compute copy number variation (CNV) events across all epithelial cells, employing a cutoff of 0.1. Epithelial cells from healthy lung tissues within the uniLUNG core were chosen as the reference. To quantify the occurrence of CNV events across all regions within cells, we normalized the gene expression levels of each cell to a range of -1 to 1 as described by Peng J et al in 2019 ^124^. Then, we computed the squared sum of the normalized values as the CNV score, indicating the extent of CNV events within each cell across all regions.

#### Evaluation of cell differentiation potential

For each cell type or cell subpopulation, CytoTRACE ^125^ was used to evaluate its differentiation potential, generating a UMAP visualization coloured by age group, as well as a boxplot representing the overall differentiation potential across different cell groups.

#### Trajectory construction

Pseudotime analysis was conducted using Monocle3 ^126^ to investigate the differentiation relationship among all malignant cells from LUAD, LUSC and SCLC patient. Gene exhibiting significant changes over pseudotime were identified through the graph_test function, with q value < 1e-4. We used the plot_genes_branched_pseudotime function to plot differential gene expressions.

#### Survival analysis

LUAD samples in TCGA were selected for survival analysis based on GEPIA2 ^122^.

#### Preprocessing and integration of spatial transcriptomics data

We performed basic filtering and normalization of spatial transcriptomics data from tumour samples of LUAD and LUSC patients using Scanpy ^127^. The filtering criteria included a maximum count of 50,000, less than 20% mitochondrial counts, and a minimum of 50 cells per gene. Based on 2,000 highly variable genes, we applied the SEDR ^128^ algorithm specifically developed for spatial transcriptomics to compute the embedding layer, which was then utilized for data integration via Harmony.

#### Identification of spatial regions

We utilized the mclust_R function in SEDR to cluster the LUAD and LUSC datasets, while employing the Cancer-Finder ^71^ algorithm to identify malignant spots. Based on clustering results and the identification of malignant cells, we categorized all spots into normal, pre-malignant, and malignant regions, with consideration of EPCAM expression levels.

#### Analysis of spatial gene expression patterns and ligand-receptor co-localization

To assess the widespread impact of the NSCLC-like SCLC subset on LUAD and LUSC tumours, we employed SpaGene ^129^ to identify gene expression patterns in samples exhibiting high NSCLC-like SCLC characteristics, setting nPattern to 10. We then used the SpaGene_LR function to evaluate ligand-receptor co-localization in each sample, filtering for shared ligand-receptor pairs between LUAD and LUSC samples.

#### Identification of spatial communication signal flow

We analysed communication signal flow in samples with and without high NSCLC-like SCLC characteristics in LUAD and LUSC using COMMOT ^80^. All procedures were conducted following the official guidelines.

#### Spatial mapping of cell types using cell2location

We selected LUAD and LUSC samples from LuCA as reference datasets. After removing all mitochondrial genes, we applied the filter_genes function for further filtering (cell_count_cutoff=5, cell_percentage_cutoff2=0.03, nonz_mean_cutoff=1.12) to prepare for spatial mapping and cell type annotation. The filtered reference dataset was then used to train a regression model, with ‘cell_type_major’ and ‘dataset’ as batch and label variables, respectively, and ‘platform’ as a covariate. The regression model was trained on all cells in the dataset with default parameters, achieving convergence within 250 epochs.

For the subsequent spatial mapping, we set the expected number of cells per spot to 30 and the regularization parameter for RNA detection sensitivity variation within slides or batches to 20 (default). The model was trained on the complete LUAD and LUSC datasets, converging after 40,000 iterations. We extracted cell abundances mapped to spatial coordinates using the 5th percentile of the posterior distribution. Additionally, we calculated cell type-specific expression for each gene at each spatial location, providing input for NCEM cell communication analysis.

#### Non-negative matrix factorization based on spatial mapping

Building on the cell2location mapping results, we employed non-negative matrix factorization (NMF) to further identify spatial co-localization of cell types. After training within a factor range of 3-10, we selected 6 factors for cell compartment identification and visualization in the LUAD and LUSC datasets.

#### Analysis of intercellular dependency as a spot composition function

We used the NCEM method ^130^ to assess the influence of spot/ecological composition on all genes. To focus on biologically relevant genes, we selected gene sets described in WikiPathways from the molecular signature database (MSigDB), yielding 6,716 genes. All 24 cell types in LuCA’s ‘cell_type_major’ variable were included in the NCEM analysis. Following type coupling analysis on the filtered dataset, we further dissected these couplings based on specific gene effects, including the impact of all senders on a single receiver (receiver effect analysis) and the influence of a single sender on all receivers (sender effect analysis). We retained dependencies involving at least 500 differentially expressed genes, with a q-value < 0.05 and absolute log fold change > 1 (1.5 for LUSC). Finally, sender similarity analysis was performed to characterize the sender profiles of CD4 T cells in LUAD patients and cDC2 cells in LUSC patients under different conditions.

## DECLARATIONS

### Ethics approval and consent to participate

Not applicable.

### Data availability

The data discussed in this publication can be found in uniLUNG (https://lung.unifiedcellatlas.org), and all the originally published data are also available from the GEO repository with the following codes in Supplement table 8 or ‘Reference’ of uniLUNG.

### Code availability

Codes for reproducing this work are available at https://github.com/Poison-pill/unilung.

## Acknowledgements

This publication is part of the Human Cell Atlas – www.humancellatlas.org/publications/. The graphical abstract was created with https://www.biorender.com/.

## DECLARATION OF GENERATIVE AI AND AI-ASSISTED TECHNOLOGIES IN THE WRITING PROCESS

During the preparation of this work, the authors used ChatGPT in order to proofread the manuscript. After using this tool, the authors reviewed and edited the content as needed and take full responsibility for the content of the publication.

