## Supplementary Figures for "A universal single-cell transcriptomics atlas of human lung decodes multiple pulmonary diseases"

**Supplementary Figure 1: Cell types in uniLUNG core.**
**Supplementary Figure 2: Re-annotation of uniLUNG core based on uHAF.**
**Supplementary Figure 3: Comparison of the uniLUNG core re-annotation with the original labels.**
**Supplementary Figure 4: Marker genes expression profile for some potential misannotated and rare cell types.**
**Supplementary Figure 5: Heterogeneity of B cells and monocytes between different lung states.**
**Supplementary Figure 6: NMF analysis of B cells and monocytes.**
**Supplementary Figure 7: Identification of B cell and monocyte subpopulations.**
**Supplementary Figure 8: Transcription factor analysis of monocytes.**
**Supplementary Figure 9: Transcription factor analysis of B cells.**
**Supplementary Figure 10: Mapping of cell subpopulations using LuCA as reference.**
**Supplementary Figure 11: Estimation of copy number variation events in epithelial cells.**
**Supplementary Figure 12: CNV score and differentiation potential estimation.**
**Supplementary Figure 13: Spatial region division and EPCAM expression in each sample.**
**Supplementary Figure 14: Scoring of NSCLC-like SCLC subset characteristic gene sets in LUAD and LUSC samples.**
**Supplementary Figure 15: Identification of shared gene expression patterns in samples with high NSCLC-like SCLC features.**
**Supplementary Figure 16: Identification of shared LRI in samples with high NSCLC-like SCLC features.**
**Supplementary Figure 17: Cell type annotation of spatial transcriptome samples of LUAD-P10.**
**Supplementary Figure 18: Cell type annotation of spatial transcriptome samples of LUAD-P15 and LUAD-P16.**
**Supplementary Figure 19: Cell type annotation of spatial transcriptome samples of LUAD-P24 and LUAD-P25.**
**Supplementary Figure 20: Cell type annotation of spatial transcriptome samples of LUSC-P17 and LUSC-P19.**
**Supplementary Figure 21: Differences in spatial distribution of immune cells.**
**Supplementary Figure 22: Differences in intercellular dependence between samples of LUAD and LUSC with high and low NSCLC-like SCLC feature scores.**
**Supplementary Figure 23: Sender effect analysis of LUAD samples with high and low NSCLC-like SCLC feature scores.**
**Supplementary Figure 24: Sender effect analysis of LUSC samples with high and low NSCLC-like SCLC feature scores.**
**Supplementary Figure 25: Receiver effect analysis of LUAD samples with high and low NSCLC-like SCLC feature scores.**

a

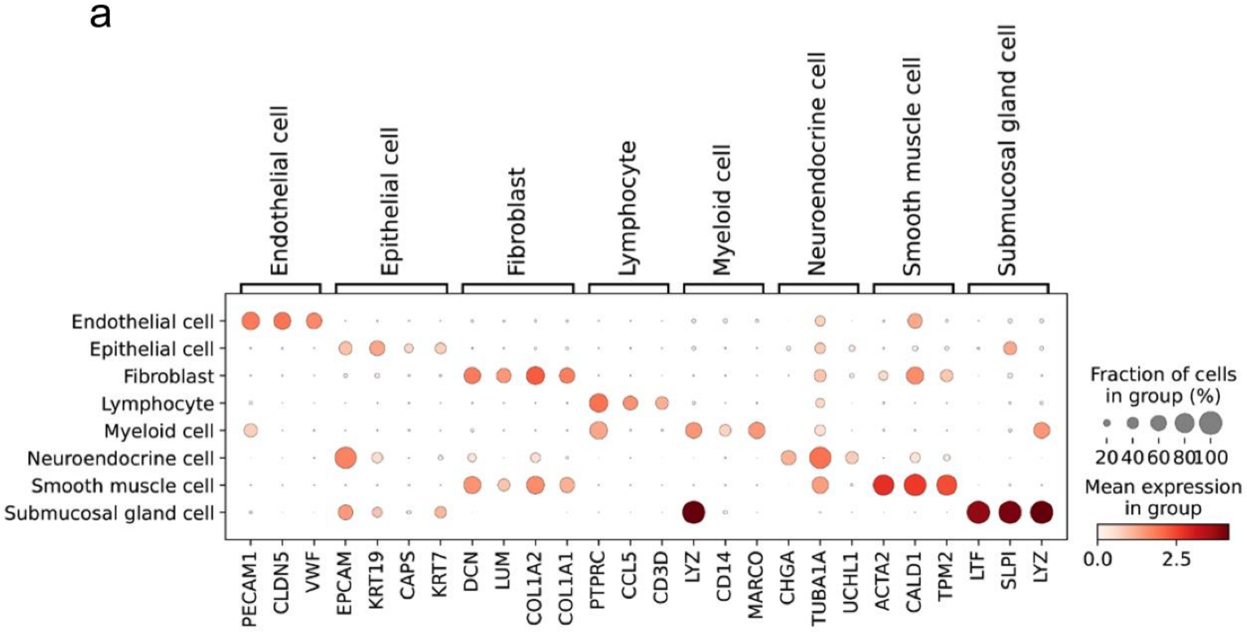



b

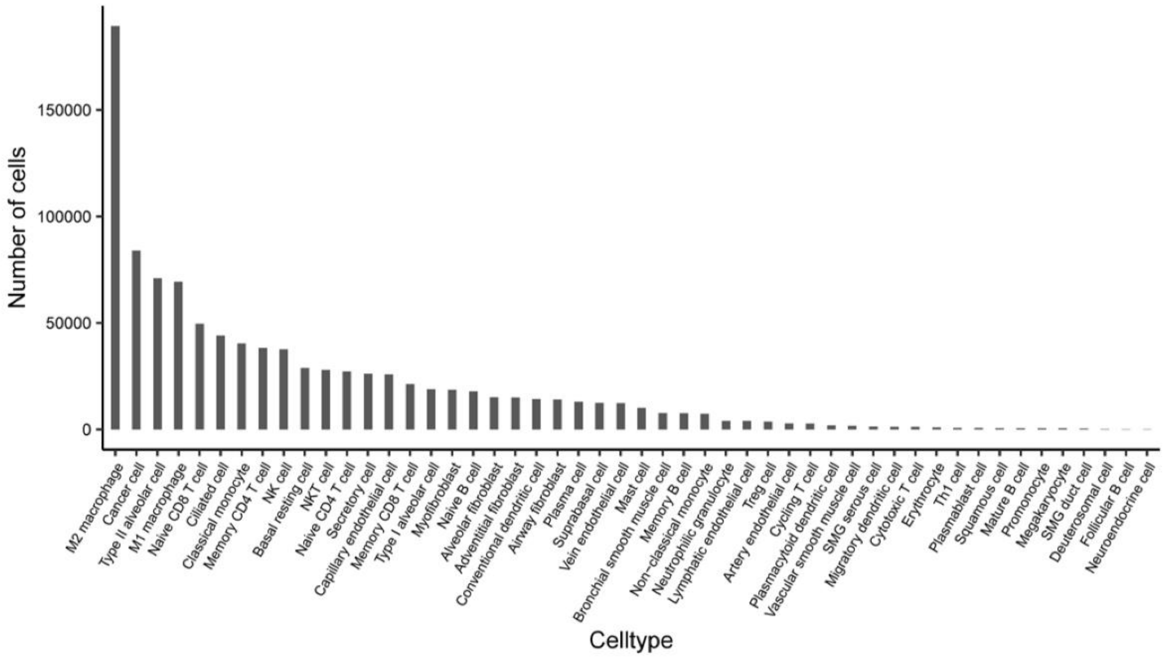



**Supplementary Figure 1: Cell types in uniLUNG core.**

**(a)** The expression profiles of marker genes for major cell types within the uniLUNG core (the first level of annotation). **(b)** Cell numbers for each cell type within the uniLUNG core.

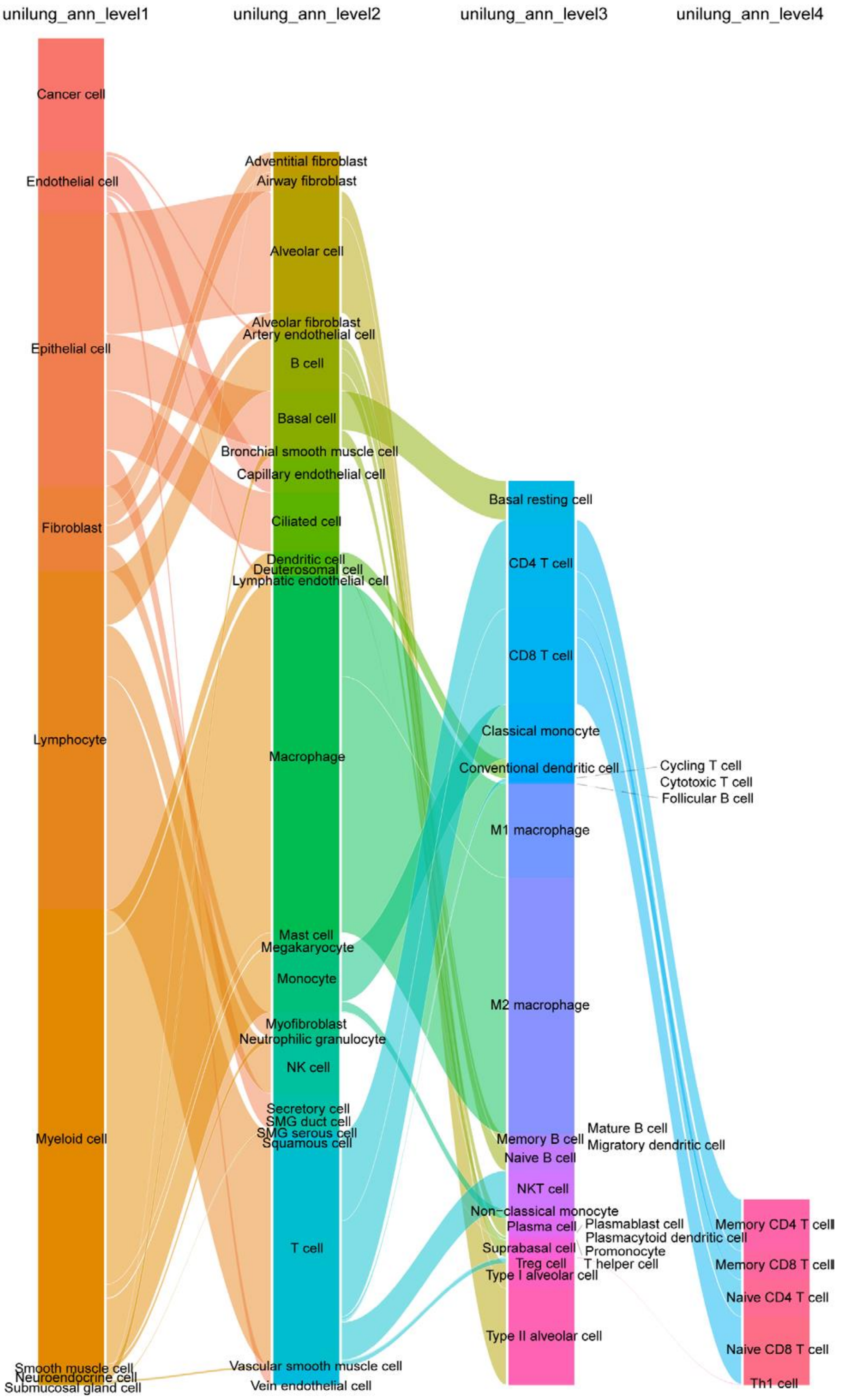


**Supplementary Figure 2: Re-annotation of uniLUNG core based on uHAF.** All cells within uniLUNG core were assigned cell type labels through hierarchical annotation, reaching a maximum resolution at the fourth level.

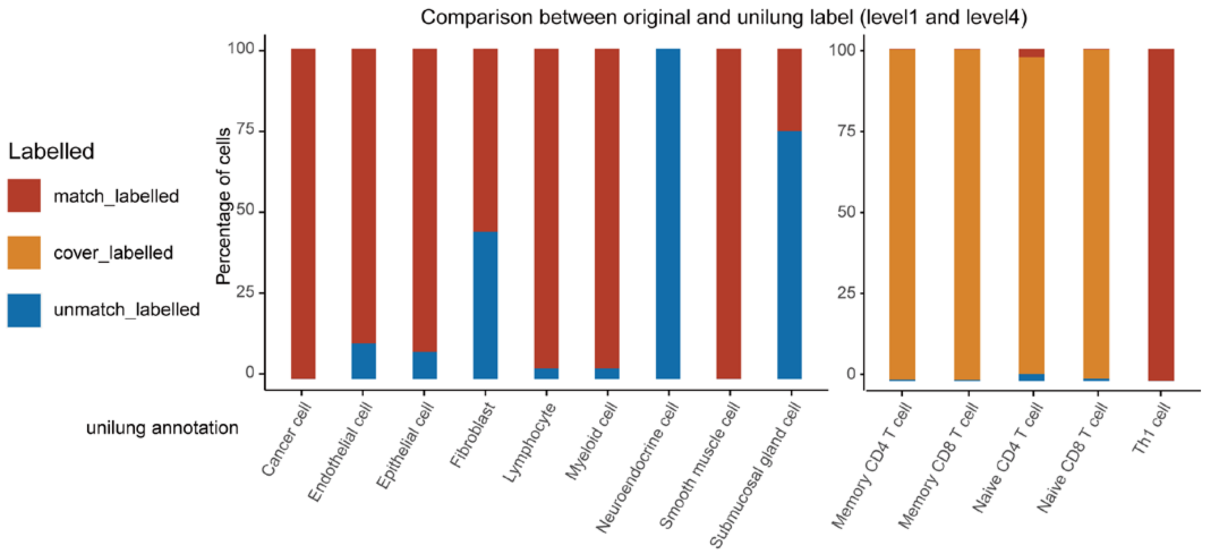


**Supplementary Figure 3: Comparison of the uniLUNG core re-annotation with the original labels.** Percentage of matching, unmatching, or covering labels in the first and fourth level of re-annotation relative to the original labels, calculated separately for each cell type.

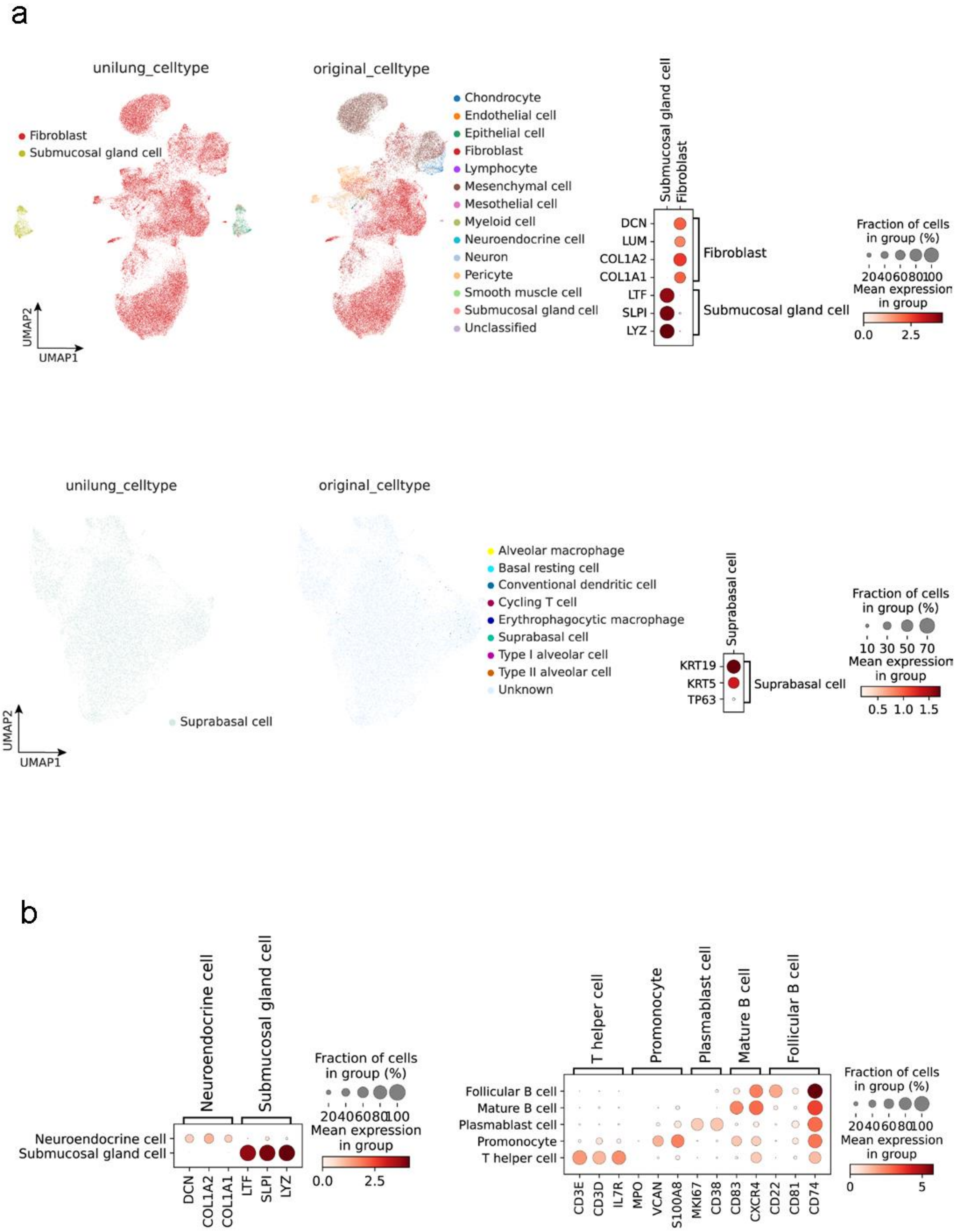


**Supplementary Figure 4: Marker genes expression profile for some potential misannotated and rare cell types.** **(a)** UMAP plots and expression profiles of marker genes for fibroblasts, submucosal gland cells and suprabasal cells, with UMAP plots color-coded based on re-annotation and original annotation. **(b)** Expression profiles of marker genes for select rare cell types.

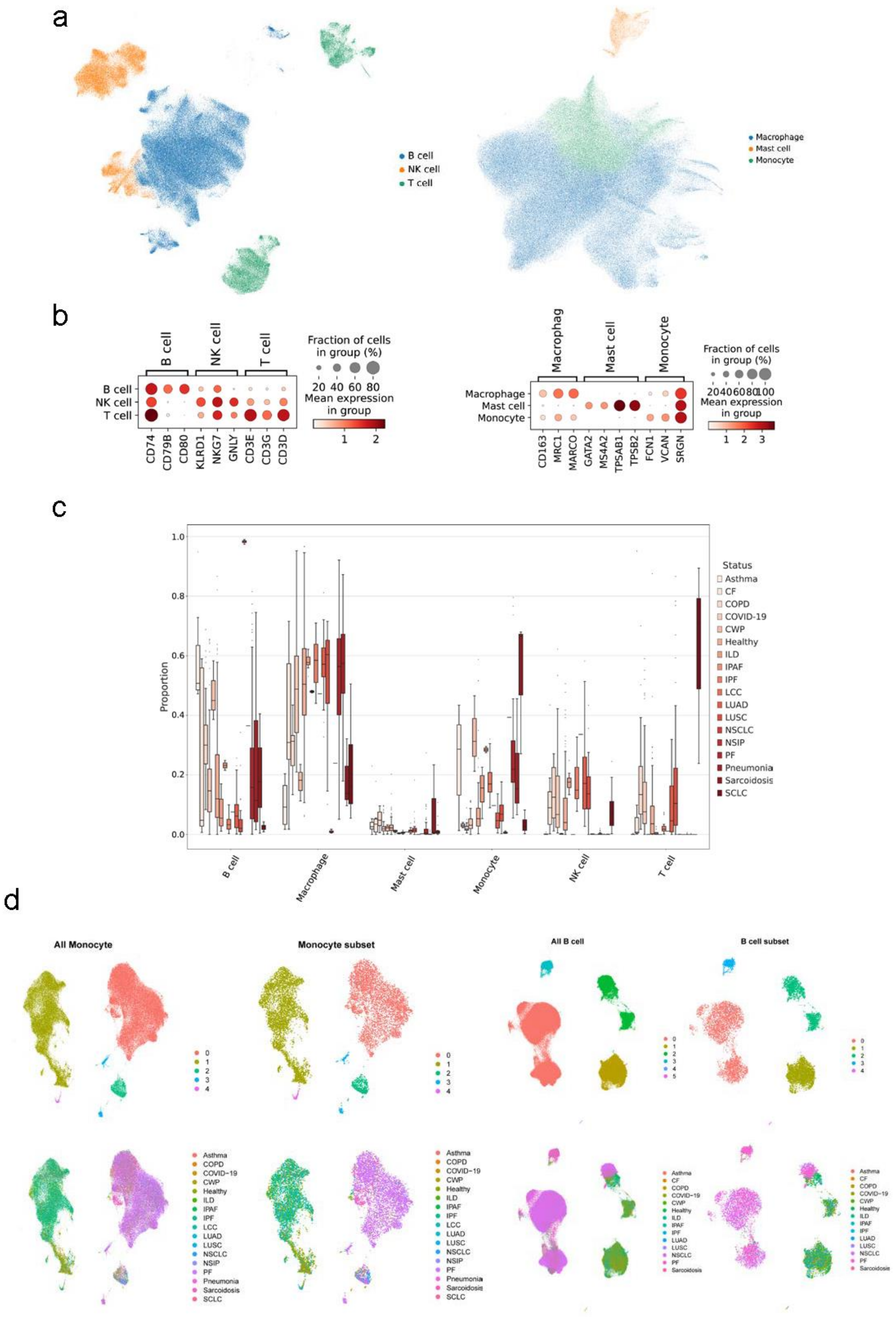


**Supplementary Figure 5: Heterogeneity of B cells and monocytes between different lung states.**

**(a-b)** Cell type annotation of lymphocytes and myeloid cells. **(c)** Differential abundance of lymphocyte and myeloid subpopulations across lung statuses. **(d)** Unsupervised clustering results of B cells and monocytes and their subsets based on Louvain algorithm.

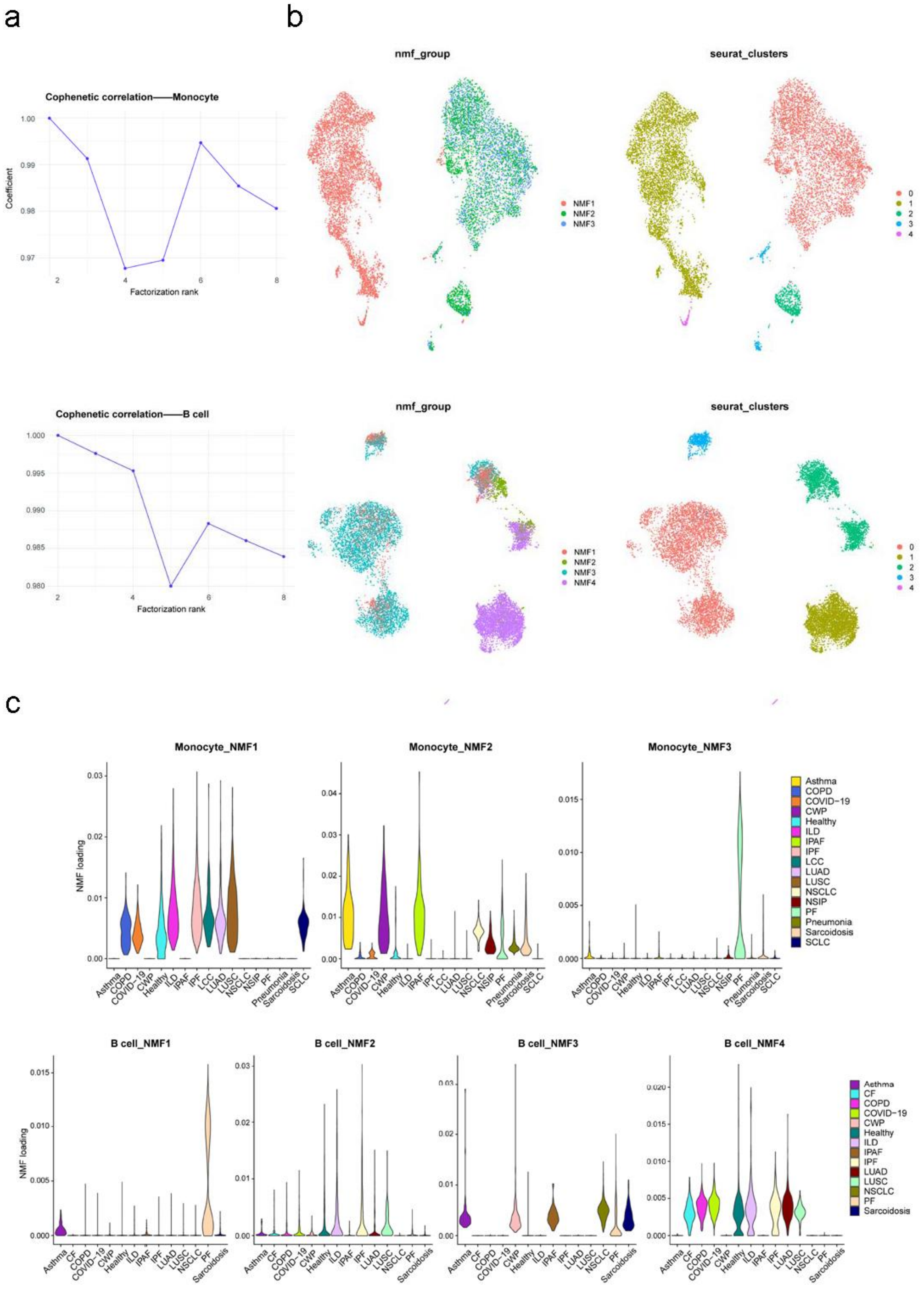


**Supplementary Figure 6: NMF analysis of B cells and monocytes.**

**(a)** Optimal k value estimation applied to NMF analysis of B cells and monocytes. **(b)** Clustering results of B cells and monocytes based on NMF algorithm. **(c)** Distribution of B cells and monocytes from different lung states at various factor levels.

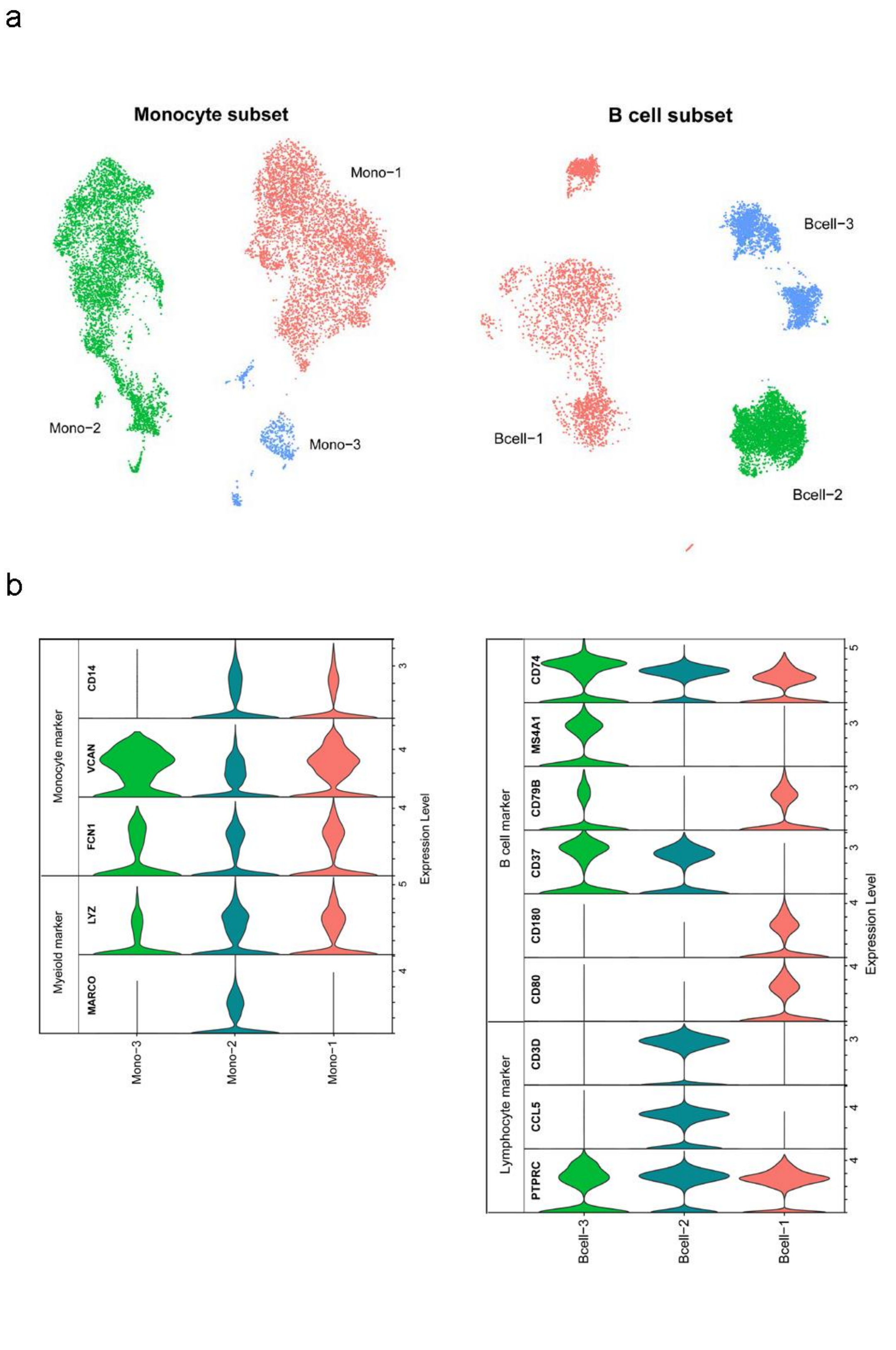


**Supplementary Figure 7: Identification of B cell and monocyte subpopulations.**

**(a)** Clustering results of B cells and monocytes. **(b)** Validation of expression levels of marker genes.

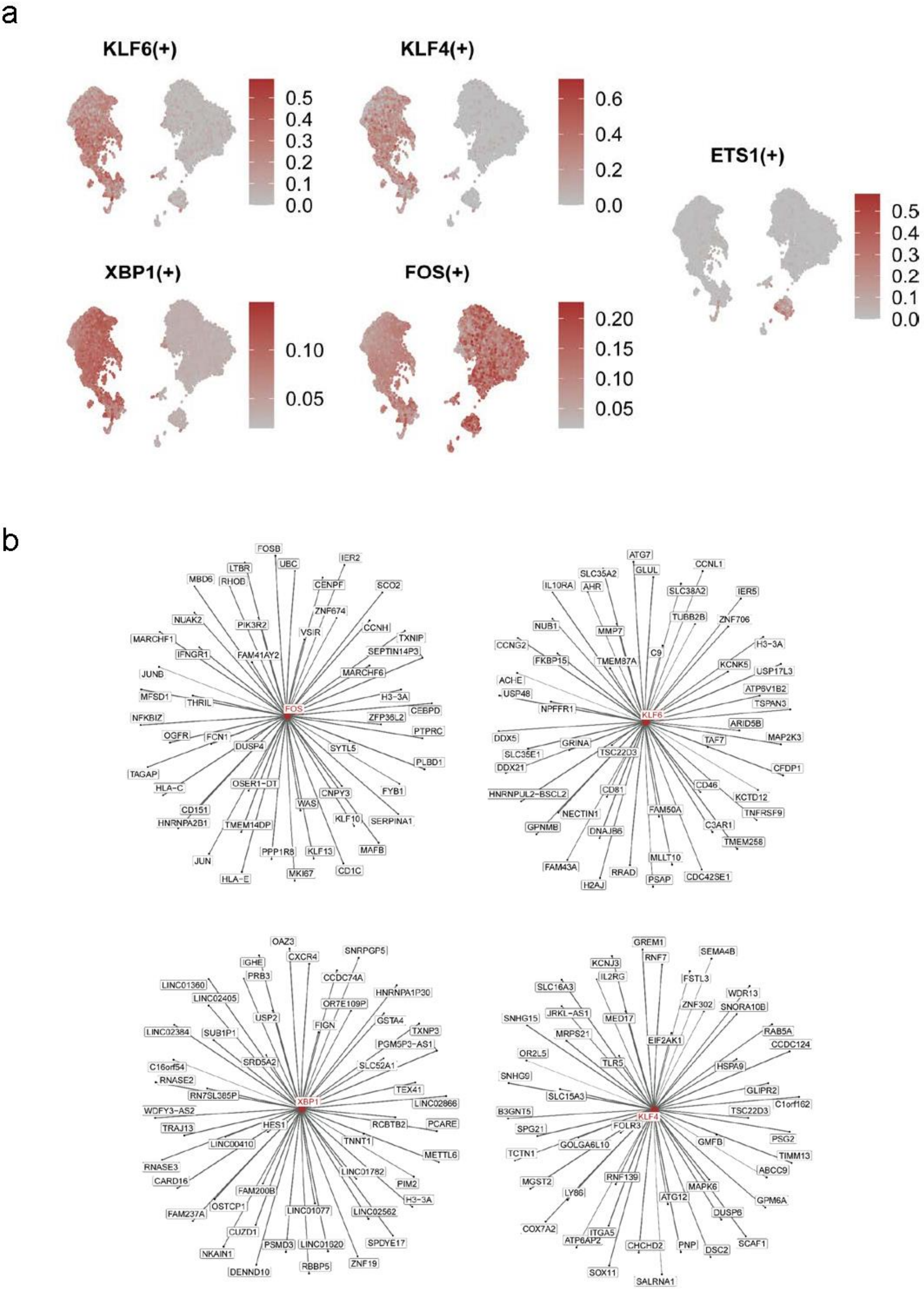


**Supplementary Figure 8: Transcription factor analysis of monocytes.**

**(a)** Active transcription factors in monocytes. **(b)** Gene regulation network centred on transcription factor.

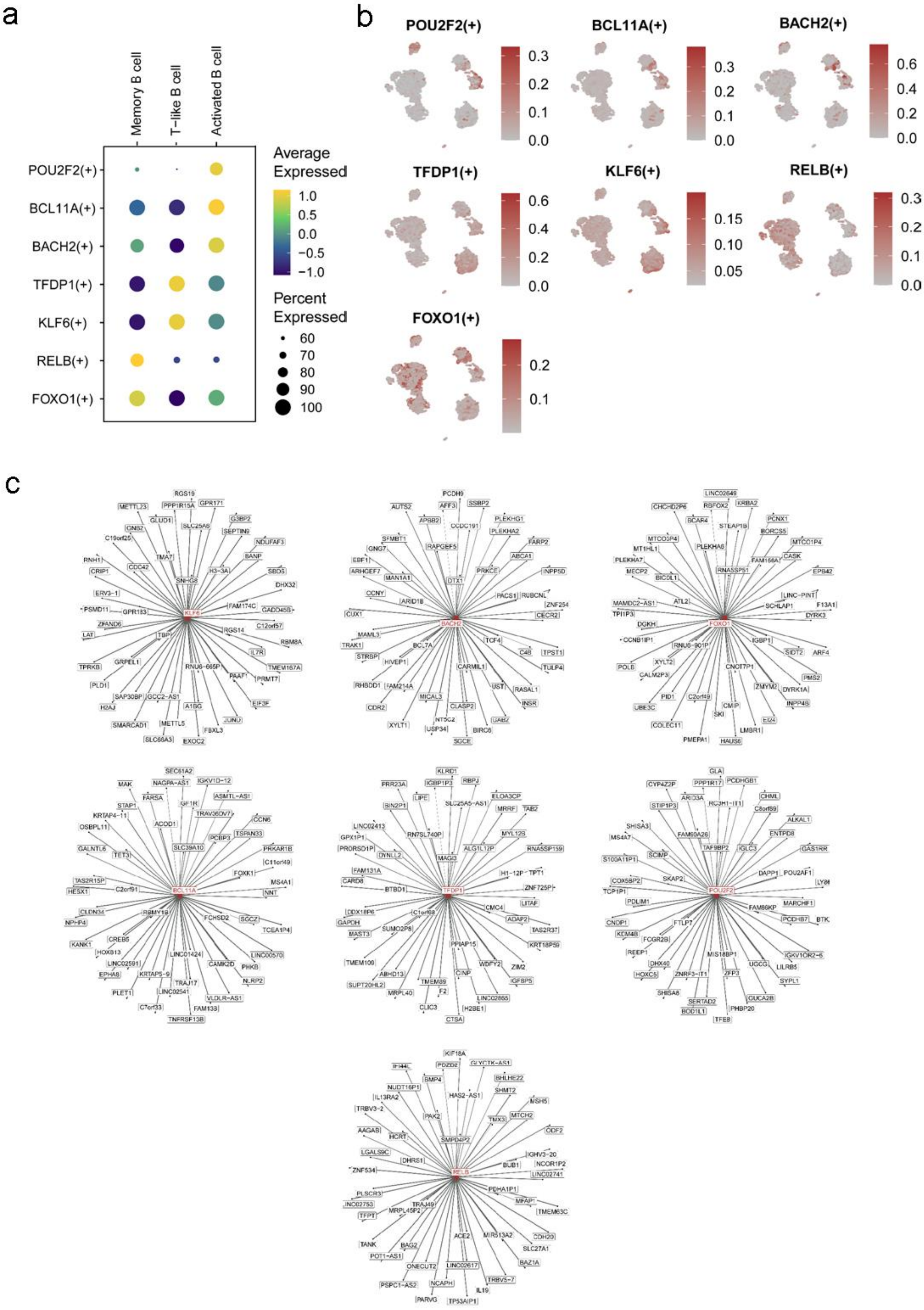


**Supplementary Figure 9: Transcription factor analysis of B cells.**

**(a-b)** Active transcription factors in different B cell subpopulations. **(c)** Gene regulation network centred on transcription factor.

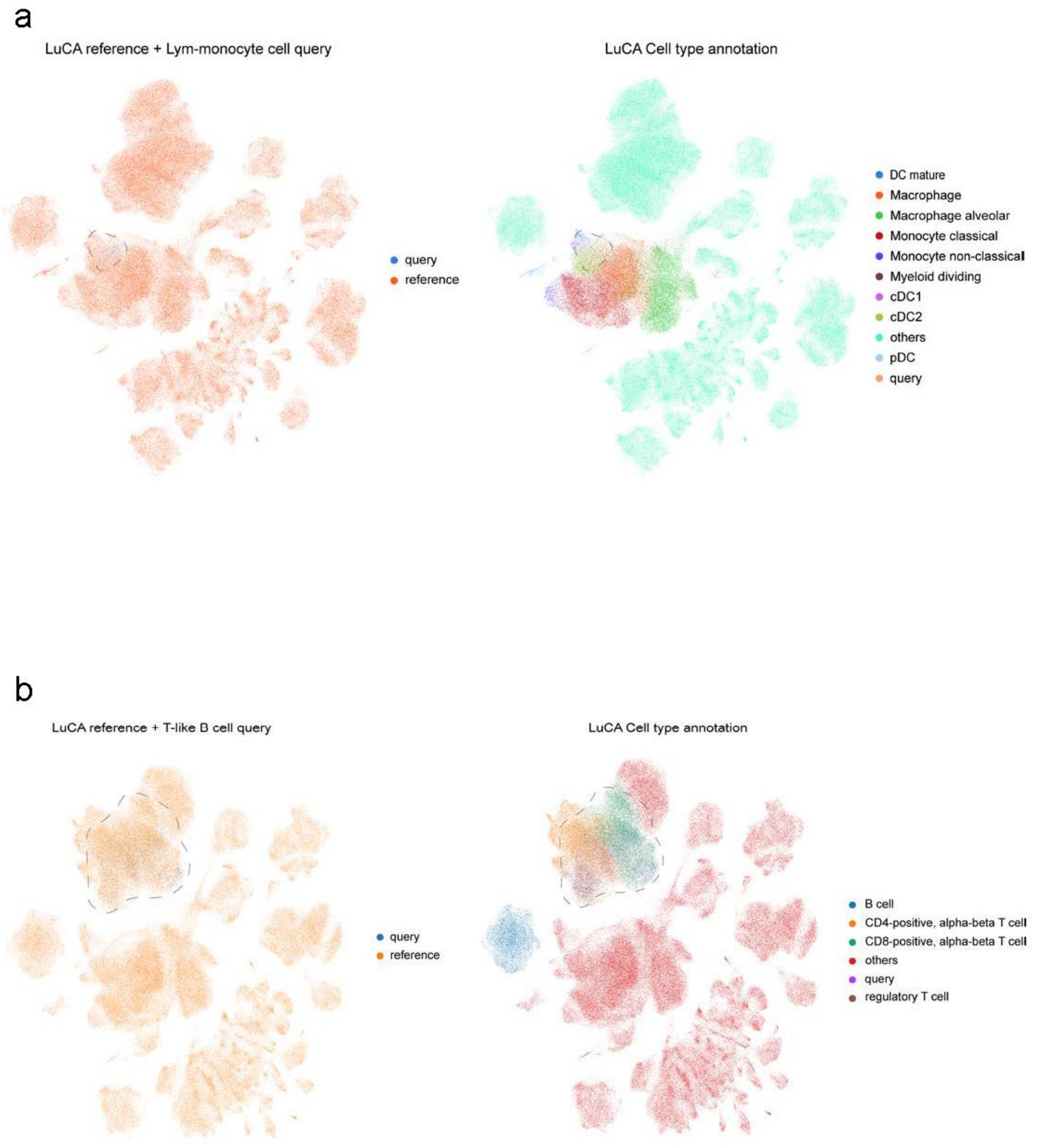


**Supplementary Figure 10: Mapping of cell subpopulations using LuCA as reference.**

**(a)** UMAP visualization of mapping Lym-monocyte to LuCA, highlighting the cell types surrounding the query data.

**(b)** UMAP visualization of mapping T-like B cell to LuCA, highlighting the cell types surrounding the query data.

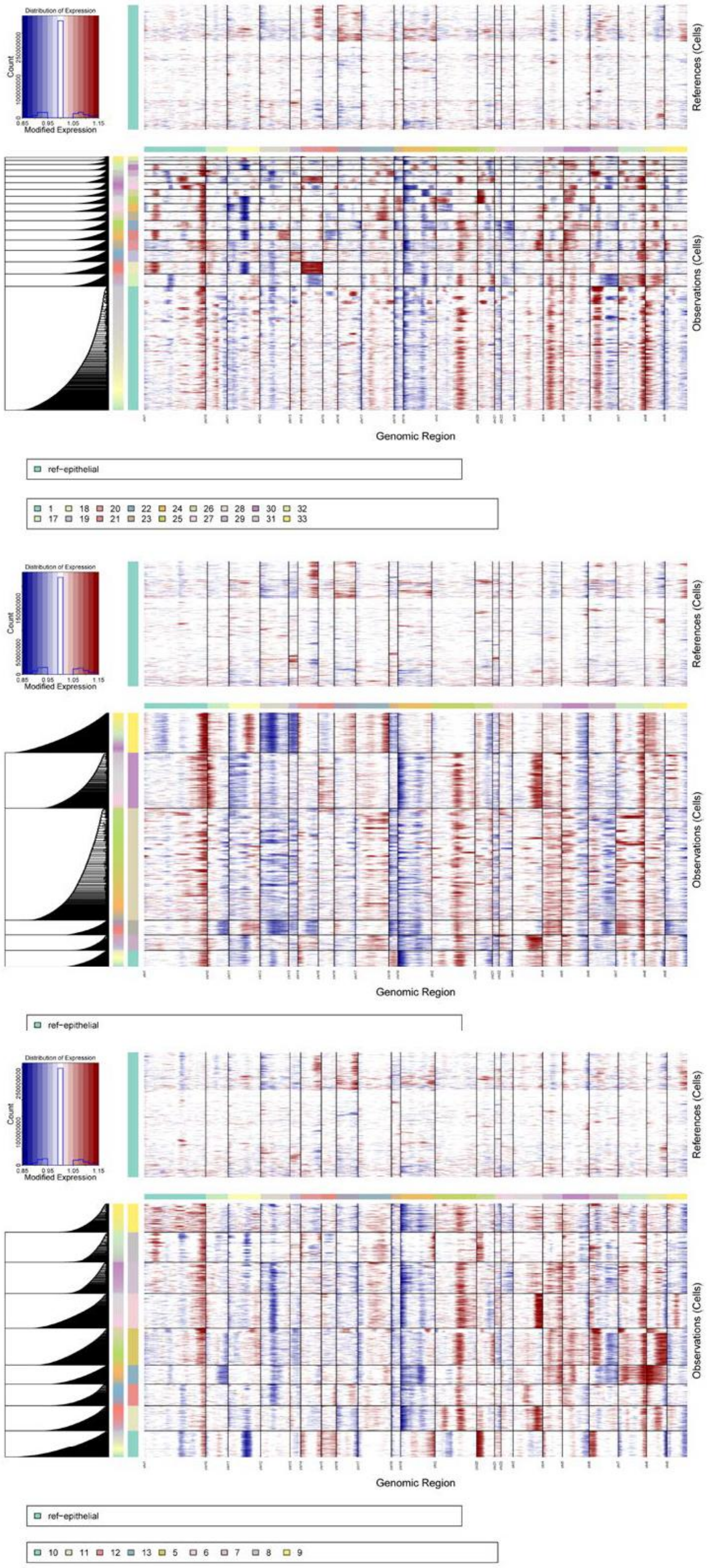


**Supplementary Figure 11: Estimation of copy number variation events in epithelial cells.** The inferCNV results for all epithelial cells were grouped based on clustering, with normal epithelial cells from healthy lungs used as a reference. Due to computational limitations, we divided cells into three cell subgroups and ran the analysis separately for each subpopulation.

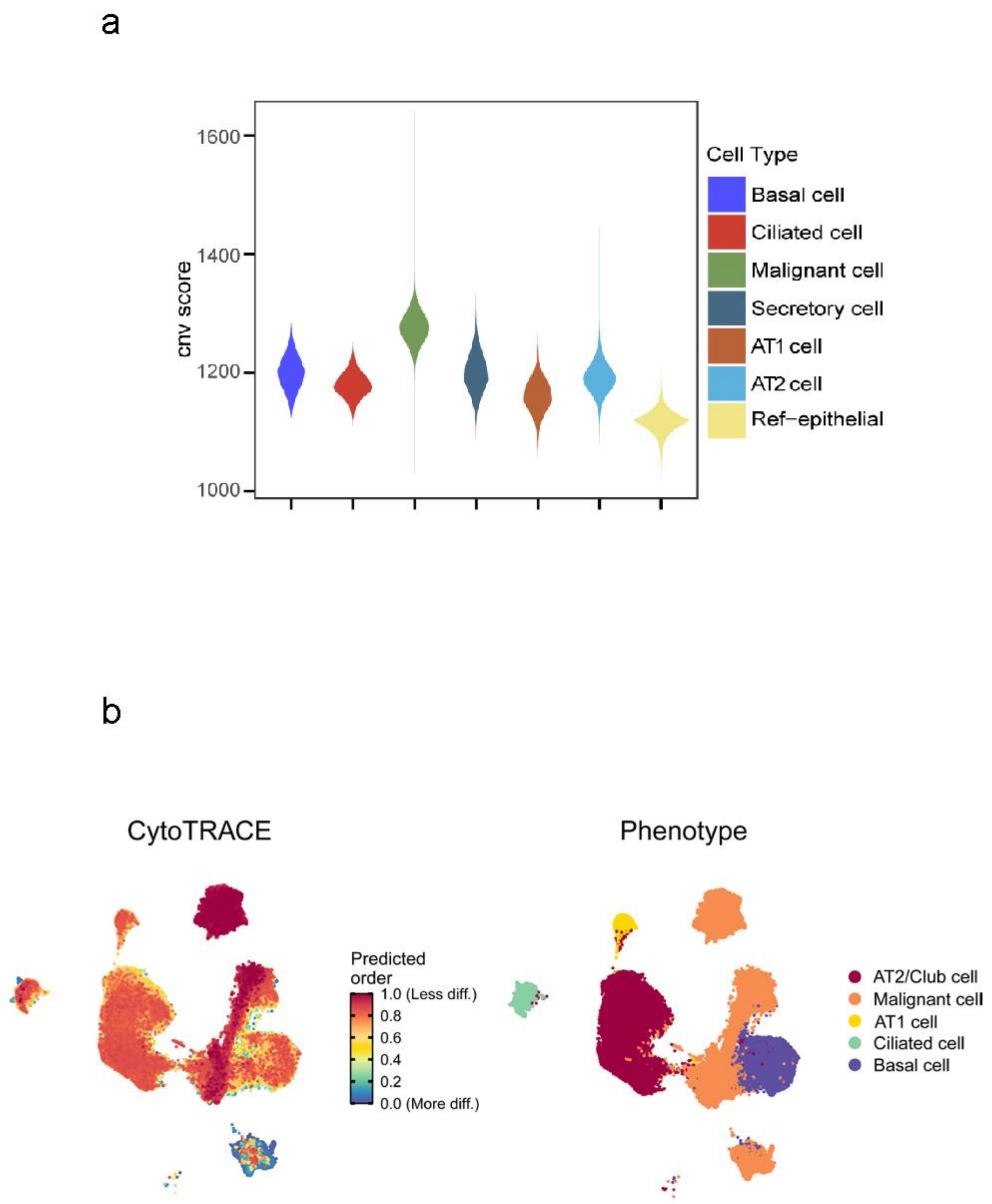


**Supplementary Figure 12: CNV score and differentiation potential estimation.**
**(a)** The inferCNV-derived results entail summing the copy number variation events across all chromosomal locations within each cell, thereby obtaining the CNV score for each cell subpopulation. **(b)** UMAP of all epithelial cells, colored by CytoTRACE-inferred differentiation potential

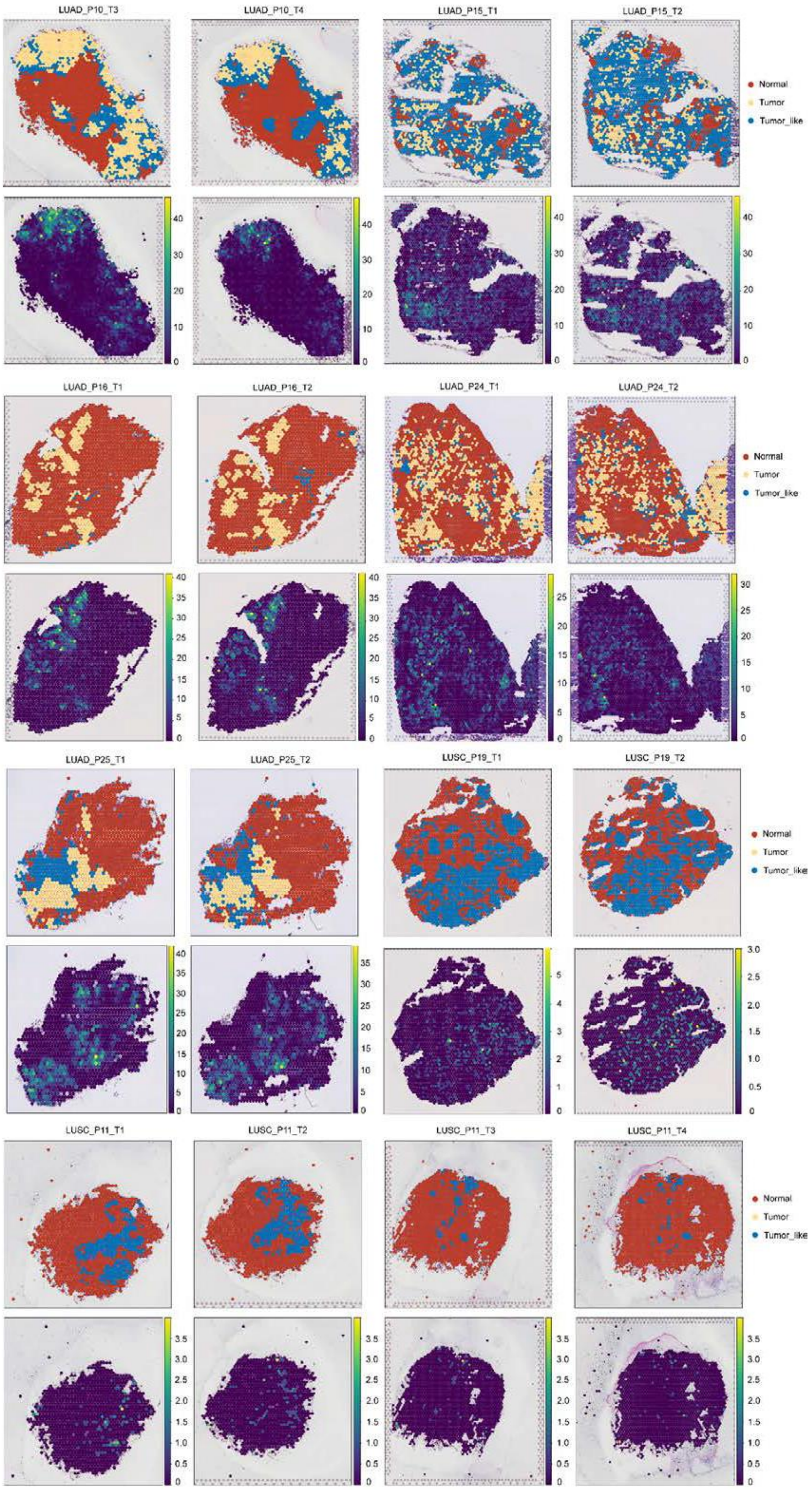


**Supplementary Figure 13: Spatial region division and EPCAM expression in each sample.** All samples were divided into normal, tumour cell and tumour-like cell regions, and the spatial expression distribution of EPCAM in each sample was examine.

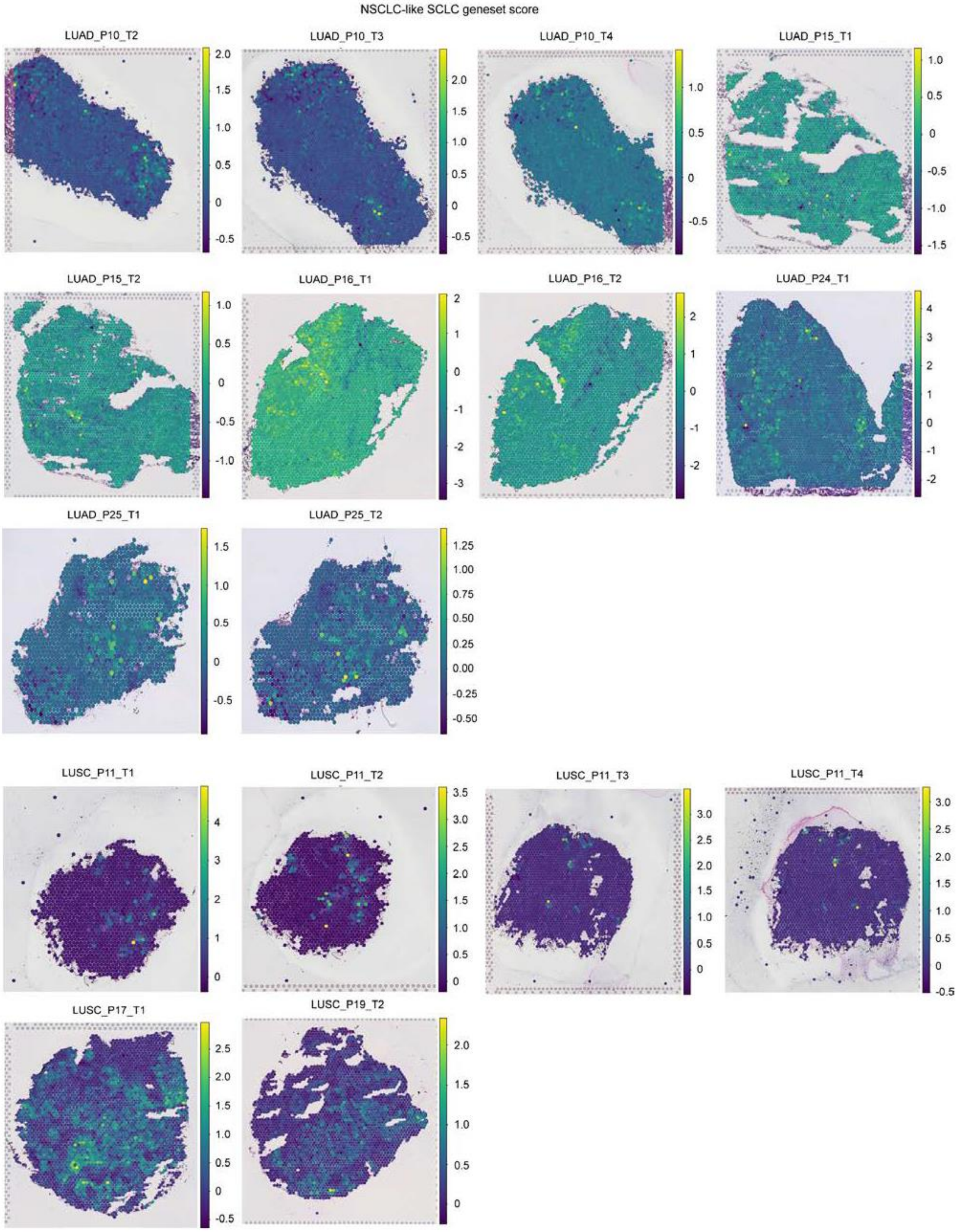


**Supplementary Figure 14: Scoring of NSCLC-like SCLC subset characteristic gene sets in LUAD and LUSC samples.** A gene-set score for the NSCLC-like SCLC subpopulation was performed in each sample, with the highlighted region showing the presence of the subpopulation.

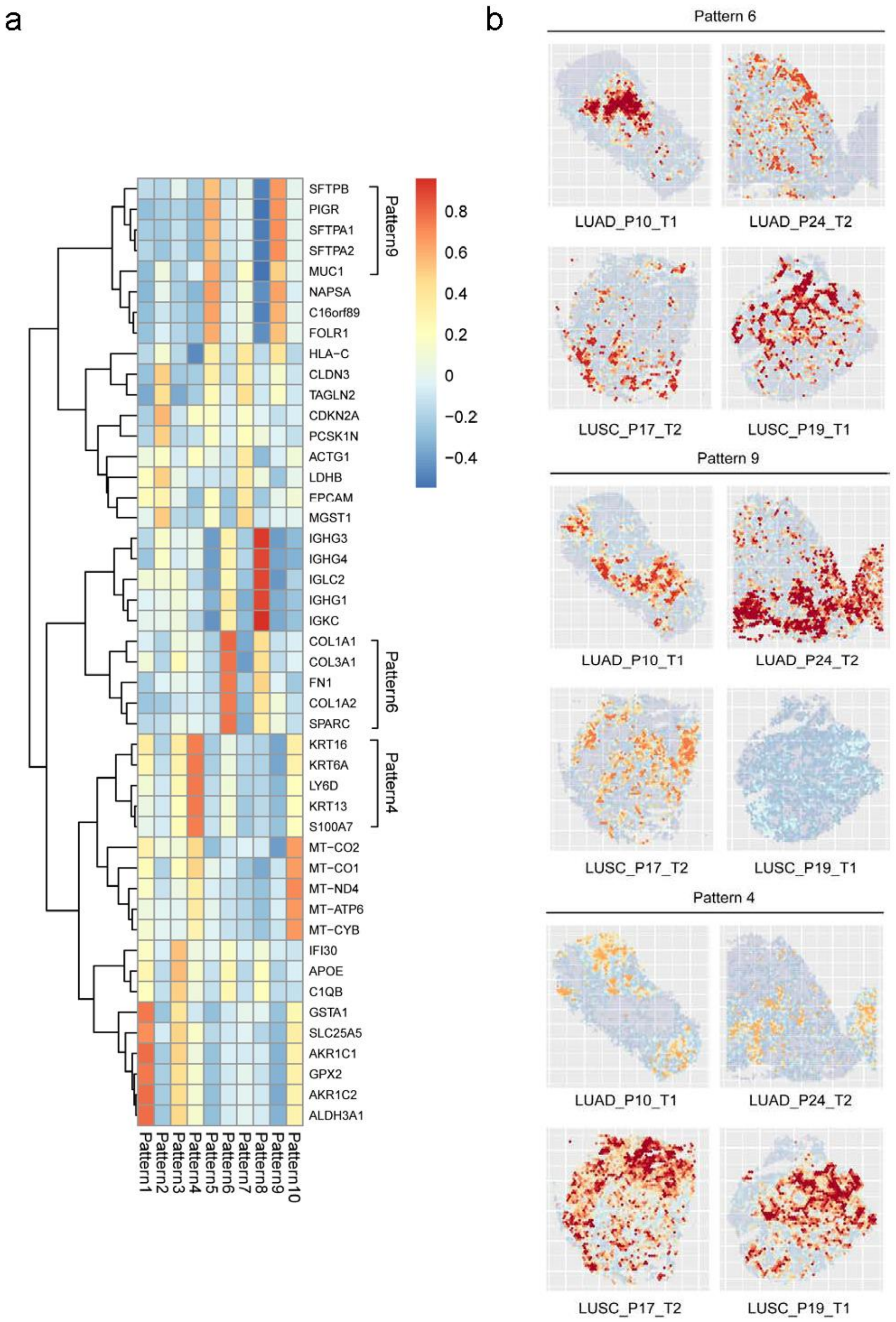


**Supplementary Figure 15: Identification of shared gene expression patterns in samples with high NSCLC-like SCLC features.**

**(a)** Shared gene expression patterns in samples with high NSCLC-like SCLC subpopulation feature scores. **(b)** In the samples with high NSCLC-like SCLC feature scores, there were gene expression patterns in all samples, LUAD samples and LUSC samples, respectively.

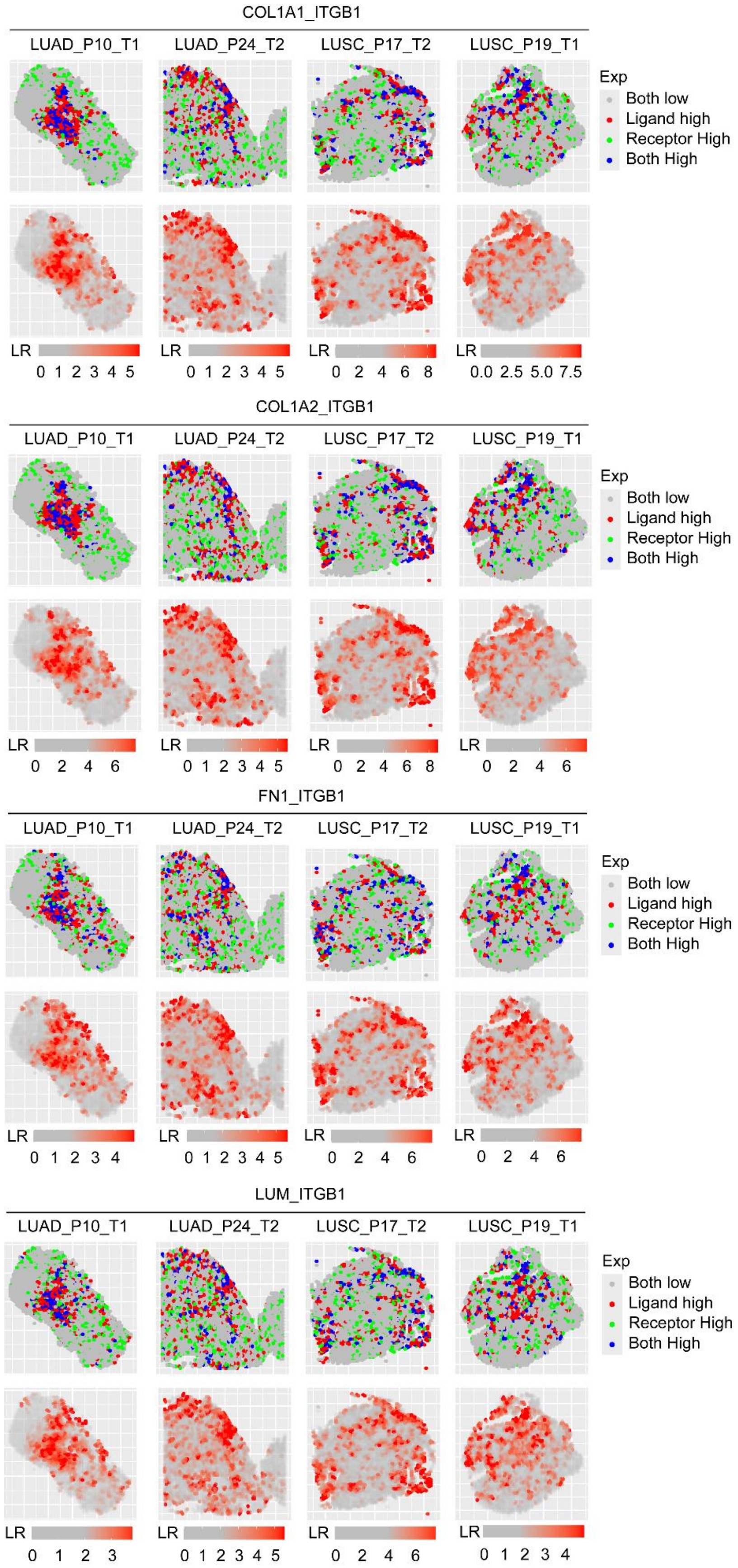


**Supplementary Figure 16: Identification of shared LRI in samples with high NSCLC-like SCLC features.** Concentrated activated LRI pairs in LUAD and LUSC samples with high NSCLC-like SCLC characteristics.

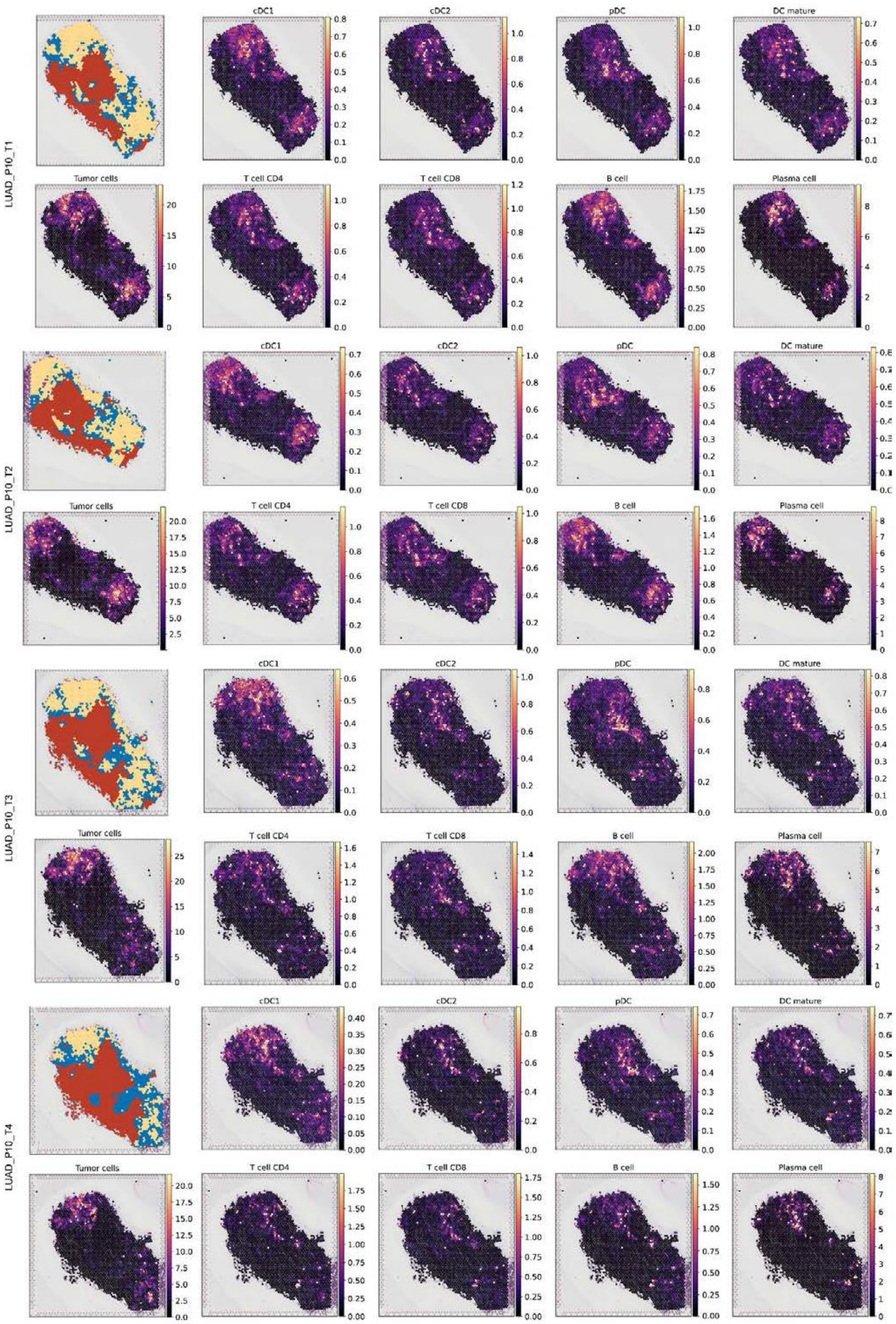


**Supplementary Figure 17: Cell type annotation of spatial transcriptome samples of LUAD-P10.** Using LuCA as reference, cell2location was used to annotate the spatial transcriptome samples and highlight the spatial distribution of immune cells.

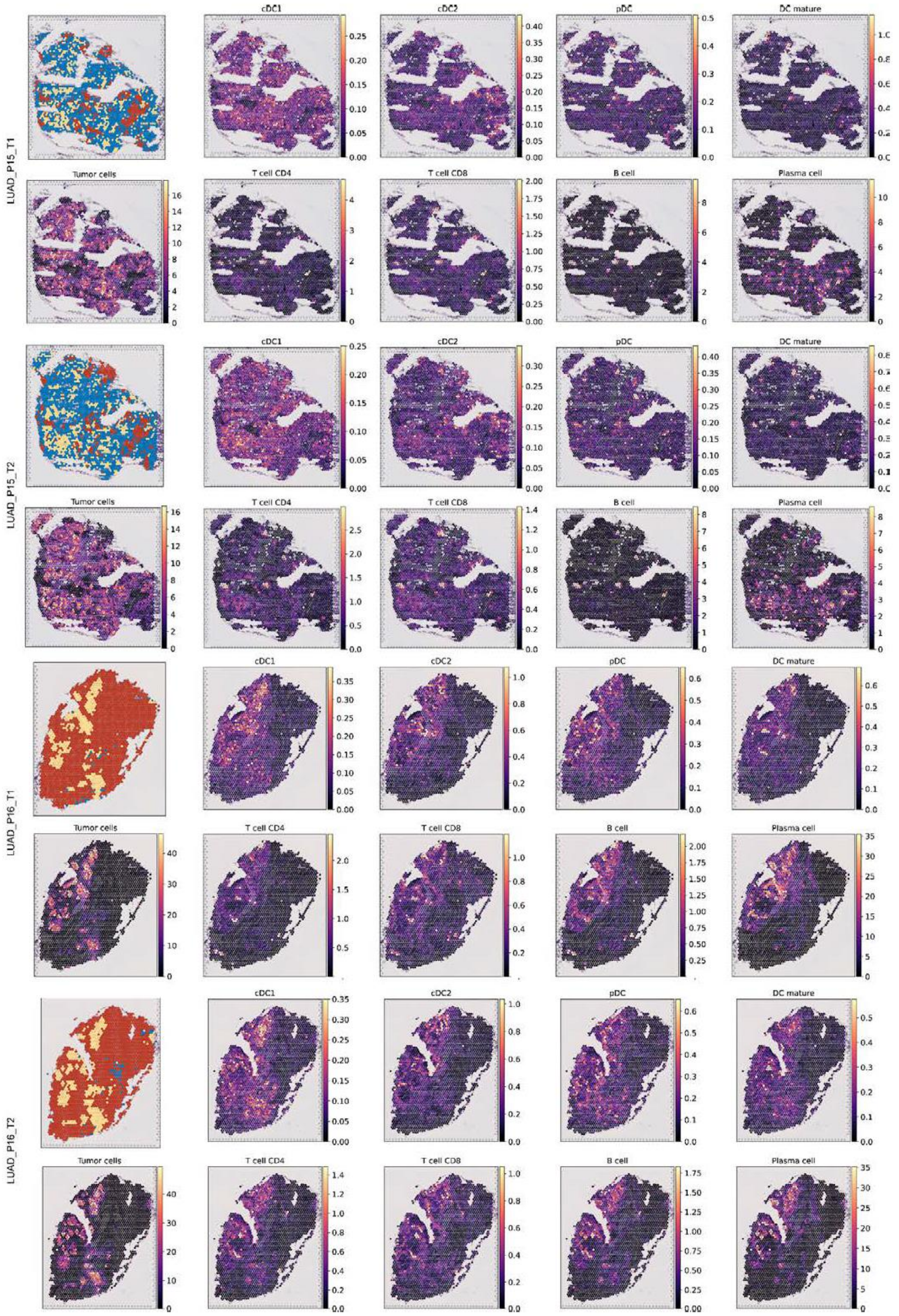


**Supplementary Figure 18: Cell type annotation of spatial transcriptome samples of LUAD-P15 and LUAD-P16.** Using LuCA as reference, cell2location was used to annotate the spatial transcriptome samples and highlight the spatial distribution of immune cells.

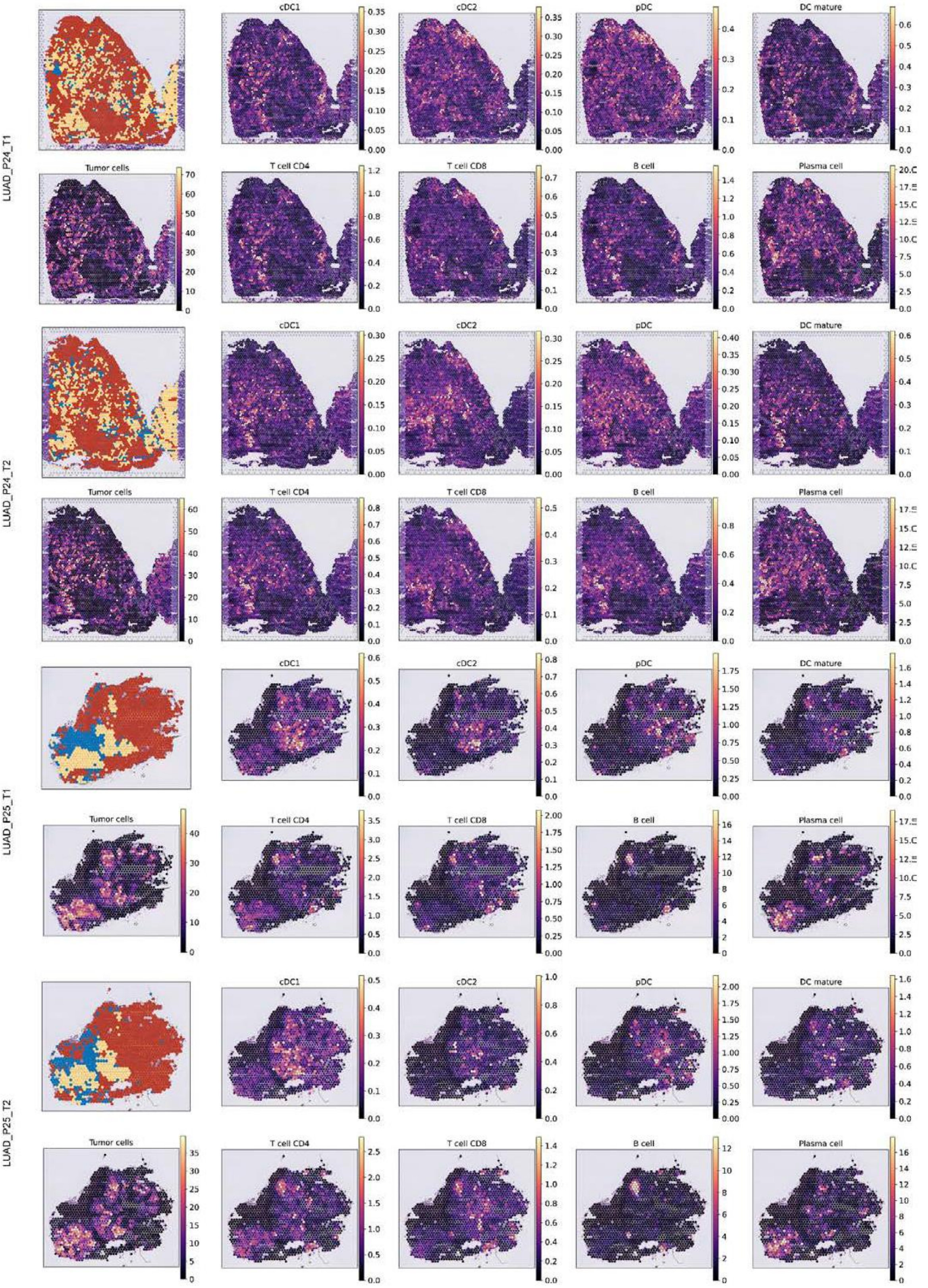


**Supplementary Figure 19: Cell type annotation of spatial transcriptome samples of LUAD-P24 and LUAD-P25.** Using LuCA as reference, cell2location was used to annotate the spatial transcriptome samples and highlight the spatial distribution of immune cells.

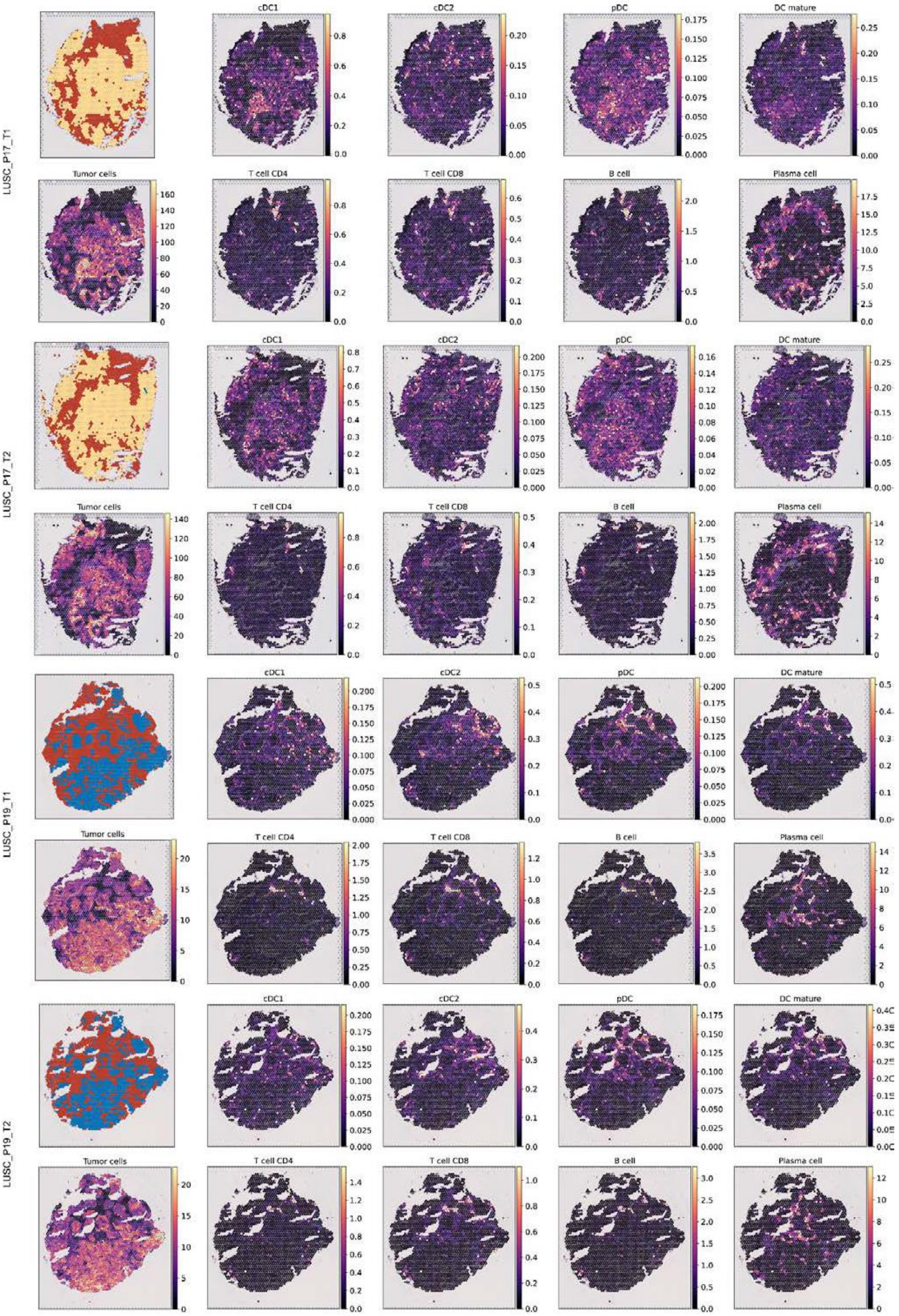


**Supplementary Figure 20: Cell type annotation of spatial transcriptome samples of LUSC-P17 and LUSC-P19.** Using LuCA as reference, cell2location was used to annotate the spatial transcriptome samples and highlight the spatial distribution of immune cells.

a

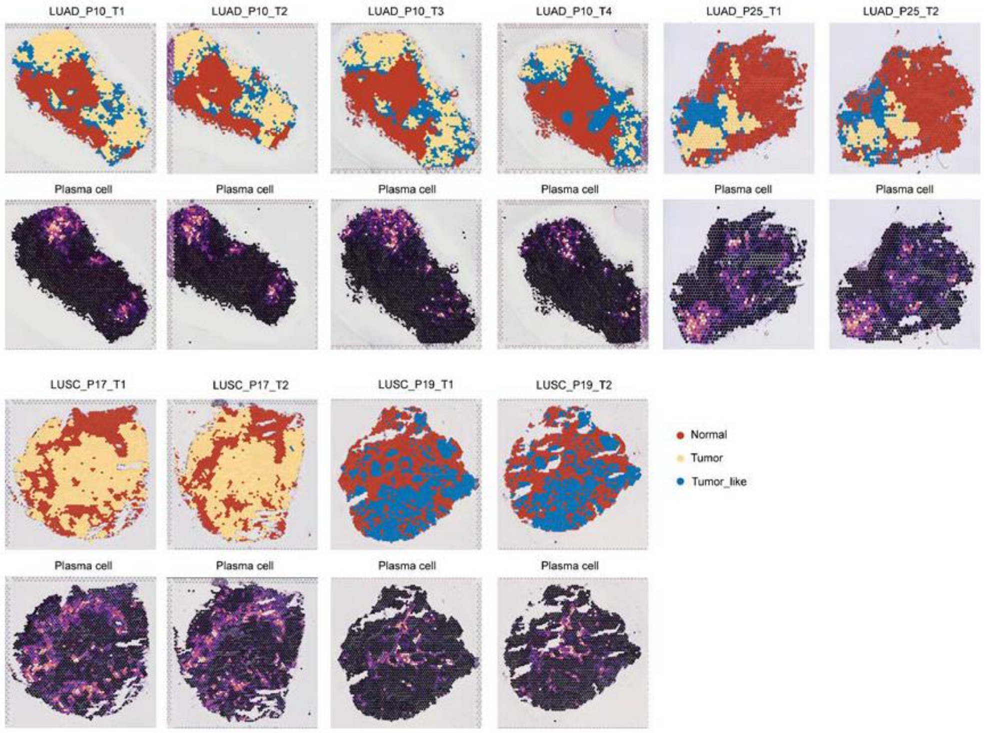


b

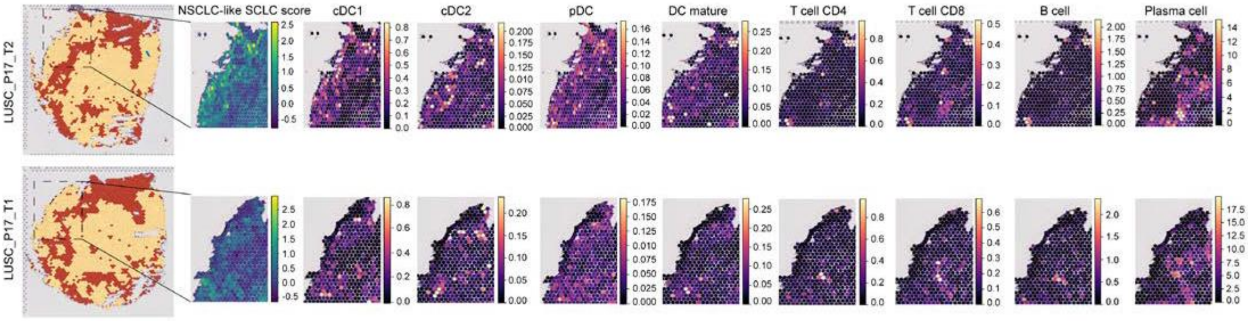


**Supplementary Figure 21: Differences in spatial distribution of immune cells.**

**(a)** In LUAD samples with high NSCLC-like SCLC features, plasma cells are concentrated in tumour cell region. In LUSC samples, plasma cells were distributed in normal areas around tumour cells. **(b)** Two tissue sections from the same LUSC patient exhibited varying levels of NSCLC-like SCLC features, with the sample showing prominent NSCLC-like SCLC features displaying only more pronounced dendritic cell infiltration in the tumour cell regions, excluding other lymphocytes.

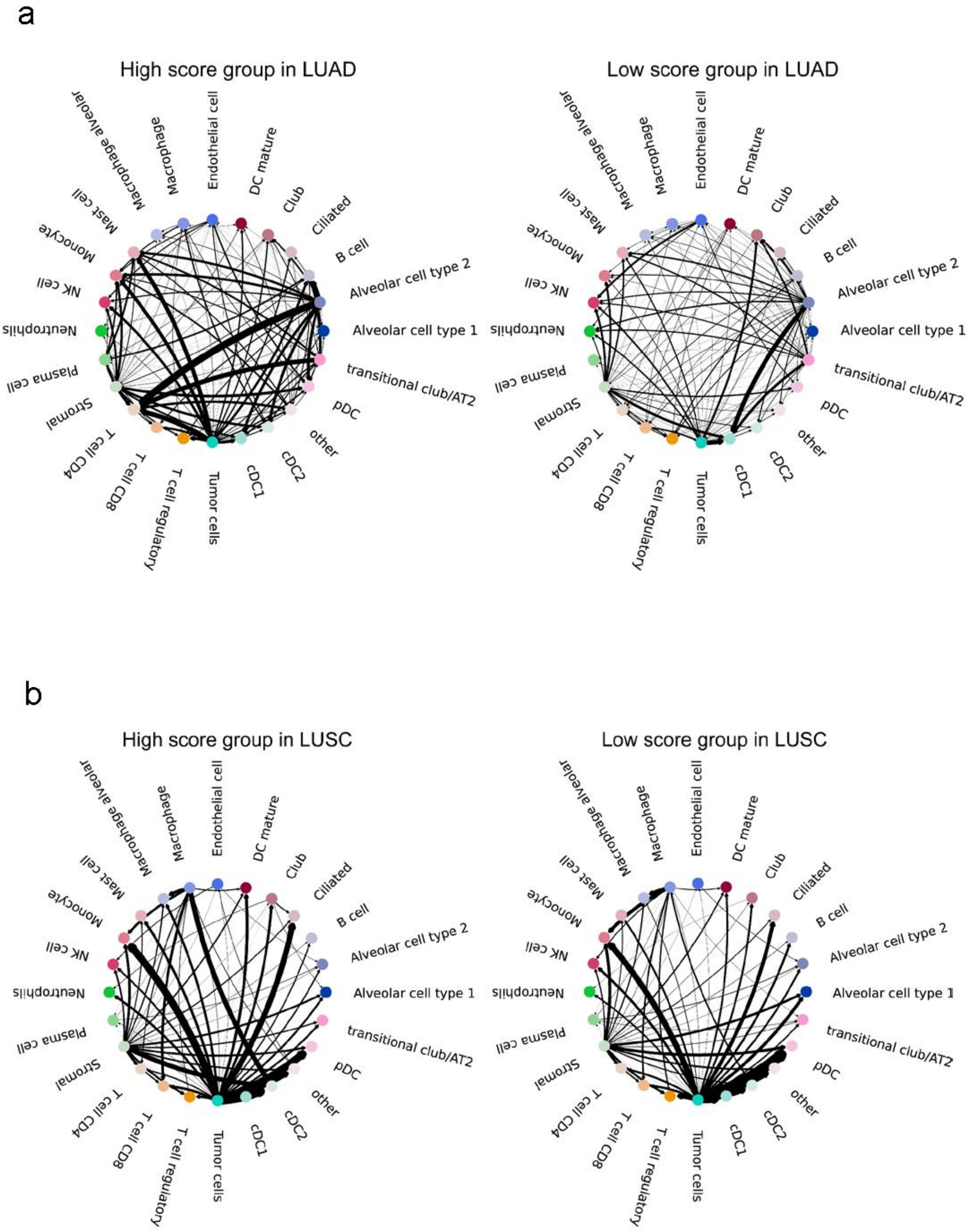


**Supplementary Figure 22: Differences in intercellular dependence between samples of LUAD and LUSC with high and low NSCLC-like SCLC feature scores.** Type coupling analysis with edge proportional to strength of directional dependencies by means of fold changes of differentially expressed genes for each pair of sender and receiver cell types between groups with different NSCLC-like SCLC score. The effect of the presence of NSCLC-like SCLC subpopulations in LUAD and LUSC on intercellular dependence was shown.

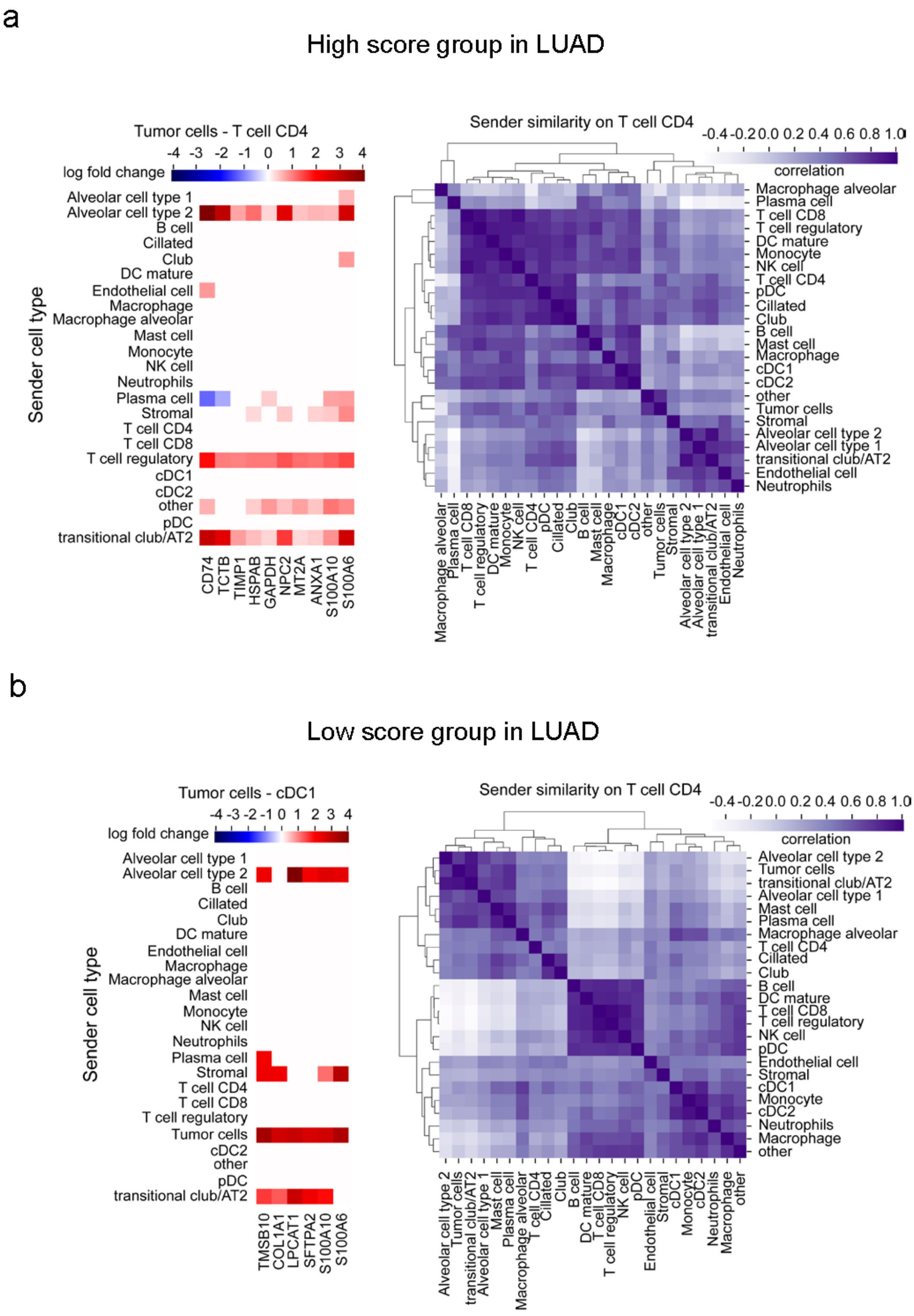


**Supplementary Figure 23: Sender effect analysis of LUAD samples with high and low NSCLC-like SCLC feature scores.** Sender effect analysis of the tumour cell-CD4 T cell axis in LUAD samples with high NSCLC-like SCLC scores and the tumour cell-cDC1 cell axis in LUAD samples with low NSCLC-like SCLC scores is shown. The y-axis represents the sender cell type, while the x-axis indicates the estimated fold change induced in the recipient cell genes. Additionally, sender similarity analysis is displayed based on the correlation between the coefficient vector of each sender cell type and the CD4 T cell receptor during the process of NSCLC-like SCLC feature changes.

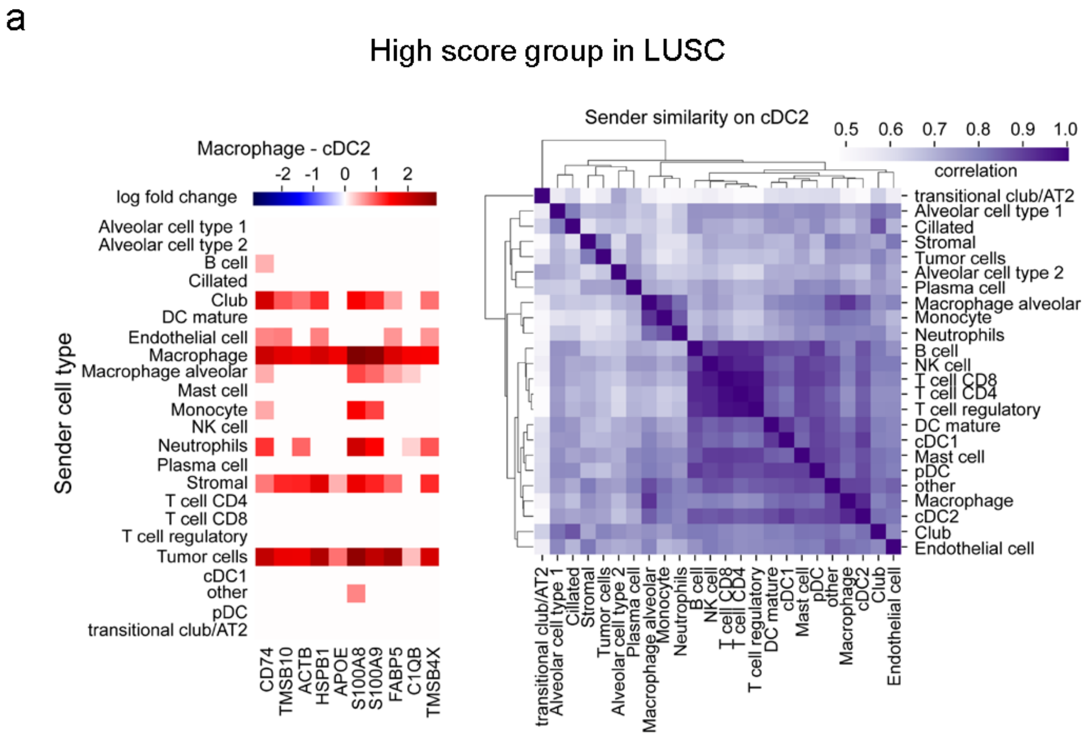


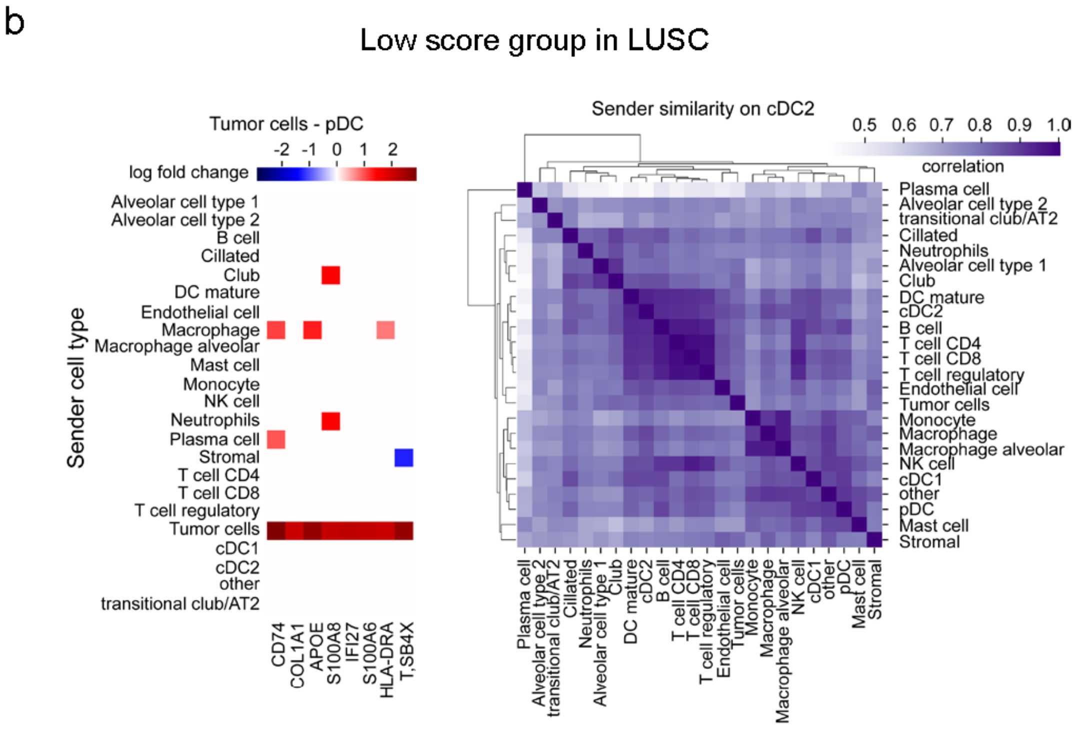


**Supplementary Figure 24: Sender effect analysis of LUSC samples with high and low NSCLC-like SCLC feature scores.** Sender effect analysis of the tumour cell-cDC2 cell axis in LUSC samples with high NSCLC-like SCLC scores and the tumour cell-pDC cell axis in LUSC samples with low NSCLC-like SCLC scores is shown. The y-axis represents the sender cell type, while the x-axis indicates the estimated fold change induced in the recipient cell genes. Additionally, sender similarity analysis is displayed based on the correlation between the coefficient vector of each sender cell type and the cDC2 cell receptor during the process of NSCLC-like SCLC feature changes.

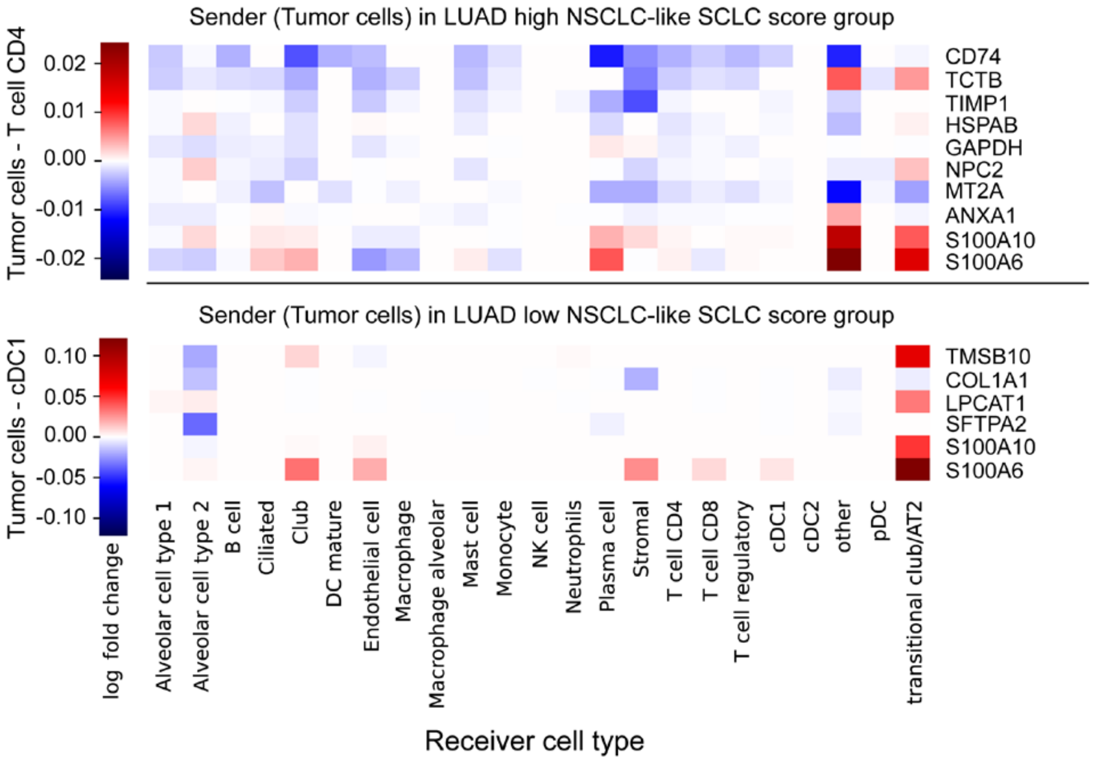


**Supplementary Figure 25: Receiver effect analysis of LUAD samples with high and low NSCLC-like SCLC feature scores.** Sender effect analysis of the tumour cell-cDC2 cell axis in LUAD samples with high NSCLC-like SCLC scores and the tumour cell-pDC cell axis in LUAD samples with low NSCLC-like SCLC scores is shown. The x-axis represents the receiver cell type, while the y-axis shows the estimated fold change induced in the sender cell (tumour cell) genes.
