## Supplementary Tables for "A universal single-cell transcriptomics atlas of human lung decodes multiple pulmonary diseases"

**Table S1. Details of uniLUNG core datasets.**

**Table S2. uHAF and references.**

**Table S3. Original cell label coordination by uHAF.**

**Table S4. Statistical of comparing the re-annotation of the uniLUNG core with the original annotation at Level 1.**

**Table S5. Statistical of comparing the re-annotation of the uniLUNG core with the original annotation at Level 2.**

**Table S6. Statistical of comparing the re-annotation of the uniLUNG core with the original annotation at Level 3.**

**Table S7. Statistical of comparing the re-annotation of the uniLUNG core with the original annotation at Level 4.**

**Table S8. Details of datasets in uniLUNG.**

**Table S9. Cell count statistics in lung multi-disease analysis.**

**Table S10. Donor information for lung cancer case studies.**

**Table S11. List of LRIs in LUAD and LUSC samples.**

**Table S12. List of active spatial signal pathways in LUAD samples based on COMMOT.**

**Table S13. List of active spatial signal pathways in LUSC samples based on COMMOT.**

**Table S14. Total 43878 gene list.**

**Table S1. Details of uniLUNG core datasets.**

| Sub-atlas | Reference | Region | Seq-tech | Expr value | Data accession | Lung status |
|---|---|---|---|---|---|---|
| Healthy | Madissoon E et al. 2019. Genome Biology,10.1186/s13059-019-1906-x | Lung | 10X | read counts | PRJEB31843 | Normal |
| Healthy | Han X et al. 2020. Nature, 10.1038/s41586-020-2157-4 | Lung | Microwell-seq | read counts | GSE134355 | Normal |
| Healthy | Holloway Emily M et al. 2020. Developmental Cell, 10.1016/j.devcel.2020.07.023 | Bronchi | 10X | read counts | E-MTAB-8221 | Normal |
| Healthy | Lukassen et al. 2020. The EMBO Journal, 10.15252/embj.20105114 | Bronchi | 10X | read counts | MendeleyData7r2cwbw44m | Normal |
| Healthy | | Lung | 10X | read counts | MendeleyData7r2cwbw44m | Normal |
| Healthy | Deprez M et al. 2020. American journal of respiratory and critical care medicine,10.1164/rccm.201911-2199OC | Lung | 10X | read counts | EGAS00001004082 | Normal |

| | | | | | | |
|---|---|---|---|---|---|---|
| Healthy | Goldfarbmuren et al. 2020. Nat Commun, 10.1038/s41467-020-16239-z | Bronchi | 10X | read counts | GSE134174 | Normal |
| Healthy | Vieira Braga FA et al. 2019. Nat Med, 10.1038/s41591-019-0468-5 | Parenchyma | Drop-seq | read counts | GSE130148 | Normal |
| Healthy | | Lung | 10X | read counts | GSE173896 | Normal |
| Healthy | Caushi JX et al. 2021. Nature. 10.1038/s41586-021-03752-4 | Lung | 10X | read counts | GSE176021 | Normal |
| Healthy | Li X et al. 2022. Life Sci Alliance. 10.26508/lsa.202201458 | Alveolus | 10X | read counts | GSE193782 | Normal |
| Healthy | Wang C et al. 2023. Immunity. 10.1016/j.immuni.2023.01.032 | Lung | 10X | read counts | GSE196638 | Normal |
| Healthy | Speir ML et al. 2021. Bioinformatics. 10.1093/bioinformatics/btab503 | Lung | 10X | read counts | GTEx V9 | Normal |
| Healthy | Salcher S et al. 2022. Cancer Cell. 10.1016/j.ccell.2022.10.008 | Lung | 10X | read counts | LuCA | Normal |

| | | | | | | |
|---|---|---|---|---|---|---|
| Healthy | L Sikkema et al. 2022. bioRxiv https://doi.org/10.1101/2022.03.10.483747 | Lung | 10X | read counts | HLCA | Normal |
| Healthy | Chan JM et al. 2021. Cancer Cell. 10.1016/j.ccell.2021.09.008 | Lung | 10X | read counts | HTAN | Normal |
| Healthy | | Lung | 10X | read counts | HuBMAP | Normal |
| Healthy | Raredon MSB et al. 2019. Sci Adv. 10.1126/sciadv.aaw3851 | Lung | Drop-seq | read counts | GSE133747 | Normal |
| Healthy | Vieira Braga FA et al. 2019. Nat Med, 10.1038/s41591-019-0468-5 | Lung | Smartseq2 | read counts | GSE148829 | Normal |
| Healthy | Liao M et al. 2020. Nat Med. 10.1038/s41591-020-0901-9 | Bronchi | 10X | read counts | GSE145926 | Normal |
| Healthy | Tsukui T et al. 2020. Nat Commun, 10.1038/s41467-020-15647-5 | Lung | 10X | read counts | GSE132771 | Normal |
| Healthy | Reyfman PA et al. 2019. Am J Respir Crit Care Med, 10.1164/rccm.201712-2410OC | Lung | 10X | read counts | GSE122960 | Normal |

| | | | | | | |
|---|---|---|---|---|---|---|
| Healthy | Szabo PA et al. 2019. Nat Commun, 10.1038/s41467-019-12464-3 | Parenchyma | 10X | read counts | GSE126030 | Normal |
| Healthy | Morse C et al. 2019. Eur Respir J, 10.1183/13993003.02441-2018 | Bronchi | 10X | read counts | GSE128033 | Normal |
| Healthy | Travaglini et al. 2020. Nature, 10.1038/s41586-020-2922-4 | Parenchyma | 10X | read counts | EGAS00001004344 | Normal |
| Healthy | | Parenchyma | Smart-seq | read counts | EGAS00001004344 | Normal |
| Healthy | Domínguez Conde C et al. 2022. Science, 10.1126/science.abl5197 | Parenchyma | Smartseq2 | read counts | E-MTAB-11536 | Normal |
| Healthy | Wei Kheng The et al. 2022. BioStudies | Lung | 10X | read counts | E-MTAB-11267 | Normal |
| Healthy | Okuda K et al. 2021. Am J Respir Crit Care Med, 10.1164/rccm.202008-3198OC. | Bronchi | 10X | read counts | GSE160664 | Normal |
| Healthy | | Bronchi | Drop-seq | read counts | GSE160673 | Normal |

| | | | | | | |
|---|---|---|---|---|---|---|
| Healthy | Miller AJ et al. 2020. Cell, 10.1016/j.devcel.2020.01.033. | Bronchi, Alveolus | 10X | read counts | E-MTAB-8221 | Normal |
| Healthy | | Lung | 10X | read counts | GSE121080 | Normal |
| Healthy | Habermann AC et al. 2020. Sci Adv, 10.1126/sciadv.aba1972 | Lung | 10X | read counts | GSE135893 | Normal |
| Healthy | Jaeger B et al. 2022. Nat Commun, 10.1038/s41467-022-33193-0 | Bronchi | 10X | read counts | GSE141939 | Normal |
| Healthy | Okuda, Kenichi et al. 2021. American journal of respiratory and critical care medicine, 10.1164/rccm.202008-3198OC | Bronchi, Alveolus | Drop-seq | read counts | GSE160673 | Normal |
| Healthy | | Bronchi, Alveolus | 10X | read counts | GSE160794 | Normal |
| Healthy | Okuda K et al. 2021. Am J Respir Crit Care Med. 10.1164/rccm.202008-3198OC | Bronchi | 10X | read counts | GSE160664 | Normal |
| Healthy | Wang A et al. 2020. Elife. 10.7554/eLife.62522 | Lung | 10X | read counts | GSE161382 | Normal |

| | | | | | | |
|---|---|---|---|---|---|---|
| Healthy | Yuan Y et al. 2021. Front Bioeng Biotechnol, 10.3389/fbioe.2021.760309 | Parenchyma | 10X | read counts | GSE164829 | Normal |
| Healthy | de Rooij LPMH et al. 2022. Cardiovasc Res, 10.1093/cvr/cvac139 | Parenchyma | 10X | read counts | GSE159585 | Normal |
| Healthy | Pisu D et al. 2021. J Exp Med, 10.1084/jem.20210615 | Broncho, Alveolus | 10X | read counts | GSE167232 | Normal |
| Immune | Domínguez Conde, C et al. 2022. Science, 10.1126/science.abl5197 | Parenchyma, PBMC | 10X | read counts | E-MTAB-11536 | Normal |
| Immune | Liu, Can et al. 2021. Cell, 10.1016/j.cell.2021.02.018 | PBMC | 10X | read counts | GSE161918 | Normal |
| Immune | Szabo, Peter A et al. 2019. Nature communications, 10.1038/s41467-019-12464-3 | Parenchyma, PBMC | 10X | read counts | GSE126030 | Normal |
| Immune | Chan, Joseph M et al. 2020. bioRxiv, 10.1101/2020.12.01.406363 | Parenchyma | 10X | read counts | cellxgene/62e8f058-9c37-48bc-9200-e767f318a8ec | Normal, SCLC, LUAD |

| | | | | | | |
|---|---|---|---|---|---|---|
| Immune | Salcher, Stefan et al. 2022. Cancer cell, 10.1016/j.ccell.2022.10.008 | Lung | 10X Smartseq2 DropSeq InDrop | read counts | cellxgene/edb893ee-4066-4128-9aec-5eb2b03f8287 | Normal, Cancer |
| Cancer | Prazanowska KH, Lim SB. 2023. Sci data, 10.1038/s41597-023-02074-6 | Lung | 10X | read counts | GSE131907 GSE136246 GSE148071 GSE127465 | Cancer |
| Cancer | Salcher, Stefan et al. 2022. Cancer cell, 10.1016/j.ccell.2022.10.008 | Lung | 10X Smartseq2 InDrop BD Rhapsody Singleron | read counts | cellxgene/edb893ee-4066-4128-9aec-5eb2b03f8287 | Cancer |
| Cancer | Zhang, Li et al. 2022. Signal transduction and targeted therapy, 10.1038/s41392-021-00824-9 | Lung | 10X | read counts | Lungcancer_chenlulab | Cancer |

| | | | | | | |
|---|---|---|---|---|---|---|
| Cancer | Chan, Joseph M et al. 2021. Cancer cell, 10.1016/j.ccell.2021.09.008 | Lung | 10X | read counts | human tumor atlas | Cancer |
| Cancer | Song, Qianqian et al. 2019. Cancer medicine, 10.1002/cam4.2113 | Lung | 10X | read counts | GSE117570 | Cancer |
| Covid-19 | Liao, Mingfeng et al. 2020. Nature medicine, 10.1038/s41591-020-0901-9 | Bronchi | 10X | read counts | GSE145926 | COVID-19 |
| Covid-19 | Lee, Jeong Seok et al. 2020. Science immunology, 10.1126/sciimmunol.abd1554 | PBMC | 10X | read counts | GSE149689 | COVID-19 |
| Covid-19 | Xu, Gang et al. 2020. Clinical and translational medicine, 10.1002/ctm2.224 | Bronchi | 10X | read counts | GSE149878 | COVID-19 |
| Covid-19 | Wilk, Aaron J et al. 2020. Nature medicine, 10.1038/s41591-020-0944-y | PBMC | 10X | read counts | GSE150728 | COVID-19 |
| Covid-19 | Yao, Changfu et al. 2021. Cell Reports, 10.1016/j.celrep.2020.108590 | PBMC | 10X | read counts | GSE154567 | COVID-19 |
| Covid-19 | Grant, R.A. et al. 2021. Nature, 10.1038/s41586-020-03148-w | Bronchi | 10X | read counts | GSE155249 | COVID-19 |

| | | | | | | |
|---|---|---|---|---|---|---|
| Covid-19 | Wang, Allen et al. 2020. eLife, 10.7554/eLife.62522 | PBMC | 10X | read counts | GSE161382 | COVID-19 |
| Covid-19 | Bernardes, Joana P et al. 2020. Immunity, 10.1016/j.immuni.2020.11.017 | PBMC | 10X | read counts | GSE161777 | COVID-19 |
| Covid-19 | Bacher, Petra et al. 2020. Immunity, 10.1016/j.immuni.2020.11.016 | PBMC | 10X | read counts | GSE162086 | COVID-19 |
| Covid-19 | Heming, Michael et al. 2021. Immunity, 10.1016/j.immuni.2020.12.011 | PBMC | 10X | read counts | GSE163005 | COVID-19 |
| Covid-19 | Combes, Alexis J et al. 2021. Nature, 10.1038/s41586-021-03234-7 | PBMC | 10X | read counts | GSE163668 | COVID-19 |
| Covid-19 | Melms, Johannes C et al. 2021. Nature, 10.1038/s41586-021-03569-1 | Parenchyma | 10X | read counts | GSE171524 | COVID-19 |
| Covid-19 | Georg, Philipp et al. 2022. Cell, 10.1016/j.cell.2021.12.040 | PBMC | 10X | read counts | GSE175450 | COVID-19 |
| Covid-19 | Choi, Baekgyu et al. 2022. Experimental & molecular medicine, 10.1038/s12276-022-00866-1 | PBMC | 10X | read counts | GSE182123 | COVID-19 |

| | | | | | | |
|---|---|---|---|---|---|---|
| Covid-19 | Khoo, Weng Hua et al. 2023. Clinical immunology (Orlando, Fla.), 10.1016/j.clim.2022.109209 | PBMC | 10X | read counts | GSE196456 | COVID-19 |
| Covid-19 | Lee, Hye Kyung et al. 2022. iScience, 10.1016/j.isci.2022.104473 | PBMC | 10X | read counts | GSE201535 | COVID-19 |
| Covid-19 | Iwamura, Chiaki et al. 2022. Proceedings of the National Academy of Sciences of the United States of America, 10.1073/pnas.2203437119 | PBMC | 10X | read counts | GSE208337 | COVID-19 |
| Covid-19 | Santer, Deanna M et al. 2022. Nature communications, 10.1038/s41467-022-34709-4 | PBMC | 10X | read counts | GSE215814 | COVID-19 |
| Covid-19 | Xu, Jintao et al. 2022. Frontiers in immunology, 10.3389/fimmu.2022.970287 | PBMC | 10X | read counts | GSE216020 | COVID-19 |
| Covid-19 | Ren, Xianwen et al. 2021. Cell, 10.1016/j.cell.2021.01.053 | PBMC | 10X | read counts | GSE158055 | COVID-19 |
| Dev&Aging | Cao, Junyue et al. 2020. Science, 10.1126/science.aba7721 | Lung | 10X | read counts | descartes | Normal |

| | | | | | | |
|---|---|---|---|---|---|---|
| Dev&Aging | Zepp, Jarod A et al. 2021. Science, 10.1126/science.abc3172 | Lung | 10X | read counts | GSE149563 | Normal |
| Dev&Aging | Sountoulidis, Alexandros et al. 2023. Nature cell biology, 10.1038/s41556-022-01064-x | Lung | 10X | read counts | GSE215895 | Normal |
| Dev&Aging | He, Peng et al. 2022. Cell, 10.1016/j.cell.2022.11.005 | Lung | 10X | read counts | E-MTAB-11278 | Normal |
| Dev&Aging | Miller, Alyssa J et al. 2020. Developmental cell, 10.1016/j.devcel.2020.01.033 | Lung | 10X | read counts | E-MTAB-8221 | Normal |
| Dev&Aging | Ren, Xianwen et al. 2021. Cell, 10.1016/j.cell.2021.01.053 | Lung | 10X | read counts | GSE158055 | Normal |
| Dev&Aging | Adams, Taylor S et al. 2020. Science advances, 10.1126/sciadv.aba1983 | Lung | 10X | read counts | GSE136831 | Normal |
| Dev&Aging | DePianto, Daryle J et al. 2021. JCI insight, 10.1172/jci.insight.143626 | Lung | 10X | read counts | GSE159354 | Normal |
| Dev&Aging | Madissoon, E et al. 2019. Genome biology, 10.1186/s13059-019-1906-x | Lung | 10X | read counts | PRJEB31843 | Normal |

| | | | | | | |
|---|---|---|---|---|---|---|
| Dev&Aging | Tian et al. 2022. Sig Transduct Target Ther, 10.1038/s41392-022-01150-4 | Lung | 10X | read counts | PRJCA006026 | Normal |
| Dev&Aging | Travaglini et al. 2020. Nature, 10.1038/s41586-020-2922-4 | Lung | 10X | read counts | EGAS00001004344 | Normal |
| Dev&Aging | Huang, Qiqing et al. 2022. Respiratory research, 10.1186/s12931-022-02293-2 | Lung | 10X | read counts | GSE171541 | Normal |

**Table S2. uHAF and references. (See also in the uHAF Items: https://lung.unifiedcellatlas.org/#/uHAFTree)**

| first level | second level | third level | fourth level | Markers | References |
|---|---|---|---|---|---|
| Endothelial cell | | | | PECAM1; CLDN5; VWF; CAV1; RAMP2 | 10.1016/j.ccell.2019.12.001<br>10.1186/s13059-019-1906-x<br>10.1183/13993003.02441-2018<br>10.1038/s41467-020-16164-1<br>10.1038/s41586-020-2157-4 |
| | Capillary endothelial cell | | | CA4; CD300LG; RAMP3 | 10.1161/CIRCULATIONAHA.120.052318<br>10.1158/0008-5472.CAN-17-2728 |
| | Vein endothelial cell | | | ACKR1; EMCN; BACE2 | 10.1016/j.ccell.2019.12.001<br>10.1038/ni.1675<br>10.1161/CIRCRESAHA.118.312913 |
| | Lymphatic endothelial cell | | | PROX1; PDPN; LYVE1; RELN | 10.1038/s41591-018-0096-5<br>10.1016/j.ccell.2019.12.001<br>10.1161/CIRCRESAHA.118.312913<br>10.1158/0008-5472.CAN-17-2728<br>10.1016/j.cell.2020.01.015 |
| | Artery endothelial cell | | | GJA5; BMX; STMN2; CYTL1 | 10.1161/CIRCRESAHA.118.312913<br>10.1161/CIRCULATIONAHA.120.052318<br>10.1016/j.ccell.2019.12.001<br>10.1158/0008-5472.CAN-17-2728 |
| | Aerocyte | | | EDNRB; TBX2; APLN; CA4 | 10.1038/s41586-020-2822-7<br>10.1161/CIRCULATIONAHA.120.052318 |

| | | | | | |
|---|---|---|---|---|---|
| | Tip cell | | | ETV5; ETV4 | 10.1242/dev.051656 |
| | Stalk cell | | | VWF; UBE2T; LRRC3B; FILIP1 | 10.1038/s41388-021-02054-3<br>10.1186/s12943-019-0987-1 |
| Acinar cell | | | | AKR1C3; PRSS2; PRSS3 | 10.15252/embr.201540946<br>10.1038/s41586-020-2157-4<br>10.1038/s41467-021-25725-x |
| Chondrocyte | | | | ACAN; SOX9; COL2A1; FMOD; COMP | 10.3233/BME-2010-0626<br>10.1016/j.joca.2021.06.010 |
| Neuroendocrine cell | | | | CHGA; CALCA; ASCL1; CRYM; HES6 | 10.1016/j.devcel.2020.01.033<br>10.1038/s41586-020-2157-4<br>10.1164/rccm.201911-2199OC |
| Submucosal gland cell | | | | LTF; SLPI; LYZ | 10.1016/j.devcel.2020.01.033<br>10.1073/pnas.2119759119 |
| | SMG mucous cell | | | MUC5B; MUC19; BPIFB2 | 10.1038/s41591-023-02327-2<br>10.1073/pnas.2119759119 |
| | SMG serous cell | | | BPIFA1; DMBT1 | 10.1038/s41591-023-02327-2<br>10.1073/pnas.2119759119 |
| | SMG duct cell | | | MGST1; KRT19; MUC5AC; KLF5; LTF | 10.1038/s41591-023-02327-2<br>10.1073/pnas.2119759119 |
| Epithelial cell | | | | EPCAM; CAPS; SNTN | 10.1038/s41591-018-0096-5<br>10.1016/j.ebiom.2017.02.015<br>10.1038/nature24489 |

| | | | | | |
|---|---|---|---|---|---|
| | Tuft cell | | | ASCL2; DCLK1; LRMP; CD24A | 10.1038/s41586-018-0394-6<br>10.1038/nature24489<br>10.1016/j.jcmgh.2022.02.007 |
| | Secretory cell | | | SCGB3A2; SCGB3A1; BPIFB1; WFDC2; SERPINB3 | 10.1038/s41467-019-09639-3<br>10.1038/s41467-020-16239-z<br>10.1038/s41586-020-2157-4<br>10.1186/s13059-019-1906-x<br>10.1126/sciadv.aba1972 |
| | Goblet cell | | | MUC5AC; MUC5B; BPIFB1; KLF4; HES6 | 10.1038/s41525-020-00151-y<br>10.1038/s41577-020-00477-9<br>10.1038/nature24489<br>10.1016/j.celrep.2019.04.052 |
| | Alveolar cell | | | AQP4; CLDN18 | 10.1038/s41591-018-0096-5<br>10.1172/JCI8258<br>10.1172/JCI99799 |
| | | Type I alveolar cell | | AGER; CAV1 | 10.1038/s41467-019-08617-z<br>10.1038/s41591-019-0468-5<br>10.14573/altex.1511131<br>10.1038/s41467-022-28062-9 |
| | | Type II alveolar cell | | SFTPA1; SFTPC; SFTPD; SERPINA1 | 10.1016/j.ajhg.2016.07.007<br>10.1038/s41586-020-2157-4<br>10.1038/s41388-020-01528-0<br>10.1038/s41591-019-0468-5 |

| | | | | | |
|---|---|---|---|---|---|
| | Ciliated cell | | | FOXJ1; PIFO; TPPP3 | 10.1126/sciadv.aaw3413<br>10.1038/s41591-019-0468-5<br>10.1183/13993003.02441-2018<br>10.1164/rccm.201712-2410OC |
| | Basal cell | | | KRT5; KRT15; KRT16; KRT6A; KRT7; S100A2; TP63 | 10.1164/rccm.201712-2410OC<br>10.1038/s41586-018-0394-6<br>10.1038/s41467-021-22801-0<br>10.1038/s41586-018-0394-6<br>10.1242/dev.143784 |
| | | Suprabasal cell | | KRT19; KRT5; MKI67; KRT13 | 10.1038/s41591-023-02327-2<br>10.1164/rccm.201911-2199OC |
| | | Basal resting cell | | KRT15; KRT17; TP63 | 10.1038/s41591-023-02327-2 |
| | Ionocyte cell | | | ASCL3; FOXI1; RARRES2; TFCP2L1 | 10.1016/j.omtn.2021.06.010<br>10.1101/cshperspect.a035733<br>10.1038/s41586-018-0394-6<br>10.1164/rccm.201911-2199OC |
| | Squamous cell | | | SCEL; SPRR1A; SPRR1B; KRT13 | 10.1164/rccm.201911-2199OC<br>10.1038/s41422-020-00455-9 |
| | FOXN4+ cell | | | FOXN4; UBE2T | 10.1038/s41586-018-0394-6 |
| | Epithelial progenitor cell | | | CEACAM6; TM4SF1 | 10.1369/0022155415603768<br>10.1038/s41586-020-2157-4 |

| | | | | | |
|---|---|---|---|---|---|
| | Deuterosomal cell | | | MCIDAS; HES6; PLK4; DEUP1; YPEL1 | 10.1038/s41577-020-00477-9<br>10.1111/all.15705<br>10.1242/dev.177428 |
| Myeloid cell | | | | LYZ; MARCO; CD14; CD68 | 10.1038/s41591-018-0096-5<br>10.1016/j.ccell.2019.12.001<br>10.1038/s41592-019-0529-1 |
| | Mast cell | | | GATA2; MS4A2; TPSAB1; TPSB2; HDC | 10.1016/j.cell.2020.03.048<br>10.1183/13993003.02441-2018<br>10.3389/fimmu.2018.02193 |
| | Neutrophilic granulocyte | | | CSF3R; CXCR2; FCGR3A; FCGR3B | 10.1073/pnas.1908576116<br>10.1007/s13238-020-00752-4<br>10.1038/s41586-020-2157-4<br>10.1016/j.immuni.2019.03.009 |
| | Eosinophilic granulocyte | | | IL5RA; GATA2; CD125; CCL13; CCL11; RNASE3 | 10.1016/S0091-6749(98)70177-0<br>10.1136/thoraxjnl-2020-215167<br>10.1038/ncomms15081<br>10.1016/j.immuni.2013.10.003 |
| | Basophilic granulocyte | | | CCR3; IL3RA; CLC; MS4A3; CD203C; CD63 | 10.1111/j.1398-9995.2010.02431.x<br>10.1054/homp.2002.0082<br>10.1016/j.jaci.2009.10.074<br>10.1016/j.jim.2006.12.002 |
| | Promyelocyte | | | ITGA4; CD33; FUT4; CTSG; PRTN3 | 10.3389/fmicb.2019.00577<br>10.3324/haematol.2019.219048 |

| | | | | | |
|---|---|---|---|---|---|
| | Dendritic cell | | | MS4A6A; CLEC10A; GD83; LAMP3 | 10.1186/s13059-019-1906-x<br>10.1038/s41586-020-2157-4<br>10.1038/s41467-020-16164-1<br>10.1038/modpathol.3880254 |
| | | Conventional dendritic cell | | CD1C; THBD; CLEC9A; BTLA; CADM1 | 10.1016/j.cell.2020.03.048<br>10.1186/s13059-019-1906-x<br>10.1084/jem.20200264<br>10.1038/s41467-020-16164-1 |
| | | Plasmacytoid dendritic cell | | CD123; CSF3R; IL3RA; LILRA4; TCF4; IRF7; NRP1; MZB1 | 10.1016/j.cell.2020.03.048<br>10.1111/imm.12888<br>10.1038/s41590-020-0743-0<br>10.3389/fimmu.2021.711329<br>10.1084/jem.20200264<br>10.1016/j.immuni.2019.03.009 |
| | | Mature dendritic cell | | LAMP3; CCL22; CCR7 | 10.1158/1078-0432.CCR-04-1448<br>10.1158/2159-8290.CD-19-0138 |
| | | Migratory dendritic cell | | CCR7; CCL17; CCL5; TMSB10; LAMP3 | 10.1126/science.abl5197<br>10.1084/jem.20222129 |

| | | | | | |
|---|---|---|---|---|---|
| | Monocyte | | | CD14; VCAN; CD16; FCN1; SRGN | 10.1016/j.cell.2020.03.048<br>10.1016/j.cell.2020.04.035<br>10.1186/s13059-019-1906-x<br>10.1016/j.immuni.2019.03.009<br>10.1038/s41467-020-16164-1<br>10.1038/s41586-020-2157-4 |
| | | Classical monocyte | | CD14; CD16-; S100A8; S100A12 | 10.1038/s41388-021-02054-3<br>10.1126/science.aah4573<br>10.1016/j.cell.2020.08.002<br>10.1038/s41392-021-00703-3 |
| | | Non-classical monocyte | | CD16; CD14-; FCGR3A; CSF1R; LILRB1; LILRB2 | 10.1016/j.immuni.2019.03.009<br>10.1126/science.aah4573<br>10.1016/j.cell.2020.08.002<br>10.1016/j.jhep.2023.02.040 |
| | | CD14 monocyte | | CD14; FCN1; S100A8; S100A9 | 10.1016/j.cell.2021.01.010<br>10.1038/s41467-024-50478-8 |
| | | CD16 monocyte | | CD16; FCGR3A; CSF1R; LILRB2; LST1 | 10.1016/j.cell.2021.01.010<br>10.1038/s41467-024-50478-8 |
| | | Intermediate monocyte | | CD14; CD16 | 10.1084/jem.20170355<br>10.1126/science.aah4573<br>10.1182/blood-2009-07-235028 |

| | | | | | |
|---|---|---|---|---|---|
| | | Alveolar macrophage | | GPNMB; FABP4; TREM2; MARCO | 10.1016/j.immuni.2020.12.003<br>10.1016/j.cell.2020.08.002<br>10.1038/s41467-020-16164-1<br>10.1152/ajplung.00223.2022 |
| | | Promonocyte | | MPO; VCAN; CD11b; S100A8 | 10.1038/s41586-021-03929-x<br>10.1038/s41467-021-25725-x |
| | Macrophage | | | CD68; CD163; MRC1; MARCO; CTSB | 10.1182/blood-2012-06-436212<br>10.1038/s41590-019-0398-x<br>10.3390/cancers11050689<br>10.1038/s41467-020-16164-1 |
| | | Erythrophagocytic macrophage | | CD5L; SCL40A1; SPIC | 10.1186/s13059-019-1906-x |
| | | Monocyte derived macrophage | | TNIP3; TGFBI; CLEC12A; CD68; MARCO- | 10.1126/science.abl5197<br>10.1038/s41593-020-00789-y<br>10.1038/s41467-018-06318-7<br>10.1038/s41574-022-00675-6 |
| | | Interstitial macrophage | | MERTK; ADGRE1; C5AR1; C1QC; C1QA | 10.1016/j.cellimm.2018.02.001<br>10.1002/JLB.3RU0720-418R<br>10.1152/ajplung.00223.2022 |
| | | M1 macrophage | | CD86; CD80; CD282; IFIT1; ISG15; IFITM1 | 10.3389/fimmu.2015.00263<br>10.1016/j.redox.2022.102463<br>10.1038/s41587-020-0462-y<br>10.1016/j.jare.2022.04.006 |

| | | | | | |
|---|---|---|---|---|---|
| | | M2 macrophage | | CD163; MRC1(CD206); ARG1; CD200R | 10.1016/j.cellimm.2013.01.010<br>10.3389/fimmu.2015.00263<br>10.1016/j.redox.2022.102463<br>10.1038/s41587-020-0462-y<br>10.1016/j.jare.2022.04.006 |
| | | CD169 macrophage | | CD169 | 10.1038/nm.3057<br>10.3389/fimmu.2015.00263 |
| | Megakaryocyte | | | PPBP; PF4; ITGA2B | 10.1126/sciadv.abm5900<br>10.1016/j.cell.2011.01.004<br>10.1182/blood.2020006229 |
| | Megakaryocyte–erythroid progenitor cell | | | GATA2; CD45RA; CD38; KLF1 | 10.1038/s41586-019-1652-y<br>10.1016/j.stem.2012.01.006 |
| | Granulocyte-monocyte progenitor cell | | | CD38; CALR | 10.1016/j.cell.2011.01.004<br>10.1038/s41590-017-0001-2 |
| | Hematopoietic progenitor cell | | | CD34; SPINK2; PRSS57 | 10.1183/13993003.00827-2019<br>10.1182/blood.2020006229<br>10.1172/JCI147343 |
| | Common myeloid progenitor cell | | | CD34; CD38; MPO | 10.1183/13993003.00827-2019<br>10.1016/j.stem.2012.01.006<br>10.1172/JCI147343<br>10.1038/s41422-020-0378-6 |

| | | | | | |
|---|---|---|---|---|---|
| | Erythrocyte | | | HBB; HBA1; ALAS2; PTPRC; GYPA; TUBB1 | 10.1038/s41586-020-2157-4<br>10.1038/s41598-018-30047-y<br>10.1186/s13059-020-02048-6<br>10.1038/s41556-022-01064-x |
| Smooth muscle cell | | | | CNN1; MYH11; ACTA2; TAGLN; CSPG4; TPM2 | 10.1126/sciadv.aba1983<br>10.1038/s41586-020-2157-4<br>10.1002/ijc.33995<br>10.1016/j.devcel.2022.09.015 |
| | Vascular smooth muscle cell | | | ACTC1; ACTA2; NOTCH3; PDGFRB | 10.1038/s41586-020-2922-4<br>10.1038/s41592-019-0529-1<br>10.1016/j.devcel.2022.09.015 |
| | Bronchial smooth muscle cell | | | DACH2; CSPG4-; CHRM2 | 10.1038/s41586-020-2922-4<br>10.1016/j.devcel.2022.09.015<br>10.1038/s41556-022-01064-x |
| | Fibromyocyte | | | MYH11; SCX; LGR6; TCF21; CNN1; TAGLN | 10.1038/s41586-020-2922-4<br>10.1038/s41556-022-01064-x<br>10.1038/s41569-019-0255-5 |
| Mesothelial cell | | | | MSLN; KRT18; KRT19; WT1; UPK3B | 10.1371/journal.pone.0025391<br>10.1016/j.celrep.2018.03.010<br>10.1038/s41556-022-01064-x<br>10.1152/ajplung.00424.2012<br>10.1038/ncb2610 |

| | | | | | |
|---|---|---|---|---|---|
| Mesenchymal cell | | | | COL1A22; ACTA22; PDGFRB; ZEB1; CDH2 | 10.1016/j.ygyno.2017.01.015<br>10.1038/s41556-022-01064-x<br>10.1038/s41586-020-2922-4 |
| Pericyte | | | | CSPG4; RGS5; RERGL; PDGFRB | 10.1126/science.aay5356<br>10.1038/s41556-022-01064-x |
| Schwann cell | | | | MPZ; S100A1; CDH19; PLP1 | 10.1002/jemt.10304<br>10.1038/s41556-022-01064-x<br>10.1016/j.stemcr.2017.04.011 |
| Neuron | | | | ASCL1; PRPH; NRG1; PHOX2B; NEUROD1 | 10.1002/jbm.b.33055<br>10.1089/neu.2010.1579<br>10.1038/s41556-022-01064-x |
| Fibroblast | | | | DCN; LUM; COL1A2; COL1A1; SPARC | 10.1038/s41467-017-02289-3<br>10.1038/s41421-020-00200-x<br>10.1016/j.immuni.2019.03.009<br>10.1038/s41586-020-2157-4 |
| | Myofibroblast | | | ACTA2; COL3A1; ASPN; WIF1; COL6A3 | 10.1152/ajplung.90264.2008<br>10.1038/s41586-020-2922-4<br>10.1038/s41556-022-01064-x |
| | Alveolar fibroblast | | | CFD; LUM; GPC3; FGFR4 | 10.1038/s41586-020-2922-4<br>10.1038/s41556-022-01064-x |
| | Adventitial fibroblast | | | SFRP2; SERPINF1; TAGLN | 10.1038/s41586-020-2922-4<br>10.1038/s41586-020-2922-4<br>10.1038/s41556-022-01064-x |

| | | | | | |
|---|---|---|---|---|---|
| | Airway fibroblast | | | ASPN; S100A4; FGF7; TNC | 10.1016/j.cell.2022.11.005<br>10.1038/s41556-022-01064-x |
| | Subpleural fibroblast | | | APOC1; MMP23B | 10.1038/s41591-023-02327-2<br>10.3389/fphar.2015.00113 |
| | Pathological fibroblast | | | CXCL10; CCL19; COL1A1; CTHRC1 | 10.1038/s41586-021-03569-1<br>10.1016/j.medj.2022.05.002 |
| | Lipofibroblast | | | APOE; PLIN2; LPI; MLC1; PTPN12 | 10.1038/s41467-020-15647-5<br>10.1038/s41586-020-2922-4<br>10.1038/s41556-022-01064-x |
| Lymphocyte | | | | PTPRC; JCHAIN; CCL5; CD3D; CD45 | 10.3390/jcm4081600<br>10.1186/s13059-019-1830-0<br>10.3390/jcm4081600 |
| | B cell | | | CD19; CD79A; CD79B; CD74; MS4A1; IL7R; RAG1; CD80 | 10.1016/j.cellimm.2005.08.026<br>10.1371/journal.pone.0193539<br>10.1016/j.immuni.2019.03.009<br>10.1016/j.it.2022.01.003<br>10.1186/1471-2164-7-115 |
| | | Plasma cell | | JCHAIN; MZB1; IGHG1; IGKC; PRDM1; XBP1 | 10.1126/science.aat1699<br>10.1126/sciadv.aba1972<br>10.1038/s41467-020-16164-1<br>10.1002/ctm2.1346 |

| | | | | | |
|---|---|---|---|---|---|
| | | Plasmablast cell | | MKI67; CD38 | 10.1111/apm.12738<br>10.1038/s41591-020-0944-y<br>10.1016/j.immuni.2020.11.017 |
| | | Naive B cell | | MS4A1; CD22; CD37; CD19; TCL1A | 10.1155/2022/2079389<br>10.4049/jimmunol.2100132<br>10.1038/s41392-021-00753-7<br>10.1002/ctm2.1346 |
| | | Memory B cell | | CD27; CD19; CD80; CD180 | 10.1007/s40120-018-0101-4<br>10.1007/978-1-4939-1161-5_4<br>10.1016/j.immuni.2020.07.009<br>10.4049/jimmunol.182.2.890<br>10.1038/ni.2914 |
| | | Immature B cell | | CD19; CD24; IGHM; RAG2; IGLL5 | 10.1038/nri1633<br>10.1016/j.it.2022.01.003<br>10.1038/s41556-019-0439-6 |
| | | Mature B cell | | CD19; CD20 (MS4A2); PAX5; CD79A; CXCR4; RSP27 | 10.1038/nri3487<br>10.1038/nri1633<br>10.1016/j.it.2022.01.003<br>10.1126/science.abf9277 |
| | | Follicular B cell | | CD69; CD22; CD21-; CD74; CD81 | 10.1111/imm.12724<br>10.1111/j.1365-2567.2006.02386.x<br>10.1002/ctm2.1346 |

| | | | | | |
|---|---|---|---|---|---|
| | | Transitional B cell | | CD38; CD24 | 10.1111/apm.12738<br>10.1038/s41586-019-1922-8 |
| | | Marginal zone B | | CD27; CD22; CD1; CD180; CD23- | 10.1038/nri799<br>10.1038/s41421-020-0157-z |
| | | Germinal center B cell | | MKI67; HGAL | 10.1097/pai.0b013e31826399aa<br>10.1002/ctm2.1346 |
| | | Pre-B cell | | CD24; RAG1; IL2RA; IGH; CD19; IL7R | 10.1038/nri1633<br>10.1038/s41556-019-0439-6<br>10.1126/science.abf9277<br>10.1016/j.it.2022.01.003 |
| | | Pro-B cell | | CD19; CD10; RAG2; VPREB; CD179B; MKI67; IL7RA | 10.1038/nri1633<br>10.1126/science.abf9277<br>10.1038/s41590-020-0772-8<br>10.1016/j.stem.2020.11.015 |
| | T cell | | | CD3D; CD3E; CD3G | 10.1038/s41592-019-0529-1<br>10.1038/nbt.3973<br>10.1016/j.immuni.2019.03.009 |
| | | Exhausted T cell | | CD160; BTLA; CD58; LAG3; TIGIT; TIM3 | 10.1038/s41467-019-12464-3<br>10.1038/s41591-018-0045-3<br>10.1038/s41467-020-16164-1<br>10.1126/science.abe6474<br>10.1038/ni.2035 |

| | | | | | |
|---|---|---|---|---|---|
| | | CD4 T cell | | CD4; IL7R; CD3D; CCR7 | 10.1038/s41586-018-0792-9<br>10.1038/s41467-020-16164-1<br>10.1038/s41467-021-25101-9<br>10.1016/j.immuni.2020.11.017 |
| | | | Naive CD4 T cell | CD45RA; CD45RO-; CCR7; CD27; CD28; CXCR3; CCR4; CCR6; TMSB4X | 10.1038/s41467-019-12464-3<br>10.1038/s41586-018-0792-9<br>10.1038/s41467-019-12464-3<br>10.1038/s41591-018-0045-3 |
| | | | Memory CD4 T cell | CD45RA-; CD45RO; IL2RB; CD95; CD122; TRAC; S100A11; PTPRC | 10.3390/cancers8030036<br>10.1016/0167-5699(88)91212-1<br>10.1016/bs.ircmb.2018.05.007 |
| | | | Effector CD4 memory T cell | CD45RA-; CD45RO; CD95; CD122; IL7R; PTPRC | 10.3390/cancers8030036<br>10.1016/bs.ircmb.2018.05.007<br>10.4049/jimmunol.178.7.4112 |
| | | NKT cell | | CD3E; KLRF1; NKG7; GNLY; GATA3; PITPNC1; CCL5 | 10.1126/science.aat1699<br>10.1136/gutjnl-2021-325915 |
| | | Treg cell | | CD25; FOXP3; IL2RA; CTLA4; TGFB1 | 10.1038/s41590-018-0051-0<br>10.1038/s41586-018-0792-9<br>10.1182/blood-2013-02-482539<br>10.1038/nri2785 |
| | | T helper cell | | CD3D; CD3E; CCR6; IL7R; CXCR3 | 10.1186/ar3257<br>10.1126/science.aat1699 |

| | | | | | |
|---|---|---|---|---|---|
| | | | Th1 cell | CD182; CXCR3; CCR4-; CCR6-; ITGA4 | 10.1089/dna.2015.3105<br>10.1002/cptx.26<br>10.1556/APhysiol.99.2012.3.7<br>10.1146/annurev-immunol-032414-112056<br>10.1038/s41467-021-21043-4 |
| | | | Th2 cell | CCR4; CXCR4; GATA3; CCR6-; CXCR3-; CD184 | 10.1002/cptx.26<br>10.1016/bs.ircmb.2018.05.007<br>10.4049/jimmunol.1002828<br>10.1084/jem.20040774<br>10.1146/annurev-immunol-032414-112056 |
| | | | Th9 cell | TGFB1; IL9; IL4 | 10.3390/cancers8030036 |
| | | | Th17 cell | CCR6; CXCR3-; CCR4; IL17A; IL17F; CD146; CD161 | 10.1111/j.1346-8138.2012.01544.x<br>10.1016/j.cell.2020.10.001<br>10.1111/j.1346-8138.2012.01544.x<br>10.1002/eji.200940257 |
| | | | Follicular helper T cell | CXCR5; IL6; PDCD1; ICOS; TIGIT; TNFRSF4; CXCL13 | 10.3390/cancers8030036<br>10.1016/bs.ircmb.2018.05.007<br>10.4049/jimmunol.1002828<br>10.1146/annurev-immunol-032414-112056 |
| | | CD8 T cell | | CD8A; CD8B | 10.1038/s41586-018-0792-9<br>10.1016/j.cell.2018.09.006 |

| | | | | | |
|---|---|---|---|---|---|
| | | | Naive CD8 T cell | CCR7; CXCR4; LEF1; CD45RA; CD45RO-; CD127; CD28 | 10.3390/cancers8030036<br>10.1038/nm.2446<br>10.1016/bs.ircmb.2018.05.007<br>10.1136/gutjnl-2021-325915 |
| | | | Memory CD8 T cell | CD45RA; CD45RO-; CCR7; CD27; CXCR3; CD28; CD122; CD58 | 10.1128/jvi.02528-08<br>10.1016/bs.ircmb.2018.05.007<br>10.1182/blood-2002-11-3577 |
| | | | Effector CD8 memory T cell | CCR5; CX3CR1; PDCD1; IFNG | 10.1016/j.cell.2017.05.035<br>10.1016/bs.ircmb.2018.05.007<br>10.4049/jimmunol.178.7.4112<br>10.1136/gutjnl-2021-325915 |
| | | Cytotoxic T cell | | PRF1; CD3; CD8 | 10.1038/s41416-020-01048-4<br>10.3389/fimmu.2022.816005<br>10.1016/j.tranon.2021.101042 |
| | | Effector T cell | | CD39; PTPRC; IL2RB; KLRG1; GNLY | 10.2500/ajra.2013.27.3958<br>10.1172/jci.insight.154646<br>10.1136/gutjnl-2021-325915 |
| | | MAIT cell | | IL32R; SLC4A10 | 10.1016/j.cell.2017.05.035<br>10.1038/s41591-020-1003-4<br>10.1136/gutjnl-2021-325915 |
| | | Cycling T cell | | PCNA; KI67 | 10.1186/s12865-022-00516-1<br>10.1016/j.immuni.2020.11.016<br>10.1038/s41421-020-0157-z |

| | | | | | |
|---|---|---|---|---|---|
| | | Proliferating T cell | | MKI67; TK1; CENPW | 10.1038/s41591-023-02327-2<br>10.1136/gutjnl-2021-325915<br>10.1111/imm.13388 |
| | | Gamma delta T cell | | TRDV2; SCART1 | 10.1016/j.cell.2021.01.053<br>10.1016/j.molimm.2010.03.002 |
| | NK cell | | | KLRD1; NKG7; FCGR3A; NCR1 | 10.1038/ncomms14049<br>10.1038/s41467-019-11947-7<br>10.1016/j.immuni.2019.03.009 |
| | Innate lymphoid cell | | | CD161; CRTH2; CD127; GTA3 | 10.1038/s41422-020-00445-x<br>10.1038/ni.2104<br>10.1016/j.jaci.2021.07.025<br>10.1002/hep.32444 |
| | Lymphoid progenitor cell | | | IL7R; CD7; CD34 | 10.1038/sj.leu.2404488<br>10.1016/j.immuni.2019.09.008 |
| Cancer cell | | | | | |

**Table S3. Original cell label coordination by uHAF.**

| First level | Second level | Third level | Fourth level | Carraro, G_2021_Nature medicine | Li, X_2022_Life science alliance | Sikkema, L_2023_Nature medicine | Wang, A_2020_eLife | Melms, J_2021_Nature | Miller AJ_2020_Developmental Cell | Adams TS_2020_Sci Adv | Misiukiewicz-Stępień P_2022_Int J Mol Sci | Madissoon, E_2020_Genome Biol | Chan JM_2021_Cancer Cell | Salcher S_2022_Cancer Cell | Travaglini KJ_2020_Nature | Vieira Braga_2019_Nature medicine | Okuda, K_2021_American journal of respiratory and critical care medicine | Wang, A_2020_eLife | Sarah Kim-Hellmuth_2020_Science | Ren X_2021_Cell |
|---|---|---|---|---|---|---|---|---|---|---|---|---|---|---|---|---|---|---|---|---|
| Endothelial cell | | | | | | | | Endothelial cells (other)<br>Endothelial cells (general)<br>Lymphatic endothelial cell | Endothelial | Multiplet_VascEndoB_VascEndoD<br>Multiplet_VE_Stromal | endothelial cells | Blood_vessel | Endothelial | | Bronchial Vessel 1 | Endothelial<br>Endothelium | Endothelial | bronchial vessel | | |
| | Capillary endothelial cell | | | | | EC general capillary<br>EC aerocyte capillary | Cap1<br>Cap2 | Capillary endothelial cells | | VE_Capillary | | | | capillary endothelial cell | Capillary Intermediate Capillary | | | Cap | Endothelial cell (vascular) III | |
| | Vein endothelial cell | | | | | EC venous systemic<br>EC venous pulmonary | veins<br>bronchial vessel | Pulmonary venous endothelial cells<br>Systemic venous endothelial cells | | VE_Venous | | | | vein endothelial cell | | | | veins | Endothelial cell (vascular) II | |
| | Lymphatic endothelial cell | | | | | Lymphatic EC mature<br>Lymphatic EC | lymphatics | | | Lymphatic-Endothelial | | Lymph_vessel | | endothelial cell of lymphatic vessel | Lymphatic | Lymphatic | | lymphatics | Endothelial cell (lymphatic) | |

| | | | | | | | | | | | | | |
|---|---|---|---|---|---|---|---|---|---|---|---|---|---|
| | | | differentiating | | | | | | | | | | |
| | Artery endothelial cell | | EC arterial | arteries | Arterial endothelial cells | | VE_Arterial | AE1 | pulmonary artery endothelial cell | Artery | | Artery | Endothelial cell (vascular) I |
| | Aerocyte | | | | | | | | | Capillary Aerocyte | | | |
| | Tip cell | | | | | | | | | | | | |
| | Stalk cell | | | | | Bud tip adjacent | | | | | | | |
| Acinar cell | | | | | | | | | | | | | |
| Chondrocyte | | | | chondrocytes | | Cartilage | | | | | | chondrocytes | |
| Neuroendocrine cell | | | Neuroendocrine | | | Neuroendocrine | | Neuroendocrine | | | | | |
| Submucosal gland cell | | | | | | Submucosal gland | | | | | | | |
| | SMG mucous cell | | SMG mucous | | | | | | | | | | |
| | SMG serous cell | | SMG serous (bronchial)<br>SMG serous cell (nasal) | | | | | | | | | | |
| | SMG duct cell | | SMG duct | | | | | | | | | | |
| Epithelial cell | | Epi | | | ECM-high epithelial<br>Cycling epithelial | Epithelial | Mystery_Disease_Epithelial | AE1<br>AEP | | | Transformed epithelium | Epi | |

| | | | | | | | | | | | | | | | | | |
|---|---|---|---|---|---|---|---|---|---|---|---|---|---|---|---|---|---|
| | | | | | Airway mucous | | | | | | | | | | | | |
| Tuft cell | | | Tuft | | Tuft-like | | | | | Tuft | | | | | | | |
| Secretory cell | | Secretory | pre-TB secretory<br>Club (non-nasal)<br>AT0<br>Club (nasal) | club cells | Airway club | Secretory progenitor<br>Club-like secretory | Club<br>Club_CellCycle<br>Multiplet_Secretory_Ciliated | club cells | Secretory_club | Club | club cell | Club | Secretory | Secretory | club cells | Epithelial cell (club) | Epi-Secretory |
| Goblet cell | | | Goblet (bronchial)<br>Goblet (nasal)<br>Goblet (subsegmental) | goblet cells | Airway goblet | Goblet-like secretory | Goblet<br>Goblet_MT-tRNAs | | | | | Goblet | | | gobleT cell | | |
| Alveolar cell | | | | | | | | | | | | | | Alveolar | | | |
| | Type I alveolar cell | | AT1 | alveolar type 1 cells | AT1 | | ATI | alveolar type 1 cells | Alveolar_Type1 | | type I pneumocyte | Alveolar Epithelial Type 1 | Type 1 | | alveolar type 1 cells | Epithelial cell (alveolar type I) | |
| | Type II alveolar cell | | AT2<br>proliferating AT2 | alveolar type 2 cells | AT2 | | ATII_High-Surfactants<br>ATII_Low-Surfactants | alveolar type 2 cells | Alveolar_Type2 | | type II pneumocyte | Alveolar Epithelial Type 2<br>Signaling Alveolar Epithelial Type 2 | Type 2 | | alveolar type 2 cells | Epithelial cell (alveolar type II) | Epi-AT2 |
| Ciliated cell | | Ciliated | Multiciliated (non-nasal)<br>Multicili | ciliated cells | Airway ciliated | Multiciliated precursor<br>Multiciliated cell | Ciliated<br>Multiplet_Secretory_Ciliated<br>Ciliated_CellCycle | ciliated cells | Ciliated | Ciliated | multi-ciliated epithelial cell | Ciliated | Ciliated | Ciliated | ciliated cells | Epithelial cell (ciliated) | Epi-Ciliated |

| | | | | | | | | | | | | | | | | | | | |
|---|---|---|---|---|---|---|---|---|---|---|---|---|---|---|---|---|---|---|---|
| | | | | | ated (nasal) | | | Intermediate ciliated | | | | | | | | | | | |
| | Basal cell | | Basal | | Hillock-like | basal cells | Airway basal | Basal cell | Basal | | Basal | Basal | | Basal | Basal | Basal | basal cells | Epithelial cell (basal) | |
| | | Suprabasal cell | | | Suprabasal | | | | | | | | | | | Suprabasal | | | |
| | | Basal resting cell | | | Basal resting | | | | | | | | | | | | | | |
| | Ionocyte cell | | Ionocyte | | Ionocyte | | | | | | | Ionocyte | | | Ionocytes | | | | |
| | Squamous cell | | | | | | | | | | | | | | | | | | Epi-Squamous |
| | FOXN4+ cell | | FOXN4+ | | | | | | Ciliated_FOXN4+ | | | | | | | | | | |
| | Epithelial progenitor cell | | | | | | | Epithelial progenitor cell | | | | | | | | | | | |
| | Deuterosomal cell | | | | Deuterosomal | | | | | | | | | | | | | | |
| Myeloid cell | | | | Cyc.Mye | | | | | | | | | myeloid cell | | | | Cyc.Mye | | |
| | Mast cell | | | | Mast cells | mast cells | Mast cells | | Mast Multiplet_Mast_Mac | | Mast_cells | Mast | mast cell | | | | | Immune (mast cell) | |
| | Neutrophilic granulocyte | | NE | | | | | | | neutrophils | | Neutrophil | neutrophil | Neutrophil | | | | | Neu_c1-IL1B<br>Neu_c2-CXCR4(low)<br>Neu_c3-CST7<br>Neu_c4-RSAD2<br>Neu_c5-GSTP1(high)OASL(low) |

| | | | | | | | | | | | | | | | | | | | |
|---|---|---|---|---|---|---|---|---|---|---|---|---|---|---|---|---|---|---|---|
| | | | | | | | | | | | | | | | | | | | Neu_c6-FGF23 |
| | Eosinophilic granulocyte | | | | | | | | | | | | | | | | | | |
| | Basophilic granulocyte | | | | | | | | | | | | | Basophil | | | | | |
| | Promyelocyte | | | | | | | | | | | | | | | | | | |
| | Dendritic cell | | | | | dendritic cells | Dendritic cells | | Dendritic<br>Dendritic_CellCycle<br>Dendritic_Langerhans<br>Dendritic_MT-tRNAs | dendritic cells | DC_activated | DC | dendritic cell | IGSF21+ Dendritic | Dendritic cells | | dendritic cells | | |
| | | Conventional dendritic cell | | DC1/DC2 | DC1/DC2 | | | | | | DC_2<br>DC_1 | | cDC | | | | DC1/DC2 | | DC_c1-CLEC9A<br>DC_c2-CD1C |
| | | Plasmacytoid dendritic cell | | pDCs | Plasmacytoid DCs | | | | Dendritic_Plasmacytoid | | DC_plasmacytoid | | plasmacytoid dendritic cell | Plasmacytoid Dendritic | | | pDCs | | DC_c4-LILRA4 |
| | | Mature dendritic cell | | | | | | | | | | | | | | | | | DC_c3-LAMP3 |
| | | Migratory dendritic cell | | Mig.DCs | Migratory DCs | | | | | | | | | | | | Mig.DCs | | |
| | Monocyte | | | Mono | | monocytes | Monocytes | | Monocyte<br>Monocyte_Outlier | monocytes | Monocyte | | | Intermediate Monocyte | | | Mono | | Mono_c1-CD14-CCL3<br>Mono_c2-CD14-HLA- |

| | | | | | | | | | | | | | | | | | |
|---|---|---|---|---|---|---|---|---|---|---|---|---|---|---|---|---|---|
| | | | | | | | | | | | | | | | | | DPB1<br>Mono_c4-<br>CD14-<br>CD16<br>Mono_c5-<br>CD16 |
| | Classical<br>monocyte | | Classica<br>l<br>monocyt<br>es | | | | | | | classical<br>monocyte | | | | | | | |
| | Non-<br>classical<br>monocyte | | Non-<br>classical<br>monocyt<br>es | | | Monocyte_Non-<br>Classical | | | | non-<br>classical<br>monocyte | Nonclassi<br>cal<br>Monocyte | | | | | | |
| | Intermedi<br>ate<br>monocyte | | | | | | | | | | | | | | | | |
| | Alveolar<br>macropha<br>ge | AMs | Alveolar<br>macroph<br>ages<br>Alveolar<br>Mph<br>CCL3+<br>Alveolar<br>Mph<br>prolifera<br>ting<br>Alveolar<br>Mph<br>MT-<br>positive | Alveol<br>ar<br>macrop<br>hages | Alveolar<br>macrophages | Macrophage_Alv<br>eolar<br>Macrophage_Alv<br>eolar_Outlier<br>Macrophage_Alv<br>eolar_CellCycle | | | | alveolar<br>macropha<br>ge | | | | AMs | Immune<br>(alveolar<br>macrophage<br>activated) | |
| | Promono<br>cyte | | | | | | | | | | | | | | | | Mono_c3-<br>CD14-<br>VCAN |
| Macrophage | | | | | | Macrophage_Out<br>lier<br>Macrophage_Out<br>lier_HSPs<br>Macrophage_MT<br>-tRNAs | macro<br>phages | Macrophage<br>_MARCOpo<br>s<br>Macrophage<br>_MARCOne<br>g | Macroph<br>age | macropha<br>ge | Macropha<br>ge | Luminal_<br>Macrophag<br>es | Macroph<br>age | | Immune<br>(macrophag<br>e) | Macro_c2-<br>CCL3L1<br>Macro_c3-<br>EREG<br>Macro_c4-<br>DNAJB1 |

| | | | | | | | | | | | |
|---|---|---|---|---|---|---|---|---|---|---|---|
| | | | | | | | | Multiplet_Macrophage_Low-Info | Macrophage_Dividing | | Macro_c5-WDR74 |
| | | Erythrophagocytic macrophage | | | | | Hematopoietic, Macrophage | | | | |
| | | Monocyte derived macrophage | FOLR2.IMs/SPP1.Ims | Monocyte-derived Mph | | Monocyte-derived macrophages Transitioning MDM | | | | | |
| | | Interstitial macrophage | | Interstitial Mph perivascular | Interstitial macrophages | | | | | SPP1.Ims FOLR2.Ims | Macro_c1-C1QC |
| | | M1 macrophage | | | | | | | | | |
| | | M2 macrophage | | | | | | | | | |
| | | CD169 macrophage | | | | | | | | | |
| | Megakaryocyte | | | | | | | | | | Mega |
| | Megakaryocyte–erythroid progenitor cell | | | | | | | | | | |
| | Granulocyte-monocyte progenitor cell | | | | | | | | | | |

| | | | | | | | | | | | | | |
|---|---|---|---|---|---|---|---|---|---|---|---|---|---|
| | Hematopoietic progenitor cell | | | | | | | | | | | | |
| | Common myeloid progenitor cell | | | | | | | | | | | | |
| | Erythrocyte | | | | | | | Erythrocyte | | | | enucleated erythrocytes | |
| Smooth muscle cell | | SM activated stress response<br>Smooth muscle<br>Smooth muscle FAM83D+ | | | | Smooth-Muscle Multiplet_SMCs_Pericytes | Muscle_cells | | smooth muscle cell | | Smooth muscle | | |
| | Vascular smooth muscle cell | | vascular smooth muscle | Vascular smooth muscle | | | | | | Vascular Smooth Muscle | | vascular smooth muscle | |
| | Bronchial smooth muscle cell | | airway smooth muscle | Airway smooth muscle | Airway Smooth Muscle | | | | | Airway Smooth Muscle | | airway smooth muscle | |
| | Fibromyocyte | | | | | | | | | | | | |
| Mesothelial cell | | Mesothelium | | | | Mesothelial | | | | | | | |
| Mesenchymal cell | | | matrix fibroblast 1 | | Mesenchyme RSPO2+<br>Mesenchyme SERPINF1-high | | | | | | | | |
| Pericyte | | Pericytes | pericytes | Pericytes | Pericyte | | | | pericyte | Pericyte | | pericytes | Pericyte/SMC |

| | | | | | | | | | | | | | | |
|---|---|---|---|---|---|---|---|---|---|---|---|---|---|---|
| Schwann cell | | | | | | | | | | | | | | |
| Neuron | | | | | Neuronal cells | | | | | | | | | |
| Fibroblast | | | | | Other FB/Mesothelial FB | | Fibroblast | Fibroblast | fibroblast of lung | | Fibroblasts | Fibroblast | matrix fibroblast | Fibroblast I |
| | Myofibroblast | | Myofibroblasts | myofibroblasts | | Myofibroblast | | | | Myofibroblast | | | myofibroblasts | |
| | Alveolar fibroblast | | Alveolar fibroblasts | | Alveolar FB | | | | | Alveolar Fibroblast | | | | |
| | Adventitial fibroblast | | Adventitial fibroblasts | | Adventitial FB | | | | | Adventitial Fibroblast | | | | |
| | Airway fibroblast | | Peribronchial fibroblasts | | | | | | bronchus fibroblast of lung | | | | | |
| | Subpleural fibroblast | | Subpleural fibroblasts | | | | | | | | | | | |
| | Pathological fibroblast | | | | Intermediate pathological FB/Pathological FB | | | | | | | | | |
| | Lipofibroblast | | | | | | | | | lipofibroblast | | | | |
| Lymphocyte | | Cyc.Lym/Lym | | | | Multiplet_Lymphocytes_Fibroblast<br>Multiplet_Lymphocytes_Ciliated<br>Multiplet_Lymphocyte_Secretory | | | | | | Lymphocyte | Lym | |

| | | | | | | | | | | | | | | | |
|---|---|---|---|---|---|---|---|---|---|---|---|---|---|---|---|
| B cell | | B cells | B cells | Activated B cells/B cells | Hematopoietic, B Cells | Multiplet_B_Mac<br>B cell | B cells | | B cell | B cell | B | B cells | B cells | Immune (B cell) | B_c01-TCL1A<br>B_c04-SOX5-TNFRSF1B<br>B_c05-MZB1-XBP1 |
| | Plasma cell | Plasma cells | | Plasma cells | | B_Plasma<br>B_Plasma_LowInfo | | Plasma_cells | Plasma cell | plasma cell | | | | | |
| | Plasmablast cell | | | | | | | | | | | | | | B_c06-MKI67 |
| | Naive B cell | | | | | | | B_cell_naive | | | | | | | B_c02-MS4A1-CD27 |
| | Memory B cell | | | | | | | | | | | | | | B_c03-CD27-AIM2 |
| | Immature B cell | | | | | | | | | | | | | | |
| | Mature B cell | | | | | | | B_cell_mature | | | | | | | |
| | Follicular B cell | | | | | | | | | | | | | | |
| | Transitional B cell | | | | | | | | | | | | | | |
| | Marginal zone B | | | | | | | | | | | | | | |
| | Germinal center B cell | | | | | | | | | | | | | | |
| | Pre-B cell | | | | | | | | | | | | | | |
| | Pro-B cell | | | | | | | | | | | | | | |
| T cell | | | T cells | | Hematopoietic, T Cells | T<br>Multiplet_T<br>Multiplet_T_Mac | T cells | T_cells_Dividing | T cell | | | T cells | T cells | Immune (T cell) | |

| | | | | | | | | | | |
|---|---|---|---|---|---|---|---|---|---|---|
| | Exhausted T cell | | | | | | | | | |
| | CD4 T cell | | CD4 T cells | CD4+ T cells | | T_CD4 | CD4-positive, alpha-beta T cell | | | T_CD4_c02-AQP3<br>T_CD4_c04-ANXA2<br>T_CD4_c05-FOS<br>T_CD4_c06-NR4A2<br>T_CD4_c07-AHNAK<br>T_CD4_c10-IFNG |
| | | Naive CD4 T cell | | | | | | CD4+ Naive T | | T_CD4_c01-LEF1 |
| | | Memory CD4 T cell | | | | | | | | T_CD4_c08-GZMK-FOS_h<br>T_CD4_c09-GZMK-FOS_l |
| | | Effector CD4 memory T cell | | | | | | CD4+ Memory/ Effector T | | |
| | NKT cell | | | | Multiplet_NK_T | | | Proliferating NK/T_P2<br>Natural Killer T_P2 | T and NK | T_CD4_c11-GNLY |
| | Treg cell | | | Tregs | T_Regulatory<br>Multiplet_Treg_Mac | T_regulatory | regulatory T cell | | | T_CD4_c12-FOXP3 |
| | T helper cell | | | | | | | | | |
| | | Th1 cell | | | | | | | | T_CD4_c03-ITGA4 |

| | | | | | | | | | |
|---|---|---|---|---|---|---|---|---|---|
| | | Th2 cell | | | | | | | |
| | | Th9 cell | | | | | | | |
| | | Th17 cell | | | | | | | |
| | | Follicular helper T cell | | | | | | | |
| | CD8 T cell | | CD8 T cells | CD8+ T cells | | T_CD8_CytT | CD8-positive, alpha-beta T cell | | T_CD8_c02-GPR183<br>T_CD8_c04-COTL1<br>T_CD8_c05-ZNF683<br>T_CD8_c06-TNF<br>T_CD8_c07-TYROBP<br>T_CD8_c13-HAVCR2 |
| | | Naive CD8 T cell | | | | | | CD8+ Naive T | T_CD8_c01-LEF1 |
| | | Memory CD8 T cell | | | | | | | T_CD8_c03-GZMK<br>T_CD8_c10-MKI67-GZMK |
| | | Effector CD8 memory T cell | | | | | | CD8+ Memory/ Effector T | |
| | Cytotoxic T cell | | | | T_Cytotoxic | | | | |
| | Effector T cell | | | | | | | | T_CD8_c08-IL2RB |
| | MAIT cell | | | | | | | | T_CD8_c09-SLC4A10 |

| | | | | | | | | | | | | | | | | |
|---|---|---|---|---|---|---|---|---|---|---|---|---|---|---|---|---|
| | | Cycling T cell | | | Cycling NK/T cells | | | | | | | | Cycling | | | T_CD4_c13-MKI67-CCL5_l<br>T_CD4_c14-MKI67-CCL5_h<br>T_CD8_c11-MKI67-FOS<br>T_CD8_c12-MKI67-TYROBP |
| | | Proliferating T cell | T cells proliferating | | | | | | | | | | | | | |
| | | Gamma delta T cell | | | | | | | | | | | | | | T_gdT_c14-TRDV2 |
| | NK cell | | NK cells | NK cells | NK cells | Hematopoietic, Natural Killer Cell | NK<br>Multiplet_NK_Mac | natural killer cells | NK/NK_Dividing | | natural killer cell | Natural Killer | | NK cells | Immune (NK cell) | NK_c01-FCGR3A<br>NK_c02-NCAM1<br>NK_c03-MKI67 |
| | Innate lymphoid cell | | | | | | Innate_lymphoid | | | | | | | | | |
| | Lymphoid progenitor cell | | | | | | | | | | | | | | | |
| Cancer cell | | | Cancer | | | | | | | NSCLC<br>SCLC<br>SCLC-N<br>SCLC-A<br>SCLC-P | Tumor cell | | | | | |

**Table S4. Statistical of comparing the re-annotation of the uniLUNG core with the original annotation at Level 1.**

| Cell type | Match_labelled | Cover_labelled | Unmatch_labelled |
|---|---|---|---|
| Cancer cell | 100 | 0 | 0 |
| Endothelial cell | 89.15022 | 0 | 10.84978 |
| Epithelial cell | 91.73197 | 0 | 8.268034 |
| Fibroblast | 55.47008 | 0 | 44.52992 |
| Lymphocyte | 96.75706 | 0 | 3.242941 |
| Myeloid cell | 96.77374 | 0 | 3.226262 |
| Neuroendocrine cell | 0 | 0 | 100 |
| Smooth muscle cell | 100 | 0 | 0 |
| Submucosal gland cell | 24.98615 | 0 | 75.01385 |

**Table S5. Statistical of comparing the re-annotation of the uniLUNG core with the original annotation at Level 2.**

| Cell type | Match_labelled | Cover_labelled | Unmatch_labelled |
|---|---|---|---|
| Adventitial fibroblast | 0.70145 | 9.165609 | 90.13294 |
| Airway fibroblast | 39.21555 | 55.56422 | 5.220228 |
| Alveolar cell | 68.73804 | 25.43393 | 5.82803 |
| Alveolar fibroblast | 2.642008 | 19.40555 | 77.95244 |
| Artery endothelial cell | 100 | 0 | 0 |
| B cell | 89.95412 | 1.960539 | 8.085341 |
| Basal cell | 79.14936 | 12.62197 | 8.228674 |
| Bronchial smooth muscle cell | 24.25232 | 75.74768 | 0 |
| Capillary endothelial cell | 41.27015 | 47.38841 | 11.34144 |
| Ciliated cell | 92.25651 | 3.410768 | 4.33272 |
| Dendritic cell | 100 | 0 | 0 |
| Deuterosomal cell | 0 | 11.05528 | 88.94472 |
| Erythrocyte | 53.43189 | 1.583949 | 44.98416 |
| Lymphatic endothelial cell | 81.53615 | 15.38654 | 3.077308 |

| | | | |
|---|---|---|---|
| Macrophage | 39.73804 | 56.65444 | 3.607524 |
| Mast cell | 97.92576 | 1.1711 | 0.903136 |
| Megakaryocyte | 98.40142 | 0.17762 | 1.420959 |
| Monocyte | 51.58661 | 46.38823 | 2.025154 |
| Myofibroblast | 0.150708 | 89.4666 | 10.38269 |
| NK cell | 70.74455 | 27.19921 | 2.056234 |
| Neutrophilic granulocyte | 89.89547 | 6.794425 | 3.310105 |
| SMG duct cell | 0 | 5.603448 | 94.39655 |
| SMG serous cell | 31.69277 | 0 | 68.30723 |
| Secretory cell | 62.45173 | 14.65989 | 22.88839 |
| Squamous cell | 84.43709 | 14.23841 | 1.324503 |
| T cell | 93.13533 | 4.481691 | 2.382979 |
| Vascular smooth muscle cell | 14.19122 | 85.80878 | 1.42E-14 |
| Vein endothelial cell | 42.72963 | 42.45375 | 14.81662 |

**Table S6. Statistical of comparing the re-annotation of the uniLUNG core with the original annotation at Level 3.**

| Cell type | Match_labelled | Cover_labelled | Unmatch_labelled |
|---|---|---|---|
| Basal resting cell | 14.09347 | 83.68215 | 2.224375 |
| CD4 T cell | 45.46842 | 53.51982 | 1.011766 |
| CD8 T cell | 45.21711 | 54.27943 | 0.503462 |
| Classical monocyte | 18.00366 | 79.7199 | 2.276439 |
| Conventional dendritic cell | 57.64376 | 42.35624 | 7.11E-15 |
| Cycling T cell | 100 | 0 | 0 |
| Cytotoxic T cell | 0.248756 | 94.36153 | 5.389718 |
| Follicular B cell | 0 | 100 | 0 |
| M1 macrophage | 0 | 98.71471 | 1.285288 |
| M2 macrophage | 0 | 95.54257 | 4.457432 |
| Mature B cell | 100 | 0 | 0 |
| Memory B cell | 8.740292 | 90.03554 | 1.224167 |
| Migratory dendritic cell | 0 | 100 | 0 |
| NKT cell | 0.050027 | 89.11917 | 10.8308 |

| | | | |
|---|---|---|---|
| Naive B cell | 1.806286 | 92.86433 | 5.329382 |
| Non-classical monocyte | 32.11034 | 67.49558 | 0.394075 |
| Plasma cell | 72.10482 | 11.15039 | 16.74479 |
| Plasmablast cell | 100 | 0 | 0 |
| Plasmacytoid dendritic cell | 0.106383 | 99.89362 | 0 |
| Promonocyte | 0.175747 | 94.55185 | 5.272408 |
| Suprabasal cell | 0.136033 | 77.74666 | 22.11731 |
| T helper cell | 100 | 0 | 0 |
| Treg cell | 100 | 0 | 0 |
| Type I alveolar cell | 78.05963 | 18.07975 | 3.860615 |
| Type II alveolar cell | 58.22546 | 35.42321 | 6.35133 |

**Table S7. Statistical of comparing the re-annotation of the uniLUNG core with the original annotation at Level 4.**

| Cell type | Match_labelled | Cover_labelled | Unmatch_labelled |
|---|---|---|---|
| Memory CD4 T cell | 0.135809 | 99.53773 | 0.326465 |
| Memory CD8 T cell | 0.136169 | 99.61027 | 0.253557 |
| Naive CD4 T cell | 2.529369 | 95.49559 | 1.975037 |
| Naive CD8 T cell | 0.205595 | 99.18367 | 0.610739 |
| Th1 cell | 100 | 0 | 0 |

**Table S8. Details of datasets in uniLUNG.**

| Sub-atlas | Donor status | Reference | Data accession or DOI |
|---|---|---|---|
| Cancer | LCC | Sikkema L et al. 2023. Nat Med. 10.1038/s41591-023-02327-2 | HLCA |
| Cancer | LUAD | Li M et al. 2022. Nucleic Acids Res. 10.1093/nar/gkab1020 | DISCO |
| Cancer | LUAD | Sikkema L et al. 2023. Nat Med. 10.1038/s41591-023-02327-2 | HLCA |
| Cancer | LUAD | Zhu J et al. 2022. Exp Mol Med. 10.1038/s12276-022-00896-9 | GSE189357 |
| Cancer | LUAD | Chan JM et al. 2021. Cancer Cell. 10.1016/j.ccell.2021.09.008 | HTAN |
| Cancer | LUAD | Salcher S et al. 2022. Cancer Cell. 10.1016/j.ccell.2022.10.008 | LuCA |
| Cancer | LUAD | Prazanowska KH, Lim SB. 2023. Sci Data. 10.1038/s41597-023-02074-6 | 10.6084/m9.figshare.c.6222221.v3 |
| Cancer | LUAD | Zhang L et al. 2022. Signal Transduct Target Ther. 10.1038/s41392-021-00824-9 | http://lungcancer.chenlulab.com/ |
| Cancer | LUSC | Sikkema L et al. 2023. Nat Med. 10.1038/s41591-023-02327-2 | HLCA |
| Cancer | LUSC | Salcher S et al. 2022. Cancer Cell. 10.1016/j.ccell.2022.10.008 | LuCA |
| Cancer | LUSC | Prazanowska KH, Lim SB. 2023. Sci Data. 10.1038/s41597-023-02074-6 | 10.6084/m9.figshare.c.6222221.v3 |

| | | | |
|---|---|---|---|
| Cancer | LUSC | Zhang L et al. 2022. Signal Transduct Target Ther. 10.1038/s41392-021-00824-9 | http://lungcancer.chenlulab.com/ |
| Cancer | SCLC | Chan JM et al. 2021. Cancer Cell. 10.1016/j.ccell.2021.09.008 | HTAN |
| Cancer | NSCLC | Caushi JX et al. 2021. Nature. 10.1038/s41586-021-03752-4 | GSE176021 |
| COVID-19 | COVID-19 | Li M et al. 2022. Nucleic Acids Res. 10.1093/nar/gkab1020 | DISCO |
| COVID-19 | COVID-19 | Liao M et al. 2020. Nat Med. 10.1038/s41591-020-0901-9 | GSE145926 |
| COVID-19 | COVID-19 | Lee JS et al. 2020. Sci Immunol. 10.1126/sciimmunol.abd1554 | GSE149689 |
| COVID-19 | COVID-19 | Xu G et al. 2020. Clin Transl Med. 10.1002/ctm2.224 | GSE149878 |
| COVID-19 | COVID-19 | Wilk AJ et al.2020. Nat Med. 10.1038/s41591-020-0944-y | GSE150728 |
| COVID-19 | COVID-19 | Yao C et al. 2021. Cell Rep. 10.1016/j.celrep.2020.108590 | GSE154567 |
| COVID-19 | COVID-19 | Grant RA et al. 2021. Nature. 10.1038/s41586-020-03148-w | GSE155249 |
| COVID-19 | COVID-19 | Ren X et al. 2021. Cell. 10.1016/j.cell.2021.01.053 | GSE158055 |
| COVID-19 | COVID-19 | Wang A et al. 2020. Elife. 10.7554/eLife.62522 | GSE161382 |
| COVID-19 | COVID-19 | Bacher P et al. 2020. Immunity. 10.1016/j.immuni.2020.11.016 | GSE162086 |
| COVID-19 | COVID-19 | Heming M et al. 2021. Immunity. 10.1016/j.immuni.2020.12.011 | GSE163005 |
| COVID-19 | COVID-19 | Combes AJ et al. 2021. Nature. 10.1038/s41586-021-03234-7 | GSE163668 |

| | | | |
|---|---|---|---|
| COVID-19 | COVID-19 | Melms JC et al. 2021. Nature. 10.1038/s41586-021-03569-1 | GSE171524 |
| COVID-19 | COVID-19 | Georg P et al. 2022. Cell. 10.1016/j.cell.2021.12.040 | GSE175450 |
| COVID-19 | COVID-19 | Choi B et al. 2022. Exp Mol Med. 10.1038/s12276-022-00866-1 | GSE182123 |
| COVID-19 | COVID-19 | Khoo WH et al. 2023. Clin Immunol. 10.1016/j.clim.2022.109209 | GSE196456 |
| COVID-19 | COVID-19 | Lee HK et al. 2022. iScience. 10.1016/j.isci.2022.104473 | GSE201535 |
| COVID-19 | COVID-19 | Iwamura C et al. 2022. Proc Natl Acad Sci U S A. 10.1073/pnas.2203437119 | GSE208337 |
| COVID-19 | COVID-19 | Santer DM et al. 2022. Nat Commun. 10.1038/s41467-022-34709-4 | GSE215814 |
| COVID-19 | COVID-19 | Xu J et al. 2022. Front Immunol. 10.3389/fimmu.2022.970287 | GSE216020 |
| Dev_aging | Dev_aging | Cao J et al. 2020. Science. 10.1126/science.aba7721 | GSE156793 |
| Dev_aging | Dev_aging | Zepp JA et al. 2021. Science. 10.1126/science.abc3172 | GSE149563 |
| Dev_aging | Dev_aging | Sountoulidis A et al. 2023. Nat Cell Biol. 10.1038/s41556-022-01064-x | GSE215895 |
| Dev_aging | Dev_aging | He P et al. 2022. Cell. 10.1016/j.cell.2022.11.005 | E-MTAB-11278 |
| Dev_aging | Dev_aging | Miller AJ et al. 2020. Dev Cell. 10.1016/j.devcel.2020.01.033 | E-MTAB-8221 |
| Dev_aging | Dev_aging | Ren X et al. 2021. Cell. 10.1016/j.cell.2021.01.053 | GSE158055 |
| Dev_aging | Dev_aging | Adams TS et al. 2020. Sci Adv. 10.1126/sciadv.aba1983 | GSE136831 |
| Dev_aging | Dev_aging | DePianto DJ et al. 2021. JCI Insight. 10.1172/jci.insight.143626 | GSE159354 |

| | | | |
|---|---|---|---|
| Dev_aging | Dev_aging | Madissoon E et al. 2019. Genome Biol. 10.1186/s13059-019-1906-x | PRJEB31843 |
| Dev_aging | Dev_aging | Tian Y et al. 2022. Signal Transduct Target Ther. 10.1038/s41392-022-01150-4 | PRJCA006026 |
| Dev_aging | Dev_aging | Travaglini KJ et al. 2020. Nature. 10.1038/s41586-020-2922-4 | EGAS00001004344 |
| Dev_aging | Dev_aging | Huang Q et al. 2022. Respir Res. 10.1186/s12931-022-02293-2 | GSE171541 |
| ILD | ILD | Reyfman PA et al. 2019. Am J Respir Crit Care Med. 10.1164/rccm.201712-2410OC | GSE122960 |
| ILD | ILD | Gao X et al. 2020. Cell Rep Med. 10.1016/j.xcrm.2020.100140IF: 14.3 Q1 | GSE159354 |
| ILD | ILD | Zhu L et al. 2022. Front Immunol. 10.3389/fimmu.2022.804034 | GSE190510 |
| ILD | ILD | Natri HM et al. 2023. Preprint. bioRxiv. 10.1101/2023.03.17.533161 | GSE227136 |
| ILD | ILD | Sikkema L et al. 2023. Nat Med. 10.1038/s41591-023-02327-2 | HLCA |
| ILD | IPAF | Natri HM et al. 2023. Preprint. bioRxiv. 10.1101/2023.03.17.533161 | GSE227136 |
| ILD | IPF | Li M et al. 2022. Nucleic Acids Res. 10.1093/nar/gkab1020 | DISCO |
| ILD | IPF | Reyfman PA et al. 2019. Am J Respir Crit Care Med. 10.1164/rccm.201712-2410OC | GSE122960 |
| ILD | IPF | Tsukui T et al. 2020. Nat Commun. 10.1038/s41467-020-15647-5 | GSE132771 |
| ILD | IPF | Adams TS et al. 2020. Sci Adv. 10.1126/sciadv.aba1983 | GSE136831 |
| ILD | IPF | Carraro G et al. 2020. Am J Respir Crit Care Med. 10.1164/rccm.201904-0792OC | GSE143706 |
| ILD | IPF | Liang J et al. 2022. J Clin Invest. 10.1172/JCI157338 | GSE157997 |

| | | | |
|---|---|---|---|
| ILD | IPF | Gao X et al. 2020. Cell Rep Med. 10.1016/j.xcrm.2020.100140IF: 14.3 Q1 | GSE159354 |
| ILD | IPF | de Rooij LPMH et al. 2023. Cardiovasc Res. 10.1093/cvr/cvac139 | GSE159585 |
| ILD | IPF | Natri HM et al. 2023. Preprint. bioRxiv. 10.1101/2023.03.17.533161 | GSE227136 |
| ILD | NSIP | Natri HM et al. 2023. Preprint. bioRxiv. 10.1101/2023.03.17.533161 | GSE227136 |
| ILD | PF | Habermann AC et al. 2020. Sci Adv. 10.1126/sciadv.aba1972 | GSE135893 |
| ILD | PF | Sikkema L et al. 2023. Nat Med. 10.1038/s41591-023-02327-2 | HLCA |
| ILD | PF | Li M et al. 2022. Nucleic Acids Res. 10.1093/nar/gkab1020 | DISCO |
| Normal | Healthy | Li M et al. 2022. Nucleic Acids Res. 10.1093/nar/gkab1020 | DISCO |
| Normal | Healthy | Holloway Emily M et al. 2020. Developmental Cell. 10.1016/j.devcel.2020.07.023 | E-MTAB-8221 |
| Normal | Healthy | Domínguez Conde C et al. 2022. Science. 10.1126/science.abl5197 | E-MTAB-11536 |
| Normal | Healthy | Wei Kheng The et al. 2022. BioStudies. E-MTAB-11278 | E-MTAB-11278 |
| Normal | Healthy | Travaglini KJ et al. 2020. Nature. 10.1038/s41586-020-2922-4 | EGAS00001004344 |
| Normal | Healthy | | GSE121080 |
| Normal | Healthy | Reyfman PA et al. 2019. Am J Respir Crit Care Med. 10.1164/rccm.201712-2410OC | GSE122960 |
| Normal | Healthy | Szabo PA et al. 2019. Nat Commun. 10.1038/s41467-019-12464-3 | GSE126030 |
| Normal | Healthy | Morse C et al. 2019. Eur Respir J. 10.1183/13993003.02441-2018 | GSE128033 |

| | | | |
|---|---|---|---|
| Normal | Healthy | Vieira Braga FA et al. 2019. Nat Med. 10.1038/s41591-019-0468-5 | GSE130148 |
| Normal | Healthy | Tsukui T et al. 2020. *Nat Commun*. 10.1038/s41467-020-15647-5 | GSE132771 |
| Normal | Healthy | Raredon MSB et al. 2019. Sci Adv. 10.1126/sciadv.aaw3851 | GSE133747 |
| Normal | Healthy | Goldfarbmuren KC et al. 2020. Nat Commun. 10.1038/s41467-020-16239-z | GSE134174 |
| Normal | Healthy | Han X et al. 2020. Nature. 10.1038/s41586-020-2157-4 | GSE134355 |
| Normal | Healthy | Habermann AC et al. 2020. Sci Adv. 10.1126/sciadv.aba1972 | GSE135893 |
| Normal | Healthy | Adams TS et al. 2020. Sci Adv. 10.1126/sciadv.aba1983 | GSE136831 |
| Normal | Healthy | Jaeger B et al. 2022. Nat Commun. 10.1038/s41467-022-33193-0 | GSE141939 |
| Normal | Healthy | Deprez M et al. 2020. Am J Respir Crit Care Med. 10.1164/rccm.201911-2199OC | GSE143868 |
| Normal | Healthy | Liao M et al. 2020. Nat Med. 10.1038/s41591-020-0901-9 | GSE145926 |
| Normal | Healthy | Ziegler CGK et al. 2020. Cell. 10.1016/j.cell.2020.04.035 | GSE148829 |
| Normal | Healthy | de Rooij LPMH et al. 2023. Cardiovasc Res. 10.1093/cvr/cvac139 | GSE159585 |
| Normal | Healthy | Okuda K et al. 2021. Am J Respir Crit Care Med. 10.1164/rccm.202008-3198OC | GSE160664 |
| Normal | Healthy | Okuda K et al. 2021. Am J Respir Crit Care Med. 10.1164/rccm.202008-3198OC | GSE160673 |
| Normal | Healthy | | GSE160794 |
| Normal | Healthy | Wang A et al. 2020. Elife. 10.7554/eLife.62522 | GSE161382 |

| | | | |
|---|---|---|---|
| Normal | Healthy | Schupp JC et al. 2021. Circulation. 10.1161/CIRCULATIONAHA.120.052318 | GSE164829 |
| Normal | Healthy | Pisu D et al. 2021. J Exp Med. 10.1084/jem.20210615 | GSE167232 |
| Normal | Healthy | Watanabe N et al. 2022. Am J Respir Cell Mol Biol. 10.1165/rcmb.2021-0555OC | GSE173896 |
| Normal | Healthy | Li X et al. 2022. Life Sci Alliance. 10.26508/lsa.202201458 | GSE193782 |
| Normal | Healthy | Wang C et al. 2023. Immunity. 10.1016/j.immuni.2023.01.032 | GSE196638 |
| Normal | Healthy | Natri HM et al. 2023. Preprint. bioRxiv. 10.1101/2023.03.17.533161 | GSE227136 |
| Normal | Healthy | Eraslan G et al. 2022. Science. 10.1126/science.abl4290 | GTEx V9 |
| Normal | Healthy | Sikkema L et al. 2023. Nat Med. 10.1038/s41591-023-02327-2 | HLCA |
| Normal | Healthy | Chan JM et al. 2021. Cancer Cell. 10.1016/j.ccell.2021.09.008 | HTAN |
| Normal | Healthy | Pan L et al. 2023. Nucleic Acids Res. 10.1093/nar/gkac791 | HTCA |
| Normal | Healthy | Salcher S et al. 2022. Cancer Cell. 10.1016/j.ccell.2022.10.008 | LuCA |
| Normal | Healthy | HuBMAP Consortium. 2019. Nature. 10.1038/s41586-019-1629-x | HuBMAP |
| Normal | Healthy | Lukassen et al. 2020. The EMBO Journal. 10.15252/embj.20105114 | 10.6084/m9.figshare.11981034.v1 |
| Other | Asthma | Li M et al. 2022. Nucleic Acids Res. 10.1093/nar/gkab1020 | DISCO |
| Other | CF | Sikkema L et al. 2023. Nat Med. 10.1038/s41591-023-02327-2 | HLCA |

| | | | |
|---|---|---|---|
| Other | CF | Schupp JC et al. 2020. Am J Respir Crit Care Med. 10.1164/rccm.202004-0991OC | GSE145360 |
| Other | CF | Carraro G et al. 2021. Nat Med. 10.1038/s41591-021-01332-7 | GSE150674 |
| Other | CF | Li X et al. 2022. Life Sci Alliance. 10.26508/lsa.202201458 | GSE193782 |
| Other | COPD | Adams TS et al. 2020. Sci Adv. 10.1126/sciadv.aba1983 | GSE136831 |
| Other | COPD | Watanabe N et al. 2022. Am J Respir Cell Mol Biol. 10.1165/rcmb.2021-0555OC | GSE173896 |
| Other | COPD | Salcher S et al. 2022 Cancer Cell. 10.1016/j.ccell.2022.10.008 | LuCA |
| Other | COPD | Sikkema L et al. 2023. Nat Med. 10.1038/s41591-023-02327-2 | HLCA |
| Other | CWP | Natri HM et al. 2023. Preprint. bioRxiv. 10.1101/2023.03.17.533161 | GSE227136 |
| Other | Pneumonia | Sikkema L et al. 2023. Nat Med. 10.1038/s41591-023-02327-2 | HLCA |
| Other | Sarcoidosis | Natri HM et al. 2023. Preprint. bioRxiv. 10.1101/2023.03.17.533161 | GSE227136 |

**Table S9. Cell count statistics in lung multi-disease analysis.**

| Lung status | Cells |
|---|---|
| Asthma | 7493 |
| CF | 20867 |
| COPD | 99366 |
| COVID-19 | 132758 |
| CWP | 33185 |
| Healthy | 329197 |
| ILD | 57500 |
| IPAF | 19314 |
| IPF | 127082 |
| LCC | 18123 |
| LUAD | 92700 |
| LUSC | 82651 |
| NSCLC | 124737 |
| NSIP | 26318 |

| | |
|---|---|
| PF | 350289 |
| Pneumonia | 24277 |
| SCLC | 63916 |
| Sarcoidosis | 33612 |

**Table S10. Donor information for lung cancer case studies.**

| Cancer type | Donor number | I | II | III | III/IV | IV | NA |
|---|---|---|---|---|---|---|---|
| LUAD | 290 | 112 | 22 | 26 | 37 | 58 | 35 |
| LUSC | 89 | 16 | 10 | 8 | 39 | 4 | 3 |
| SCLC | 21 | — | — | — | — | — | 21 |

**Table S11. List of LRIs in LUAD and LUSC samples.**

Due to the large size of the file, please see the supplementary Excel file on the uniLUNG website: https://lung.unifiedcellatlas.org/download/Supplementary_Table_11.xlsx

**Table S12. List of active spatial signal pathways in LUAD samples based on COMMOT.**

Due to the large size of the file, please see the supplementary Excel file on the uniLUNG website: https://lung.unifiedcellatlas.org/download/Supplementary_Table_12.xlsx

**Table S13. List of active spatial signal pathways in LUSC samples based on COMMOT.**

Due to the large size of the file, please see the supplementary Excel file on the uniLUNG website: https://lung.unifiedcellatlas.org/download/Supplementary_Table_13.xlsx

**Table S14. Total 43878 gene list.**

Due to the large size of the file, please see the supplementary Excel file on the uniLUNG website: https://lung.unifiedcellatlas.org/download/Supplementary_Table_14.xlsx
